# ProPER: Ligation-free in situ proximity extension enables multiplexed molecular-state analysis

**DOI:** 10.64898/2026.04.24.718482

**Authors:** Jorge Trojanowski, Jennifer Zilke, Valeriy Pak, Eva-Maria Geissen, Renata M. Chalupska, Robert Reinhardt, Sama AlMehallawi, Jocelyn Y. Kishi, Jennifer Heck, Kristina Haase, Sinem K. Saka

**Affiliations:** Genome Biology Unit, European Molecular Biology Laboratory (EMBL) Heidelberg, Germany; Tissue biology and disease modelling Unit, EMBL Barcelona, Spain; Faculty of Biosciences, Collaboration for Joint PhD Degree between EMBL and Heidelberg University, Germany; Data Science Centre, EMBL, Heidelberg, Germany; Digital Biology, Inc., MA, USA

## Abstract

Molecular proximity reports molecular interactions, local environments, and functional states in intact cells, but current in situ proximity detection methods remain limited in sensitivity, spatial resolution, multiplexing, and quantitative performance. Here we introduce Proximity Primer Exchange Reaction (ProPER), a spatially resolved ligation-free proximity detection strategy for multiplexed molecular-state analysis in situ. In ProPER, spatial proximity directly triggers DNA barcode extension and controlled signal amplification, autonomously converting molecular proximity into compact linear DNA concatemers within a single polymerase-driven reaction. ProPER integrates directly with established in situ assays, including immunofluorescence, FISH, and metabolic labeling, and enables high-resolution visualization of diverse molecular proximities, including RNA-RNA and protein-protein interactions, even in densely labelled environments. We show that ProPER enables sensitive and quantitative proximity measurements, supports pairwise and multivalent detection schemes, and allows intramolecular multiplexing to simultaneously measure multiple proximity-defined states on the same molecular target. We apply this capability to simultaneously resolve transcriptional, splicing, and translational RNA life-cycle states by fluorescence imaging with single-molecule-level quantification. These multiplexed molecular-state measurements enabled quantitative kinetic modeling of inflammatory gene induction directly from spatial imaging data. Specifically, we show that the differential induction delay between the TNF-α response genes TNFAIP3 and ICAM1 arises from regulation distributed across multiple expression steps, including transcript production, splicing kinetics, feedback regulation, and transcript stability, rather than a single rate-limiting bottleneck. Furthermore, our spatial data distinguished between transcription-site-associated and dispersed nucleoplasmic splicing, revealing their differential kinetics. By converting spatial proximity into programmable DNA readouts, ProPER enables multiplexed molecular-state measurements that extend spatial biology beyond molecular abundance and localization, while providing a direct path toward integration with existing spatial omics workflows.

## Introduction

The rise of single-cell RNA sequencing and, more recently, spatial transcriptomics has enabled cellular states to be defined primarily by mRNA abundance and spatial localization. However, many cellular processes are regulated not only by the amount and position of individual molecules, but also by their molecular neighbourhoods, interaction partners and functional states. These proximity-defined states can report whether a molecule is being synthesized, processed, modified, transported, localized or translated, and can therefore provide information that is not accessible from abundance or localization measurements alone. For example, nascent RNA production may begin hours before a peak cellular response is achieved at the level of mature protein. Gene expression delays are a fundamental feature of regulatory modules that ensure timely and coordinated responses to stimuli^1^, and can also drive cell-autonomous oscillations^2^. Determining which step in gene expression, such as RNA transcription, processing, transport, localization, modification, or translation, is rate limiting can help reveal the molecular mechanisms that govern these delays.

Methods that detect molecular proximity in situ offer a route to measuring these functional states directly in intact cells. Metabolic labelling or run-on sequencing (GRO-seq^3^) can resolve nascent RNAs, whereas proximity detection methods have the potential to annotate a broader range of molecular states defined by interactions or spatial localization. Although proximity alone does not necessarily indicate direct binding, it is widely used to infer molecular interactions and submicrometer spatial context^4^. Existing proximity detection methods can be broadly divided into four categories with distinct capabilities for multiplexing, multimodal detection and spatial readout: (i) Crosslinking-based methods that exploit short-range chemical crosslinkers to covalently capture molecular neighborhoods for downstream mass spectrometry or next-generation sequencing (NGS) readouts; (ii) Enzymatic proximity labeling techniques, such as BioID^5^, APEX^6^, which are often used in live samples but are typically not readily multiplexed; (iii) Fluorescence resonance energy transfer-based methods, which are preferred for dynamic detection of proximity, but are hard to multiplex and require very close proximity interactions, typically 1-10 nm, limiting their versatility, especially when probes such as antibodies are used; and (iv) DNA-based methods, which which exploit conditional interactions between oligonucleotides attached to neighbouring molecular probes.

Among DNA-based approaches, the most widely used method is the in situ proximity ligation assay (PLA or isPLA)^7–9^, in which dual affinity probes generate a circular DNA template through enzymatic ligation that subsequently undergoes rolling-circle amplification (RCA)^10^. Variants that use connector probes or splint designs with barcoded padlock probes further support in situ multiplexed proximity detection and cyclic in situ sequencing^11–16^. Despite these advances, current DNA-based proximity assays exhibit several technological limitations that restrict their broader application in quantitative spatial biology. Ligation-dependent workflows reduce reaction efficiency and quantitative performance, while bulky rolling-circle products compromise spatial resolution and hinder detection in crowded molecular environments. Furthermore, existing architectures largely treat individual proximity events as independent measurements, limiting simultaneous interrogation of multiple proximity relationships on the same molecular target. Ligation-dependent encoding of proximities without in situ amplification has been used to generate NGS readouts for higher parallelization^17^ and was recently combined with capture-based spatial transcriptomics assays^18^. Passive ligation-free approaches based on DNA co-hybridization, including proximity-dependent Hybridization Chain Reaction (HCR)^19,20^ or direct detection by DNA-PAINT for super-resolution imaging^21^, avoid enzymatic ligation but primarily rely on conditional hybridization rather than active information recording. In contrast, polymerase-based proximity writing strategies such as such as Proximity Extension Assay (PEA)^22,23^ or auto-cyclic proximity recording (APR)^24^ actively convert molecular recognition events into new DNA sequences (“DNA writing”), yet they do not offer signal amplification and have not been implemented for spatially resolved in situ measurements.

To address these challenges, we sought to develop a ligation-free, polymerase-driven proximity writing strategy that combines efficient DNA synthesis, compact signal amplification, programmable molecular encoding and compatibility with diverse spatial molecular assays. Here we present Proximity Primer Exchange Reaction (ProPER), where spatial proximity directly triggers autonomous DNA barcode extension without ligation, and the resulting proximity barcode is converted into compact linear DNA concatemers for controlled signal amplification within a single polymerase-driven reaction. ProPER integrates directly with established in situ assays, including immunofluorescence, fluorescence in situ hybridization and metabolic labelling, while enabling efficient, multiplexed and quantitative molecular-state measurements in intact cells. We demonstrate these capabilities by simultaneously resolving multiple stages of RNA life-cycle progression within individual cells, enabling quantitative modelling of inflammatory gene induction kinetics. Together, these results establish ProPER as a broadly applicable framework for programmable molecular-state analysis through spatial proximity in intact biological samples.

### Design

The technological requirements outlined above demand a proximity detection strategy that combines efficient DNA writing, compact signal amplification, programmable molecular encoding and compatibility with diverse biological modalities. Primer Exchange Reaction (PER) is a powerful isothermal DNA engineering strategy originally developed for the autonomous synthesis of deterministic single-stranded DNA concatemers^25,26^. DNA synthesis occurs in solution from freely diffusing oligonucleotide reactants through cycles of hairpin-mediated strand displacement and branch migration. The resulting DNA concatemers can subsequently be purified or directly applied as programmable DNA reagents for a wide range of downstream applications. Most notably, PER-generated concatemers have been widely adopted as tunable signal amplifiers for spatial molecular profiling, including SABER and Immuno-SABER, for multiplexed imaging of RNA and proteins in cells and tissues^27–30^. The modularity and sequence orthogonality of PER-generated concatemers have also been instrumental for highly multiplexed detection with mass cytometry and X-ray fluorescence^31,32^. Beyond imaging, PER has evolved into a versatile platform for programmable molecular sensing and information processing, capable of coupling diverse molecular recognition events to customizable DNA outputs^33^. Together, these properties make PER an ideal foundation for creating a programmable proximity-writing system.

In these earlier applications, PER functions as a solution-phase DNA synthesis reaction in which freely diffusing reactants generate programmable DNA products independently of molecular targeting. We asked whether this chemistry could instead be performed directly on immobilized biological targets, such that autonomous DNA synthesis itself becomes conditional on molecular proximity within intact cells. To transition PER from an *in vitro* synthesis tool into a *in situ* spatially resolved and programmable proximity reporting technology, ProPER, we distributed the essential PER components across separate biological targets such that new DNA synthesis is initiated only when the corresponding probes are brought into spatial proximity **(Fig. 1a)**. Molecular proximity therefore directly triggers localized PER-mediated DNA synthesis, autonomously generating compact linear DNA concatemers that encode proximity information into programmable DNA products **(Supplementary Movie)**. Unlike conventional proximity assays that generate independent signals for individual proximity events, ProPER is intrinsically designed to encode pairwise and higher-order proximity relationships as programmable DNA outputs through conditional addition of barcode units to a growing DNA strand. This gives rise to a versatile framework that naturally supports multivalent detection, molecular logic operations and combinatorial molecular-state measurements directly in intact cells.

**Figure 1.**
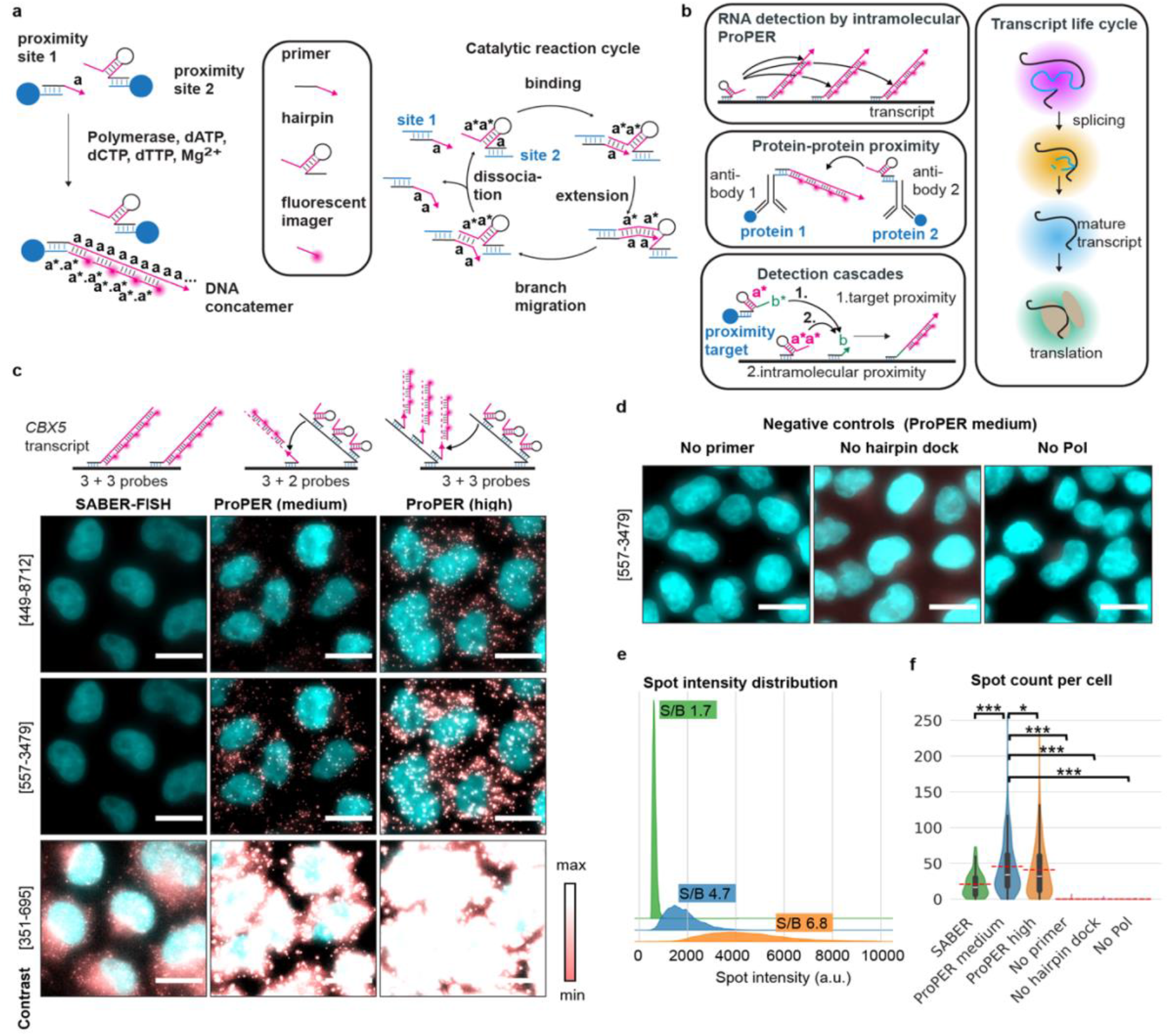
ProPER principle and in situ evaluation. **(a)** Principle of ProPER (left) and the reaction mechanism (right). The docking sites on the target immobilize the primer and hairpin. When a cognate primer and hairpin pair are in close proximity, PER^25,27,28^ takes place. The polymerase extends the primer with copies of sequence *a* and generates a linear ssDNA concatemer through autonomous and isothermal cycles of: (i) primer-hairpin binding, (ii) extension of the primer by a strand-displacing polymerase (here BST large fragment) using the hairpin’s *a\** sequence as a template, (iii) branch migration, (iv) dissociation of the primer from the hairpin. Extended DNA concatemer is detected by fluorescently labeled imager oligos. **(b)** Applications of ProPER. Primer and hairpin are docked to RNA via ISH probes or to proteins via DNA-labeled antibodies to detect proximity of the molecules or coincidence of probes on the same target. Bottom: In detection cascades different proximity barcodes can be concatenated in a programmed order. Right: The life cycle of an RNA can be revealed by multiplexed detection of proximity to life-cycle markers. **(c)** RNA-FISH and ProPER for *CBX5* gene in HeLa cells under limiting probe numbers (5-6 oligo probes). SABER-FISH was performed with in vitro pre-extended concatemers, whereas for ProPER primers were extended in situ after docking to a single site (ProPER medium) or multiple sites on a pre-extended primary probe (ProPER high). Representative images show max projections of FISH/ProPER signal (orange to white gradient) and DAPI (cyan). **(d)** Negative controls for the ProPER medium condition, where primer, hairpin dock or polymerase were left out. **(e)** Raw signal intensity of the spots (in arbitrary fluorescence units, a.u.) and **(f)** spot counts per cell showed higher values for ProPER compared to SABER-FISH, when small number of ISH probes are used. Signal to background ratio (S/B) is noted above each curve (see **Methods**). Mann-Whitney-Wilcoxon test, two-sided, * p<0.05, *** p<0.001 (exact p-values in **Supplementary Table S3**). N=105-782 cells per condition. Scale bars 10 μm.

To enable quantitative, high-resolution cell and systems biology applications, ProPER operates as a one-pot ligation-free reaction that autonomously grows compact linear DNA concatemers. This linear architecture preserves single-molecule spatial resolution while minimizing steric hindrance in crowded molecular environments. To maintain modularity across diverse biological modalities, we employ short DNA docking sequences to immobilize ProPER primers and hairpins onto in situ hybridization probes or DNA-barcoded antibodies **(Fig. 1b, left)**. Optionally, when secondary readouts or signal tuning are required, branched DNA architectures carrying multiple docking sites can be incorporated without altering the core reaction chemistry. Crucially, the high catalytic efficiency and compact spatial footprint of ProPER permit multiple orthogonal primer–hairpin reactions to proceed simultaneously on a single molecular target, enabling true intramolecular multiplexing and simultaneous readout of multiple proximity-defined molecular states. We utilize these capabilities to simultaneously interrogate spatial proximity of individual molecules to multiple functional markers within the same cell and quantitatively map the life cycle stage of RNA targets in situ **(Fig. 1b, right).**

## Results

The conceptual transition from solution-phase PER to localized proximity-dependent DNA synthesis in ProPER fundamentally changes the reaction environment. Whereas conventional PER operates with freely diffusing reactants supplied in excess, ProPER performs DNA synthesis between immobilized molecular probes whose local concentrations, stoichiometry and encounter frequencies are dictated by biological target organization. Consequently, successful implementation required substantial engineering of both the reaction architecture and experimental workflow. Here we first refine and evaluate the resulting platform using multiple complementary test systems that allow us measure and benchmark key performance metrics including background, reaction efficiency, resolution, performance in crowded cellular environments. We then we establish compatibility with multiplexing and single-molecule level multiplexed molecular state detection for quantitative analysis by applying ProPER to resolve RNA life-cycle progression in situ.

### Establishing ProPER for in situ proximity detection and signal amplification

For single-molecule FISH (smFISH), typically dozens of probes are tiled on target RNAs to achieve robust detection. As probe numbers per transcript decrease, specific detection of target transcripts gets increasingly compromised. In such a case, coincidence detection, which requires two independent detection events for the signal to be generated, can boost specificity. We used this as a test case for the capacity of PER to detect the proximity of hairpin and primer probes hybridized in situ to neighboring sites on the same transcript (intramolecular ProPER) via appended docking sequences. We targeted the *CBX5* transcript in HeLa cells with five to six primary probes that were either pre-extended in vitro by PER and directly visualized by imagers (single-molecule FISH with signal amplification, SABER-FISH), extended by ProPER in situ (ProPER medium) or served as pre-extended amplified docking site for ProPER primers and hairpins (ProPER high) (**Fig. 1c**). With these low probe numbers, signal-to-background (S/B) for smFISH was low despite additional amplification with SABER and was greatly improved in a tunable fashion with ProPER medium and high (**Fig. 1c** and **Fig. S1a).**

Importantly, in the absence of the primer oligo, the hairpin docking site or polymerase ProPER did not produce detectable signal (**Fig. 1d**). On average, ProPER yielded up to >10-fold higher signal and 4-fold higher S/B (**Fig. 1e** and **Fig. S1a**). As expected, this increased sensitivity boosted the number of detected spots per cell and average spot numbers were stable for both schemes of ProPER amplification, showing the signal is tunable and specific without generating artificial spots with increased signal amplification (**Fig. 1f**). We also tested commercially accessible chemically synthesized single-stranded DNA oligos as branches that carry the docking sites for primer and hairpin oligos (replacing in vitro pre-extension of primary probes). The defined length, e.g. ∼200 nucleotides (nt), of these synthetic branches allows controlling the number of primer and hairpin docking sites more precisely and provides a limited and reproducible reach distance within which the hairpin can bind and extend its cognate primers. Depending on the application, the branch length can be tuned to adjust the reach distance. Moreover, branches can bind to both the 5’- and 3’-end of primary probes increasing the number of available docking sites per probe and making the system modular. Performing a ProPER experiment with this setup comprises four steps: (1) hybridization of the primary probe, (2) hybridization of branches, hairpins and primers, (3) in situ PER and (4) imager hybridization (**Fig. S1b**). We tested this setup on the *ICAM1* transcript, which provides a model system for an inducible gene by TNF-α in normal human lung fibroblasts (NHLF). A larger set of primary probes maximized the number of docking sites for subsequent multiplex experiments. We found that a successful proximity reaction was dependent on TNF-α-induction of cells and primary probes, hairpin, primer, their respective docking branches and polymerase (**Fig. S1c-d**). While background levels were generally very low, we noted that the highest background was present in the no-primer-dock control, which we used as the strictest negative control for most subsequent experiments.

### Benchmarking ProPER for detection of protein-protein interactions

Next, we tested the ability of ProPER for visualization of protein-protein proximities and evaluated its detection efficiency against the current standard method PLA. We reasoned that ProPER might achieve more dense and quantitative labeling of proximity sites because it does not require an in situ ligation reaction, that has limited efficiency (e.g. due to non-circular products^13^), but only relies on extension of linear primers. It has also been observed before that the RCA is highly sensitive to reaction competition causing sparse labeling or coalescence of reaction products, particularly for dense targets^34^. Moreover, the step-by-step linear extension by a less processive polymerase in ProPER is expected to yield a more controlled and even concatemerization and lead to a more compact product compared to RCA which creates massive DNA nanoballs that often reach diameters greater than one micrometer thereby limiting resolution^34^. We chose the tubulin fiber network in NHLF cells as a defined target structure and used primary antibodies against beta-tubulin and acetylated alpha-tubulin to create the proximity detection sites. DNA barcodes on conjugated secondary antibodies were used as docking sites for primer sequences or for an adapter for hairpin docking (**Fig. 2a**). The primer and hairpin dock additionally carried imager binding sites for direct visualization of antibody binding by an adaptation of the Immuno-SABER protocol. After running ProPER we observed a dense labeling of the tubulin network (**Fig. 2b**, **Fig S2a**), whereas no ProPER signal was produced without the primary antibodies (**Fig. S2b**). Notably, a punctate signal outside the fiber network structure that was visible for the primary staining, presumably corresponding to tubulin-monomers or antibody aggregates, was absent in the ProPER signal (arrows in zoom-in, **Fig. 2b**). Accordingly, ProPER showed the highest agreement with the consensus image of primary stainings (**Fig. S2d**), measured by the pixel-wise correlation coefficient (**Fig. 2e**). In comparison, isPLA produced a specific (**Fig. 2c, S2c**), but discontinuous and punctate pattern (**Fig. 2c**), making it hard to discern the fiber structure. Detected ProPER tubule fragments (**Fig. 2d, S2f**) were significantly more elongated than the more spherical PLA structures, as measured by fragment eccentricity (**Fig. 2e**). The size of PLA punctae was larger compared to ProPER structures, which were close to the resolution limit, (**Fig. 2f**). We concluded that ProPER enables a relatively efficient labeling of proximity sites and offers better resolution owing to smaller fluorescent product sizes.

**Figure 2.**
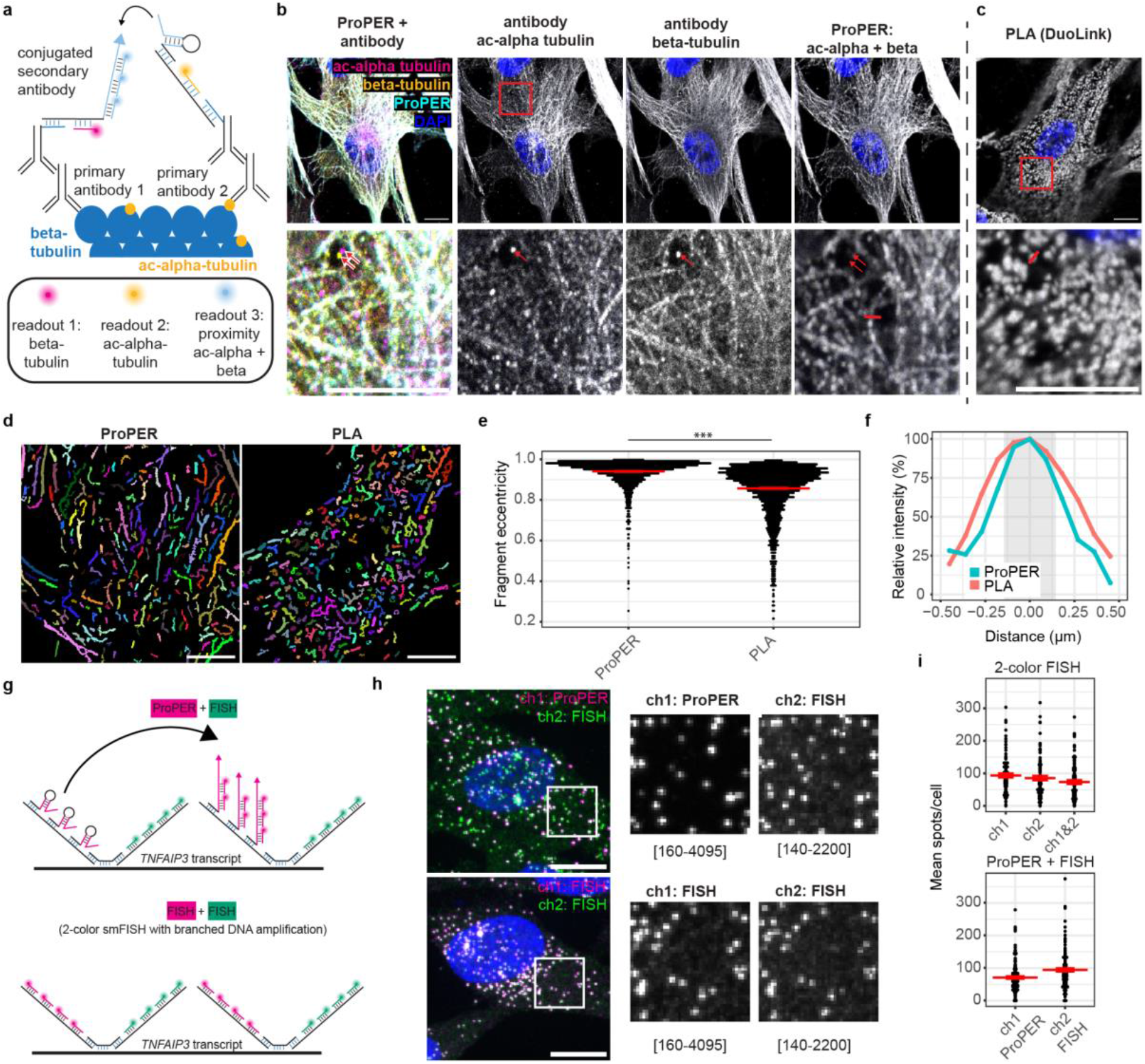
Evaluation of efficiencies in protein-protein proximity and RNA-transcript detection by ProPER. (a-e) Proximity detection of acetylated alpha-tubulin and beta-tubulin in normal human lung fibroblasts. **(a)** The detection mechanism relies on DNA conjugated secondary antibodies that bind the ac-alpha and beta-tubulin primary antibodies. They provide a binding platform for ProPER hairpin and primer, as well as for imagers for direct visualization of antibody patterns. **(b)** Multicolor merge (left) and single channel images show that the cytoskeleton morphology displays high agreement between the individual antibody channels and the ProPER proximity signal. Staining artifacts (red arrows) visible in the zoom-in region (bottom row) in individual channels did not propagate to the ProPER signal. **(c)** In comparison, proximity detection with isPLA for the same antibodies showed a more discontinuous pattern of spots. **(d)** Automatically generated segmentation masks for tubular structures for ProPER and PLA (Single-scale Frangi-algorithm, see **Methods**). Full FOV is shown in **Fig. S2f**. **I** The fragment eccentricity was calculated for the detected tubular objects, indicating more elongated structures for ProPER and more circular objects for PLA. Mann-Whitney-Wilcoxon test, two-sided, *** p=1.5e-213. N=1733-2774 fragments in 14-24 cells. **(f)** Line profiles of the representative structures shown as the signal intensity along red lines in **b-c** bottom row (normalized to the maximum intensity), showing the larger cross-section of the structures visualized by isPLA. The diffraction limit is indicated by the gray shaded area. **(g-i)** Comparison of ProPER detection efficiency to dual-color smFISH for *TNFAIP3* transcripts in NHLF cells 120 min after TNF-α exposure. **(g)** For ProPER, 40 pairs of probes having a hairpin or primer binding site alternated to be detected in channel 1 (ch1) with Alexa Fluor 647-labeled imager oligos (red). Each probe had a second arm for amplified smFISH detection in a second channel to be detected in channel 2 (ch2) with ATTO565-labeled imagers (green). For benchmarking, we performed 2-color smFISH using the same probe-branch layout and primary probe set of 80 tiling probes, where each probe presented a binding site for branch concatemers that are then detected by ATTO565 or Alexa Fluor 647 imagers. **(h)** Representative images and zoom-ins showing high co-localization between ProPER and FISH (top) or 2-color FISH (bottom). **(i)** Spots were called and counted in their individual channels (ch1/ch2 for 2-color FISH, ch-1-ProPER/ch2-FISH for ProPER). If a co-localizing spot was called in both FISH-channels within a 3-pixel radius, it was considered dual-color (ch1&2). Average spot numbers detected per cell were very similar between ProPER (ch1: 71.0, s.e.m. 4.0) and the high-confidence dual-color FISH spots (ch1&2: 73.5, s.e.m. 5.9). n=92-128 cells per condition. Red bars indicate the mean, red error bars +/- s.e.m. Scale bars 10 μm.

### Evaluation of ProPER detection efficiency

To quantify and benchmark ProPER detection efficiency, we targeted the strongly TNF-α inducible transcript, *TNFAIP3,* with signal amplified single-molecule FISH (smFISH) or ProPER. For ProPER 80 probes were assigned to 40 pairs of hairpin or primer docks **(Fig. 2g**, top). Independently, as a gold standard reference, we performed smFISH for simultaneous detection of each molecule in two fluorescence channels using the same set of 80 probes with branched DNA amplification, enabling coincidence detection that reduces false positives and yields high-confidence RNA spots (“dual-color FISH”, **Fig. 2g**, bottom). In both cases there was high visual agreement between the two readout channels (**Fig. 2h**) and the total number of spots detected per cell (**Fig. S2g**). Average number of spots per cell was similar for ProPER versus dual-color-FISH, suggesting the efficiency of proximity-based detection by ProPER is on par with dual-color FISH (**Fig. 2i**). Distance between co-localizing spots across the two channels was slightly higher in the case of ProPER versus smFISH (**Fig. S2h**), and the spot intensity variation was ∼2 times larger (**Fig. S2i**), which is expected considering potential variation in relative positioning of probe pairs under in situ conditions.

### Screening for hairpin-primer pairs to enable multiplexing

A particular advantage for in situ implementation of PER is the multiplexing potential. Previously, 50 different PER primer-hairpin pairs were designed and tested for extension efficiency and orthogonality under in vitro conditions^25,27,28^. A PER-inspired strategy using largely the same primer sequences was also used before for thermocyclic extension with linear diffusive probes, validating the simultaneous orthogonal extension of the sequences in one tube^35^.

When PER is performed in vitro, the hairpin-primer ratio can be adjusted to obtain a desired concatemer length, for example by individually optimizing the concentration of hairpins in solution to even out differences in extension efficiency of each sequence pair^28^. In contrast, when PER is performed in situ, the hairpin-primer ratio is pre-determined by the proximity sites, making it uncertain whether all hairpin-primer sequence pairs would perform well for in situ proximity detection under the same reaction conditions. We selected h./p.27, which is the most performant PER sequence pair for in vitro extension^28^, as a benchmark and chose 8 hairpin-primer pairs from the set of 50 previously validated orthogonal hairpin-primer combinations^25^ to test their performance under in situ conditions. Six out of the nine original sequences efficiently detected *ICAM1* transcripts in induced cells by intramolecular ProPER (**Fig. 3a,c**). Two sequences did not produce any signal. One of the sequences (h./p.38) was less efficient and had relatively high background counts in uninduced cells (**Fig. S3a,c**). Finally, h./p.50 pair performed well in terms of signal generation but had a higher (absolute) number of background counts (**Fig. 3a,c** and **S3a,c**).

**Figure 3.**
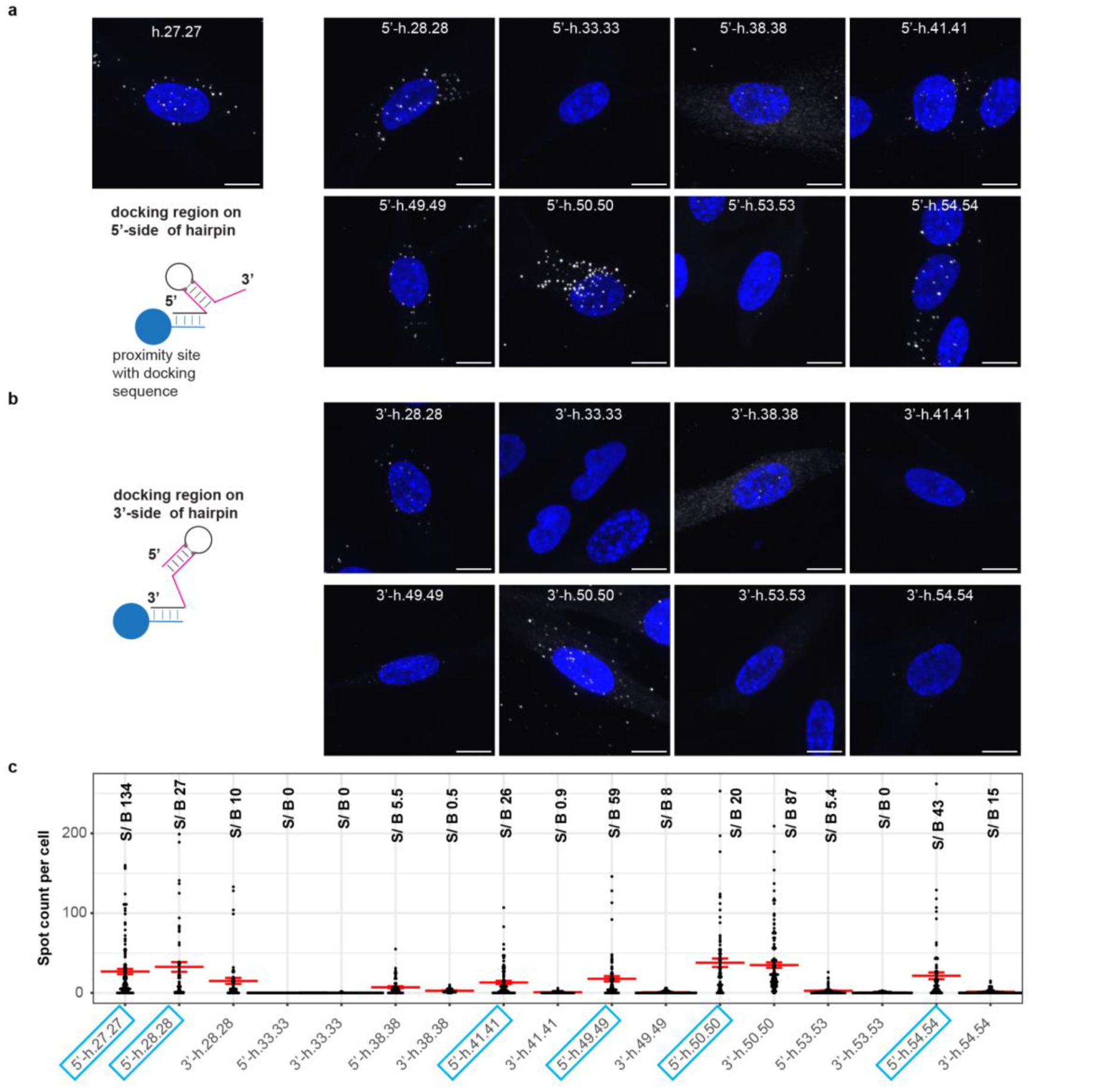
Screen for functional hairpin-primer pairs. Two designs for immobilized hairpins were tested with the docking sequence either on the 5’- or 3’-end of the catalytic hairpin. **(a-b)** Representative images of *ICAM1* transcripts in NHLF cells 120 min after TNF-α treatment with different hairpin-primer sequences with the docking sequence of the primer on the 5’- (in **a**) or 3’-end (in **b**). **(c)** Comparison of spot counts per cell for the tested hairpin-primer pairs. S/B reports the signal to background ratio corresponding to spot numbers relative to the uninduced condition. Blue boxes indicate the best performing hairpins with 5’-docking site. n=39-130 cells per condition. Red bars indicate the mean, red error bars +/- s.e.m. Scale bars 10 μm.

Performing PER in situ requires the additional design decision, whether to place the docking site of the hairpin on its 5’- (**Fig. 3a**) or 3’-end (**Fig. 3b**). When testing both orientations, we found that the hairpin design with docking site on the 5’-end performed more robustly with higher efficiencies compared to their cognate 3’-dock sequence (**Fig. 3a-c, S3**). Spot intensities were in a similar range for the well-performing sequence pairs (**Fig. S3e**). Based on these experiments, five orthogonal sequences were validated to have similar performances for in situ PER (h./p. 27, 28, 41, 49 and 54), while two additional sequences could potentially be included taking some variation into account for their signal and background levels (h./p. 38 and 50).

### Resolving RNA life-cycle stage at single molecule level with ProPER

One of the most interesting applications of proximity detection is identification of subpopulations of biomolecules based on their molecular environment or activity state. This has been recently leveraged for in situ profiling of transcribing and translating transcripts^15^. The life-cycle stage of an RNA transcript from transcription, splicing, nuclear export and translation to degradation (**Fig. 4a**) poses many levers for regulation of cellular state and response to external effects. The high detection rate and multiplexability of ProPER makes it suitable for kinetic studies of gene expression and RNA life cycle, since the flexible framework of concatemeric branches and modular hairpin-primer pairs allows simultaneous labeling of multiple life-cycle stages.

**Figure 4.**
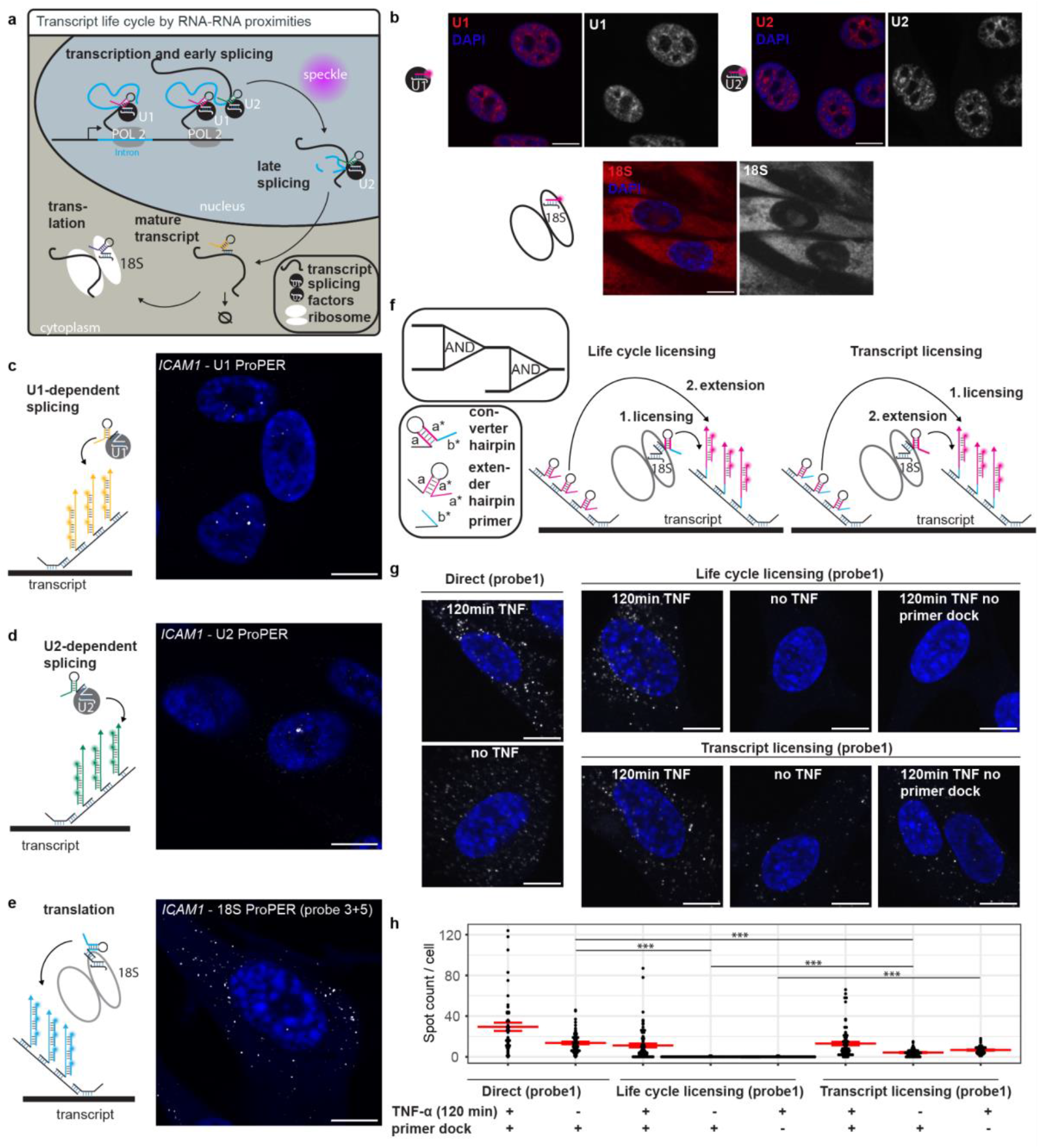
Transcript life cycle staining by ProPER and use of ProPER for in situ logic operations. **(a)** Schematic showing steps and markers of the life cycle of a transcript that can be used to visualize the transcript stage by ProPER. Life-cycle markers include small nuclear RNA splicing factors (U1, U2). Splicing can occur at the transcription site (TS) or post-transcriptionally dispersed in the nucleoplasm. After nuclear export of mature transcripts, they interact with ribosomes for translation, and we made use of proximity to the 18S RNA small ribosome subunit to mark ribosome-associated (“translating”) transcripts. All transcripts irrespective of their life-cycle stage can be simultaneously detected by ProPER coincidence detection of transcript probes (total). **(b)** Visualization of life-cycle marker RNAs (U1, U2 snRNA and 18S rRNA) by direct FISH probe binding. **(c-e)** Detection schemes and representative images for ProPER between life-cycle markers **(c)** U1, **(d)** U2, **(e)** 18S and *ICAM1* transcripts in NHLF cells 120 min after TNF-α exposure. Note that 18S life-cycle target probes 3 and 5 were used from the optimization shown in **Fig S4**. **(f)** In situ logics operations with ProPER can be utilized as a licensing strategy to suppress background when using sub-optimal life-cycle target probes (18S-probe1). Two reaction schemes were tested for increased selectivity by creating an AND-AND logic gate: For life-cycle licensing, a converter hairpin on the 18S rRNA licenses the primer on the transcript by adding a new primer sequence. Further extension by the extender hairpin (on the transcript) can then operate on the converted sequence. For transcript licensing, the converter hairpin is on the transcript and the extender on the 18S rRNA. A detailed illustration of the AND-AND gate logic is presented in **Fig. S4f**. **(g)** Representative images show reduced background levels of 18S-*ICAM1* transcript ProPER signal under life-cycle licensing (top, right) compared to direct ProPER (left) or transcript licensing (bottom, right). **(h)** Average spot numbers were counted per cell; n=47-77 cells per condition. Mann-Whitney-Wilcoxon test, two-sided, *** p<0.001 (exact p-values in **Supplementary Table S3**). Red bars indicate the mean, red error bars +/- s.e.m. Scale bars 10 μm.

We selected the splicing factors U1- and U2-snRNA as proximity markers for early RNA processing and the small subunit ribosomal RNA 18S as a life-cycle stage marker for translation. We verified that the oligo probes which would serve as docking sites for the catalytic hairpins localized to the expected cellular compartments (**Fig. 4b**). As expected, U1 and U2 were detected in the nucleus with enrichment in a punctate pattern that is in line with their nuclear speckle enrichment. 18S was mainly cytoplasmic and additionally showed signal in nucleoli where rRNAs are transcribed and assembled into ribosomes.

ProPER experiments then measured the proximity of *ICAM1* RNA to the life-cycle markers (**Fig. 4c-e**). Primers were docked onto the primary probes via the concatemeric branches, and the hairpin was docked directly on the probe targeting the life-cycle marker. U1- (**Fig. 4c**) and U2-ProPER (**Fig. 4d**) revealed nuclear transcripts in the splicing state, while 18S revealed cytoplasmic translating transcripts (**Fig. 4e**). No ProPER signal was detected when removing the primer dock or the TNF-α stimulus (**Fig. S4a-b**), confirming specificity of the reaction.

Of note, 18S rRNA constitutes a particularly challenging target for specific detection, due to its very high abundance in the cell and its strong embedding in the ribosome. Especially presentation of the hairpin probe on a very abundant structure may cause local concentration of the catalytically acting probe and increase the risk of false positives. Indeed, our first trial with a single probe on 18S resulted in a considerable level of false positives in uninduced cells (**Fig. S4c**).

To increase the detection specificity and reduce the background detections that were prominent with 18S-docking probe 1, we explored the capacity of ProPER to perform connected logic operations. The proximity detection was split into two steps (**Fig. 4d**). During the “licensing step” a converter hairpin on the ribosome attaches sequence *a* to primer *b* on the target transcript. In the extension step an extender hairpin on the target transcript attaches multiple copies of sequence *a* to the licensed primer *ba*. It thereby creates a logical AND-AND gate (**Fig. S4e**) as it requires first the proximity of the ribosome and transcript and then the proximity of the two classes of primary probes on the transcript itself. We named this approach life-cycle licensing (**Fig. 4f, left**). The second possibility to implement the detection cascade was to place the converter hairpin on the transcript and the extender hairpin on the transcript (transcript licensing, **Fig. 4f, right**). This approach, indeed, reduced false detections in the negative controls from an average of 13.7 spots per cell (using direct 18S-ProPER with probe 1) down to 0.03 or 4.2 for life-cycle and transcript licensing, respectively (**Fig. 4g-h**). The lower specificity of transcript-licensing compared to life-cycle licensing is potentially linked to the higher abundance of 18S rRNA-compared to transcript docking sites. It seems plausible that the primer conversion step has a stricter gate keeper function compared to concatemer extension and is therefore more effective on the more abundant proximity site (in this case 18S rRNA). Licensing led to a reduction of detected spots in the test condition, but this was compensated when lowering intensity thresholds without compromising specificity (**Fig. S4e**). Beyond yielding increased specificity, the licensing approach illustrates the very promising case of multivalent proximity detection with ProPER.

Alternatively, to achieve robust detection, we screened four additional primary probes on 18S rRNA (**Fig. S4f-g**) and found probes 3 and 5 to be most favorable. Probe 4 did not produce any signal, potentially due to an inaccessible binding location in the small ribosomal subunit. For 18S-proximity we settled on using two hairpin-docking probes (probes 3 and 5) on the life-cycle target which we identified as yielding highest signal vs. background, in comparable range to the licensing strategy, so for simplicity we used this approach for the subsequent experiments (**Fig. 4e** and **S4b**).

We also tested the ability of ProPER to detect nascent RNA, as an additional life-cycle target, by detection of proximity to a metabolic ethynyl-uridine (EU) label, which was provided to cells prior to fixation and was afterwards subject to click reaction with a docking oligo that hybridizes to the hairpin probe (**Fig. S5a-c**). In parallel, proximity to U1-snRNA was measured by ProPER, and total transcript abundance for TNF-α induced genes *TNFAIP3* and *ICAM1* was measured by SABER-FISH. We found that the staining patterns of EU- and U1-ProPER were very similar (**Fig. S5d**). While there was some background binding of the EU-docking click-oligo (**Fig. S5b**), this did not transmit to the ProPER signal in the negative control without EU supplementation (**Fig. S5e**). In line with the similar staining patterns for EU- and U1-associated transcripts, the majority of large (“cluster”) spots was present in both the EU- and U1-channel (**Fig. S5f**) 30 and 60 min after TNF-α induction. This indicates that U1-recruitment to the transcription site (TS) happens rapidly after the onset of transcription. We found that U1 proximity provides a reliable proxy for early stages of gene expression, obviating the need for metabolic labeling. This simplifies experimental design and enables more straightforward interpretation of induction-based time-course data.

### RNA life cycle imaging in individual cells with multiplex ProPER

Building on these feasibility tests, we aimed to visualize the full RNA life cycle from early processing by splicing to translation in each cell by establishing four-plex ProPER for total transcript and U1-, U2- and 18S-proximity. We selected four orthogonal hairpin-primer pairs (**Fig. 3c**) and designed a scheme (**Fig. 5a**) in which the target transcript carries five distinct docking sites for primers or hairpins. These are distributed on four concatemeric branches strands binding to the 5-’ and 3’-overhangs of the primary probes. One of the branches has two different types of docking sites for hairpins for intramolecular ProPER and primers for 18S proximity detection while the others are composed of repeats of the same. The two genes for which we applied this approach belong to different kinetic TNF-α response classes: *TNFAIP3* encoding A20 protein, a negative regulator of TNF-α signalling, belongs to the class of fast induced genes, while *ICAM1* encoding a surface protein has intermediate/late kinetics (**Fig. 5b**). Four-plex life cycle imaging 120 min after TNF-α stimulation showed the subcellular distribution of spots mirroring what was seen in the single-plex experiments (**Fig. 5c**). Notably, *TNFAIP3* already had high total and translating transcript numbers, whereas for *ICAM1* mainly splicing-associated spots were visible. The expression state was heterogenous between cells, especially for *ICAM1* which showed splicing spots in a small population of cells, whereas there was a wider response for *TNFAIP3* (**Fig. S6a**). Leaving out the branches carrying the primer docking sites as a negative control removed spots for all four detection modalities ( **Fig. 5d-f, Fig. S6b-c**).

**Figure 5.**
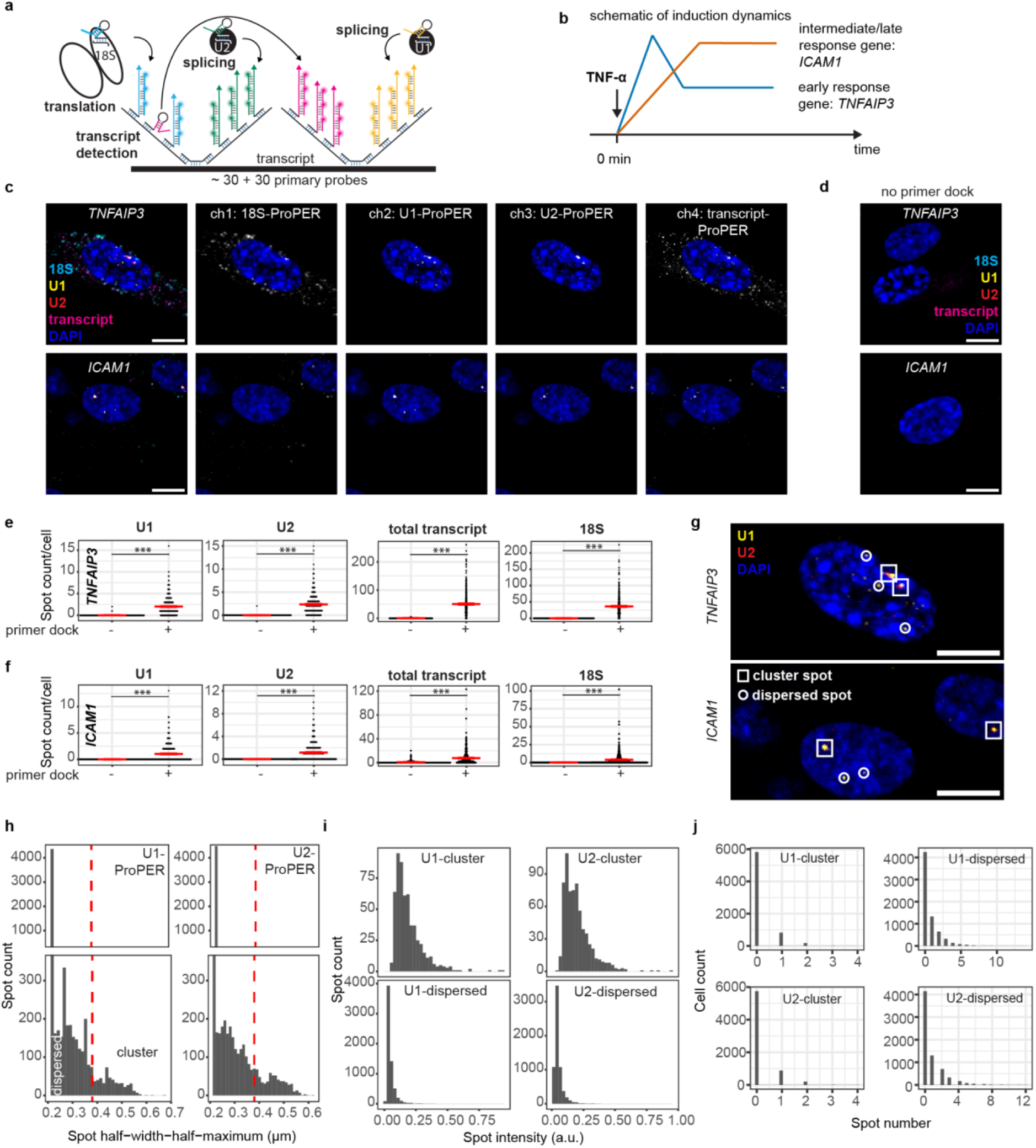
Multiplex ProPER detection. **(a)** Detection scheme using four different branches bound to the primary probes on the transcript as a binding platform for primers and the hairpin for transcript detection. Four orthogonal hairpin-primer sequence pairs were used to detect the proximity to U1, U2 and 18S, as well as the coincidence of neighboring primary probes on the transcript. Successful primer extension is visualized by cognate fluorescent imager probes for each target. **(b)** Schematic of the induction dynamics for the studied transcripts *TNFAIP3* and *ICAM1* that are from two distinct kinetic classes of TNF-α response genes: early and intermediate/late. **(c)** Representative images of fourplex ProPER for TNFAIP3 and ICAM1 transcripts 120 min after TNF-α exposure (18S: Alexa Fluor 750, cyan; U1: Alexa Fluor 647, yellow; U2: ATTO 565, red, total transcript: ATTO 488, magenta). Max. projections of three z-slices in the cell center. Images and data shown in this figure are part of the full data set presented in Fig. 6. **(d)** Negative control of 4-plex ProPER without primer docking branches. Representative multicolor images. Scale bars 10 μm. **(e-f)** Spot counts per cell in the four life-cycle channels compared to the negative control (no primer dock) at 120 min time point for **(e)** TNFAIP3 and **(f)** ICAM1. n=353-408 cells per condition pooled from three replicates. Mann-Whitney-Wilcoxon test, one-sided, *** p<0.001 (exact p-values in **Supplementary Table S3)**. **(g)** Zoom-in of nuclei in **c** showing U1- andU2 ProPER signal. Large cluster spots are annotated with a white box, small, dispersed spots with a white circle. **(h)** Distribution of spot sizes measured by the half-width-half-maximum of the spot intensity profile for U1 and U2 spots in the nucleus. Note the break in the y-axis of the distribution to ensure visibility of spot populations at the lower resolution limit (distribution peak at 0.2 μm) and the less-abundant larger spots. The red dashed line indicates the cut-off to classify spots as “dispersed” or “cluster” (manually set to separate the spot population with high HWHM). **(i)** Distribution of spot intensities for the two spot classes (dispersed/cluster). **(j)** Distribution of the number of cluster or dispersed U1/ U2 spots per nucleus. The number of cluster spots is typically between 0 and 2, compatible with the range of active TS for a diploid cell. Spots in panels **h-j** were pooled from all of the time points and both genes from the experiment in Fig. 6 from n=6838 cells. Gene-resolved spot statistics are shown in **Fig. S6d-i**.

We observed two classes of U1- and U2-associated splicing spots based on their size and intensity (**Fig. 5g**). Most splicing spots were small, but the size distribution **(Fig. 5h)** revealed a population of large spots, for which we defined a cutoff at 380 nm half-width-half maximum (HWHM) that separates these populations in the bimodal distribution. We considered spots pooled from both genes to define a shared classification criterion (**Fig. S6d-i**). The large spots had higher intensities (**Fig. 5i**) and visually displayed irregular shapes (**Fig. 5g**) that are not compatible with a fluorescence point source. We concluded that they contain more than one transcript copy and refer to them as “cluster spots” to distinguish them from the small “dispersed spots”. Nuclei typically did not contain more than two splicing cluster spots (**Fig. 5j**), which makes it seem likely that they are located at TS of the two gene alleles (also in line with the nascent RNA labeling **Fig. S5**). They could arise from co-transcriptional splicing^36^ or splicing of fully transcribed pre-mRNAs that have not yet diffused away from the TS (hence referred to as TS-bound splicing).

Besides the association of splicing with the TS, the localization relative to nuclear speckles is interesting and three classes of RNAs with constitutive, transient or no speckle-association have been previously identified^36,37^. We performed immunostaining for SON, a primary scaffold protein^38^ and marker of nuclear speckles, after ProPER for U1-/U2-bound *ICAM1* or *TNFAIP3* transcripts (**Fig. S8a**), confirming thereby that ProPER can be combined with additional stainings. Interestingly, *TNFAIP3* splicing spots had longer distances from nuclear speckles compared to *ICAM1* (**Fig. S8c**) and *ICAM1* spots were more often inside a speckle (distance of 0 μm). This difference was confirmed for dispersed spots, whereas for the relatively low number of cluster spots it was not statistically significant. Overall, this indicates that splicing of the two genes is not strictly confined to speckles, and that an association with speckles might be more prevalent for *ICAM1* than for *TNFAIP3*. These observations are compatible with a distributed mode of co- and post-transcriptional splicing rather than strict subnuclear compartmentalization of splicing activity.

In preparation for a quantitative analysis, a pipeline was set up to segment cells, and call, filter and classify spots in the four fluorescence channels (**Fig. S7**). The resulting dataset contained the coordinates of each spot, its assignment to a cell and compartment, its presence in the four readout channels and the classification as a cluster or dispersed spot (for U1 and U2). We required overlap with the total-transcript channel (intramolecular ProPER) as a filtering criterion for U1-, U2 and 18S-spots which avoids setting individual intensity thresholds for these channels and yields high-confidence spots.

### Regulation of gene induction kinetics is distributed over multiple expression steps

Next, we aimed to pinpoint which steps in the RNA life cycle are differentially regulated for *TNFAIP3* to be induced faster than *ICAM1*. Interestingly, both genes have comparable gene lengths and canonical intron numbers (TNFAIP3: 16 kbp, 8 introns; ICAM1: 15 kbp, 6 introns) and are direct targets of TNF-α signalling via NFκB^39,40^. Previously, their expression time difference has been attributed to transcript stability and splicing times based on bulk RT-PCR measurements in mouse cells^41,42^ but a comprehensive and quantitative single-cell level analysis of the life cycle progression has not been performed before. We applied life cycle labeling with multiplexed ProPER to explore the response kinetics in human cells over the course of 24 h after induction (**Fig. 6a**).

**Figure 6.**
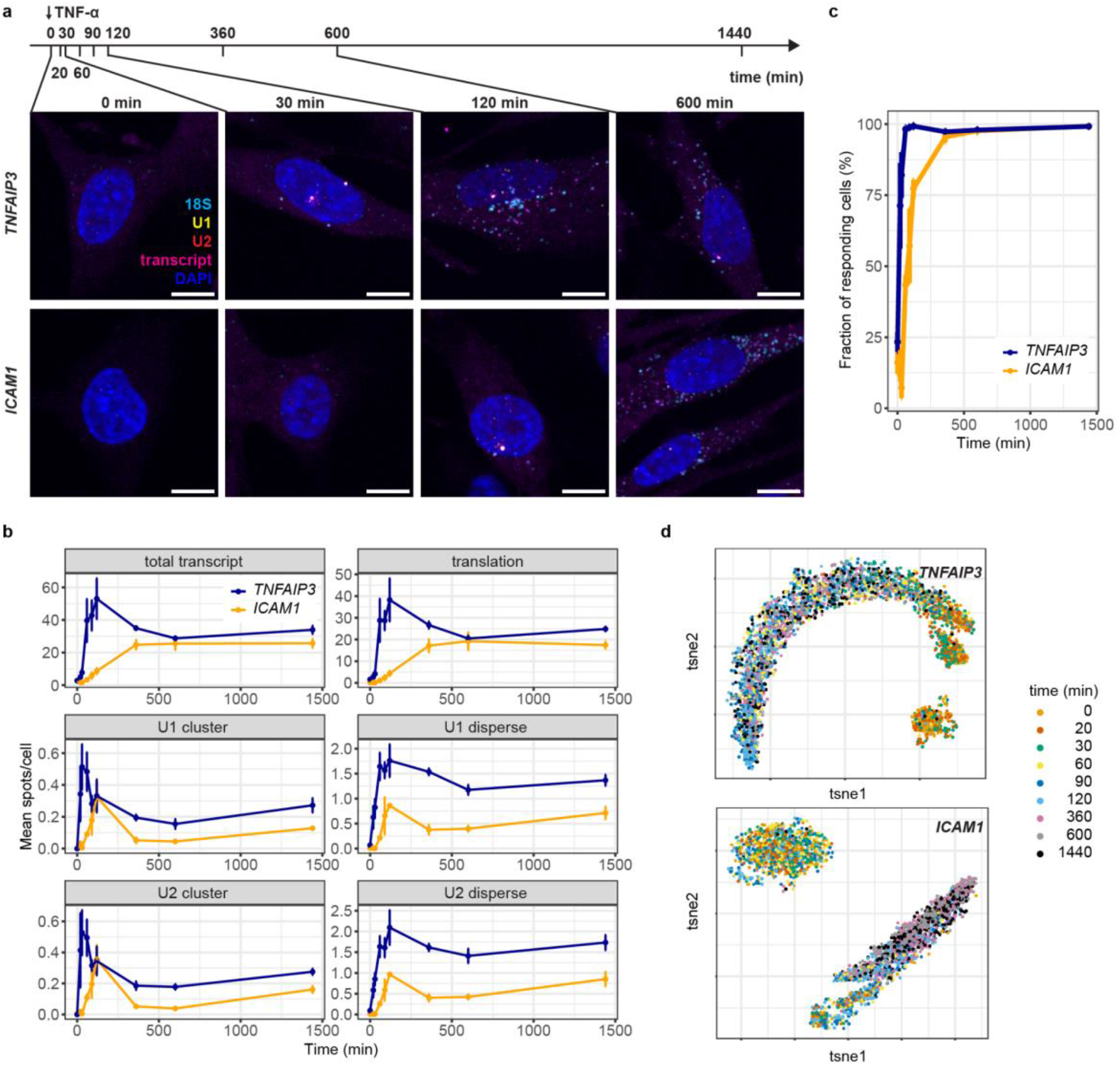
RNA life-cycle tracing of two TNF-α-induced genes. **(a)** Time points after TNF-α addition and representative multicolor images of NHLF cells after fourplex ProPER for U1-, U2-, 18S-proximity and transcript detection for *TNFAIP3* or *ICAM1*. Maximum projection of three z-planes. The complete time series is shown in **Fig. S9**. Scale bars 10 μm. **(b)** Average spot count time courses for the four ProPER readouts. U1 and U2 spots were classified as cluster or dispersed spots. Time courses for individual replicates are shown in **Fig. S10a**. 91-162 cells per time point and replicate. Error bars show +/- s.e.m. from n=3 replicates. **(c)** Fraction of responding cells over time. Responder cells were defined as having a minimum number of at least one spot type (see **Methods** for details). **(d)** t-SNE representation of single cell spot count profiles color coded by time after TNF-α addition. In addition to the spot types depicted in **b** the fraction of transcripts in the nucleus (plotted in **Fig. S10b**) was included for this representation. *TNFAIP3*: n=3386 cells, *ICAM1*: n=3452 cells.

On a qualitative level, life cycle staining with ProPER revealed the progression of the multiple stages of gene expression with expected nuclear and cytoplasmic localization steps (**Fig. 6a, Fig. S9**). For quantitative insights into the population response dynamics, we performed spot detection and filtering for each spot type (**Fig. S7**) and calculated the average spot numbers per cell and percentages of different subpopulations (**Fig. 6b, Fig. S10a-c**). At early times (20-30 min for *TNFAIP3*) up to two cluster spots per nucleus appeared in the U1- and U2-splicing channels. For ICAM1 there was an initial lag phase followed by peak TS-bound splicing at 120 min. The dynamics were nearly identical for U1 and U2 meaning that any delay between their recruitment is likely shorter than the time resolution of our experiment. For *TNFAIP3* the number of dispersed splicing spots increased with a small delay (peak at 60 min), reflecting the time needed to terminate transcription and leave the TS. TS-bound splicing (U1 and/or U2 clusters) went down between 6 and 10 h, but this was less pronounced for dispersed splicing.

The relative dynamics of total transcript numbers were comparable to the previously reported gene induction dynamics in mouse cells^41^. For *TNFAIP3* there was an overshoot at 120 min before levels dropped to an intermediate sustained plateau, whereas ICAM1 continuously approached its plateau value (around 360 min). The number of translation events (18S channel) increased in parallel. Interestingly, there was only a small delay between the rise of total transcript numbers and translation events. The fraction of translating transcripts (relative to total transcripts) reached a plateau of around 70% at 90 min for *TNFAIP3*, whereas it took 360 min for ICAM1 (**Fig. S10b**). Since the translation fraction is already high (>50%) from early time points on, the contribution of a translation delay to the expression dynamics appears to be minor.

The transcriptional response showed high variability between individual cells (**Fig. S6a**). This can be attributed to at least two processes: transcriptional bursting^43^, which accounts for the stochastic presence or absence of TS, and variable first-response time for individual cells^43,44^. Based on the presence of transcripts in any of the life-cycle stages, we, indeed, found that at early time points only a subfraction of cells responded and that for *ICAM1* the fraction of responding cells continued to rise until the 360 min time point (**Fig. 6c**). Cell populations with 1 or ≥2 splicing cluster sites emerged early for TNFAIP3 and were smaller and more transient for ICAM1 (**Fig. S10d**). We wondered if gene activation followed a single trajectory or distinct paths to activation. We defined the cell expression state based on the six life-cycle stage-relevant measures, including the distinction of U1/2 cluster and disperse, and the fraction of nuclear transcripts as a proxy for nuclear export (**Fig. S10b**). The single cell representation of the data displayed a non-responder population, enriched in cells from early timepoints, and a responder population, which coarsely reflected the temporal order (**Fig. 6d**). The responder population spanned a trajectory from low to high counts of total and translating transcripts (**Fig. S10e-f**), which corresponded to early and late times for *ICAM1*. For *TNFAIP3,* cells at late time points (600-1440 min) were enriched in the middle of the trajectory together with cells from the 60 min time point. This indicates that after the transcriptional overshoot the cells return to a steady state of transcript production and degradation. Interestingly, we found that with time cells filled the trajectory from low to high expression levels and that cells with splicing clusters and higher numbers of dispersed splicing spots were at the leading front of the trajectory (**Fig. S11**). This meant that for both genes the cells with at least one splicing cluster had higher total transcript counts for most time points **(Fig. S12 a-b)**. Even higher expression was observed if at ≥2 splicing clusters were present **(Fig. S12 c-d)**. This can best be explained if the heterogeneity between cells is mainly driven by their distributed transition to a stable responder state and less by the stochasticity of short transcriptional bursting events. Thus, the cells showed a common response pattern, but with variable onset across individual cells.

### Kinetic modeling pinpoints the critical steps for gene-specific activation

Previously in mouse cells, it has been reported that transcription and splicing can be initiated with similar dynamics across TNF-α response genes but the completion of these steps and further RNA processing can lead to very different dynamics for the mature transcript levels^42^. Our time-course data in human cells is largely consistent with these results, but we further observed additional differences in the early TS-bound splicing dynamics and a longer lag phase and delayed peak activity for *ICAM1*. To further resolve the dynamics of the expression states and the points of gene-specific regulation, we leveraged the quantitative RNA life-cycle data obtained by ProPER to construct a kinetic model (**Fig. 7a**) based on ordinary differential equations (ODEs, for a detailed description of the mathematical model and modeling results see **Supplementary Note**). The model consists of parallel steps of expression for *ICAM1* and *TNFAIP3*. The rate constants of these steps can either be shared or have gene-specific values. By systematically testing the models with either shared or gene-specific parameters it is possible to identify the expression steps that are necessary for gene specific response dynamics.

**Figure 7.**
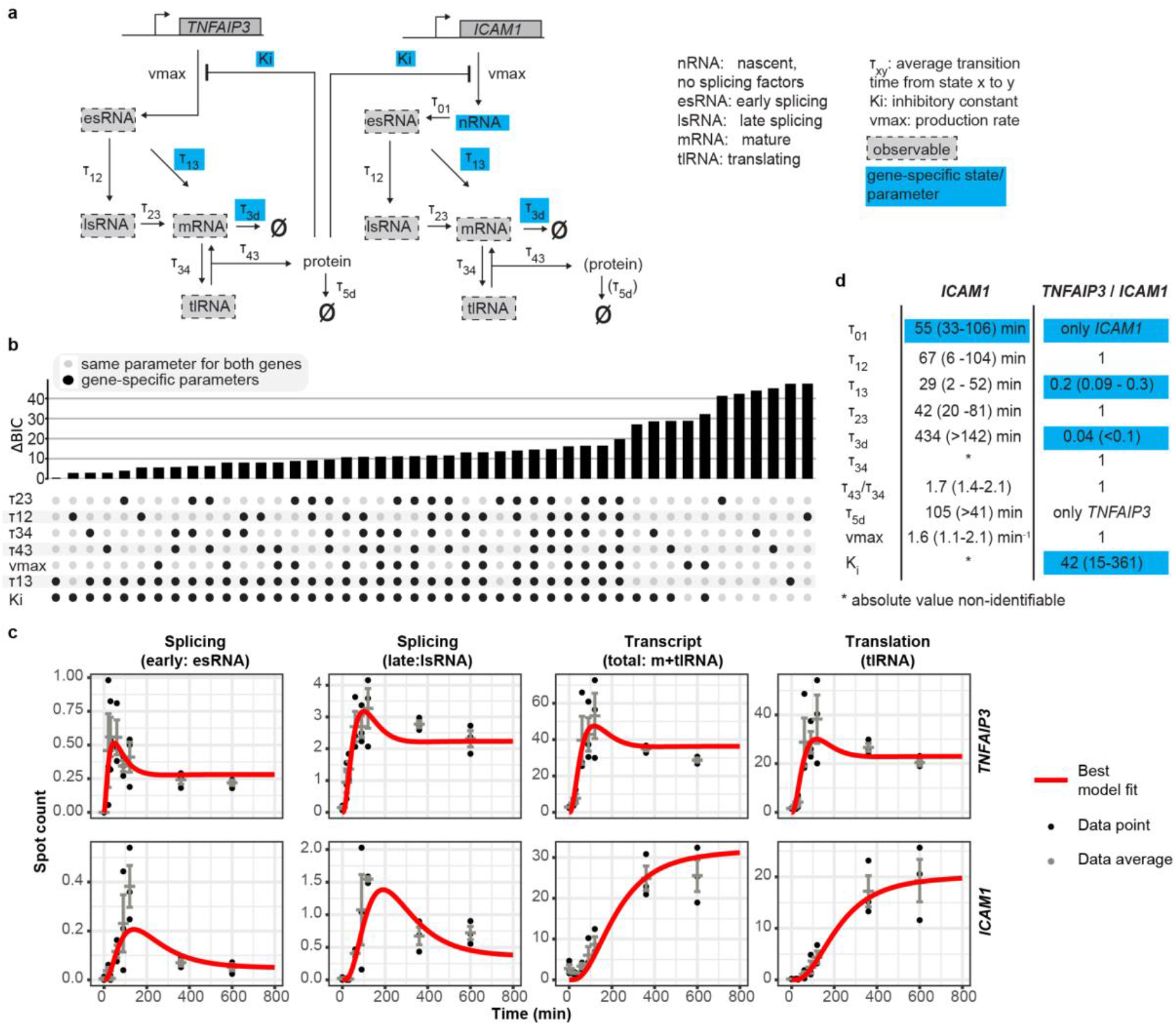
Expression steps contributing to gene-specific delay. **(a)** Gene expression model for the two TNF-α-induced genes. *TNFAIP3* and *ICAM1* have parallel, but independent states and are coupled via the negative feedback by TNFAIP3 protein. *ICAM1* features an additional delay step (nRNA) before recruitment of splicing factors. The parallel expression steps share the same kinetic parameter value, except for the gene-specific ones (cyan). The decision for shared or gene-specific parameters was based on the model selection shown in **b**. ICAM1 protein and its degradation (in brackets) were not explicitly modeled. **(b)** Systematic selection of models. According to model preselection (**Fig. S13a**) RNA degradation (τ3d) and splicing factor recruitment (for ICAM1, τ01) needed to be gene specific. Each model (shown per column) had a combination of shared (black dots) and gene-specific (grey) parameter values. The Bayesian Information Criterion (BIC) describes the goodness of fit penalized by the number of free model parameters, shown as ΔBIC compared to the best model. **(c)** Fit of best model (depicted in **a**) for the data shown in Fig. 6. The model states corresponding to the respective observable are shown in brackets. Note that the model was fitted to average spot counts of responding cells (shown in **Fig. S13b**). Model trajectories were multiplied by the responder fraction to make them compatible with the population data. Total transcripts comprise the sum of translating (tlRNA) and free mRNA. Error bars s.e.m. **(d)** Parameter estimates (maximum likelihood estimate) for best model and their 95% confidence intervals (from profile likelihood estimates, in brackets). The left column shows the parameter for *ICAM1*; the second column the ratio of parameter values for *TNFAIP3* and *ICAM1*. In cases in which the confidence interval of the ratio does not include 1 the parameter is assumed to be gene specific.

The model contains states for early and late splicing of RNA (corresponding to the TS-bound and dispersed spots we had observed), mature mRNA and translating transcripts and a negative feedback loop mediated by TNFAIP3 protein. Some modeling decisions were based on a model pre-selection (**Fig. S13a**). Among these, was the inclusion of an additional delay step in production of early splicing *ICAM1* transcripts that reflects their later appearance compared to *TNFAIP3,* which might stem from a slower activation of the ICAM1 promoter. Systematic model discrimination found positive evidence (Bayesian Information Criterion difference of 2.3, **Fig. 7b**) that early splicing (characteristic time τ_13_) and the negative feedback (inhibitory constant K_i_) were gene specific, in addition to the mRNA stability (life time τ_3d_). The selected best model was able to capture the dynamics of all observables up to the 600 min time point (**Fig. 7c, Fig. S13b**) and the estimated parameters and their confidence intervals confirmed that the initial production delay, early splicing, negative feedback and mRNA stability had gene-specific values, whereas the other expression steps were proceeding with the same kinetics for both genes (**Fig. 7d**).

We concluded that the control over gene-specific induction dynamics is spread over multiple steps of expression: For *ICAM1* there is an initial delay in the production of splicing-competent transcripts. The relatively long transition times from early to late splicing (τ_12_, 67 min) and from late splicing to mRNA (τ_23_, 42 min) indicate that for both genes the main splicing route is via TS-bound splicing and that this step is faster for *TNFAIP3* (τ_13_, 5.8 min vs. 29 min for ICAM1). Since the feedback mechanism ultimately reduces NFκB activity which controls both genes, the difference of the inhibitory constant K_i_ could potentially stem from a higher transcription factor requirement of the *ICAM1* promoter. Another major difference between the genes is their transcript stability, which leads to a gradual accumulation for *ICAM1* (life time τ_3d_, 434 min) and allows an overshoot for *TNFAIP3* (τ_3d_, 17 min).

To confront the modeling insights with a perturbation of the TNF-α response, we checked the RNA life-cycle states after transcription inhibition with Actinomycin D at 60-120 min time points (**Fig. S14a**). Kinetic simulations using the model suggested that early splicing should be the most susceptible stage (**Fig. S14b**). Indeed, for both genes transcript numbers in nearly all life-cycle stages were reduced, but the reduction was strongest for the cluster splicing spots (corresponding to the early splicing state in the model) and was comparable for both genes (**Fig. S14c-e**). While a higher turnover of fast-splicing TNFAIP3 transcripts is expected, the inhibition window at 60-120 min after TNF-α exposure coincides with the peak of ICAM1 splicing activity (**Fig. 6b**) explaining the strong effect on the splicing stage for both genes. Inhibition caused a more pronounced reduction for cluster spots compared to dispersed, consistent with the view of TS-bound splicing being the main splicing route, and dispersed splicing serving as a slower and potentially less efficient overflow mechanism.

## Discussion

Proximity detection methods provide powerful frameworks for interrogating molecular interactions, local environments and functional states within their native cellular context. Here, we established Proximity Primer Exchange Reaction (ProPER), which, to our knowledge, is the first spatially resolved in situ implementation of ligation-free proximity writing concept (**Supplementary Movie**).

With the flexibility and robustness provided by PER-based isothermal signal amplification and modular use of concatemeric DNA branches, ProPER offers high detection efficiency and versatility for various applications including background suppression, interaction partner detection or molecular state annotation for nucleic acid and protein targets. Compared to isPLA, ProPER achieved higher coverage and labeling density of proximity sites while producing more compact fluorescent products. These properties are crucial when structural and morphological context must be preserved, particularly when resolving dense targets or interrogating multiple proximity-defined states within the same physical region or on the same molecule. To support simultaneous multiplexing, we validated and characterized more than five orthogonal PER pairs for in situ proximity writing and provide an additional set of 40 previously designed pairs^25,27,28^ in **Supplementary Table S1**.

In this work, we focused on quantitatively visualizing the lifecycle stage of protein-coding transcripts based on their proximity to functional non-coding RNAs, including splicing factors U1 and U2, and 18S rRNA. Recently, a similar approach combining STARmap Plus^45^, TEMPOmap^46^, and RIBOmap^14^ for in situ sequencing of total, nascent and translating transcripts were presented^15^. However, these assays had to be performed on consecutive tissue sections, rather than on the same molecules, largely due to inefficiency of detecting the same molecule with two orthogonally generated RCA products. ProPER therefore offers a distinct advantage: multiple orthogonal proximity extension reactions can be performed simultaneously in the same cell and even on the same molecule. We used this capability to design a simultaneous detection scheme that resolves four molecular proximity states at once. In this implementation, we tiled the target RNA with four probe sets that are barcoded for docking of orthogonal PER primers. We reserved one hairpin for high-confidence coincidence detection of the target RNA and docked the other cognate hairpins on three life-cycle marker RNAs. Using this approach, we quantitatively studied the cellular response to TNF-α stimulation in human cells by resolving the RNA life-cycle dynamics for the slow response gene, *ICAM1,* and a fast response gene, *TNFAIP3*.

This application illustrates how ProPER can convert molecular proximity measurements into quantitative state-resolved models of gene regulation. Overall, the regulation of induction dynamics was observed to be distributed over several steps of gene expression. We found that low RNA stability and fast splicing at the TS enabled the *TNFAIP3* gene to have more rapid and transient induction kinetics than the similarly sized *ICAM1* gene, which has a similar number of introns. This confirmed conclusions from similar studies in mouse cells^41,42^. In addition, we found an initial delay in production of splicing-competent nascent RNA and higher sensitivity to negative feedback for *ICAM1*.

For the RNAs that are interacting with U1- or U2-snRNAs, we were able to distinguish between transcription burst site bound splicing events (0 to 2 large spots per cell) and dispersed splicing events that happen further away from the gene loci (smaller spots). Note that this definition partially overlaps but is not completely synonymous with co- and posttranscriptional splicing defined by its occurrence before or after cleavage of the 3’ end of the nascent RNA^36,47^. Here we relied on the proximity to spliceosomal snRNAs to define the splicing states of RNAs, but in future implementations ProPER could be also used for more refined splicing annotations based on individual intron-exon and exon-exon junctions. Beyond splicing, additional interactions of the spliceosomal snRNAs, like telescripting, have been defined, but are typically more relevant for longer genes than the ones studied here^48^. Our kinetic model indicated that late splicing (dispersed) outside of the TS had a smaller contribution to mature transcript production, as it was slower than TS-bound splicing overall. The rate of late splicing was not differential between the two genes and was less responsive to transcription inhibition compared to TS-bound splicing. Post-transcriptional splicing dispersed in the nucleoplasm could, thus, be an overflow mechanism when high transcription rates exceed the splicing capacity^47^. Lingering transcripts that are not fully spliced may also become substrates for nuclear RNA degradation^49,50^, however, our modeling indicated that inclusion of a rate constant for degradation of the late splicing pool does not further improve the fit of the model. Generally, association of transcripts with U1- or U2-snRNAs was not strictly confined to nuclear speckles, as defined by the marker SON. However, late splicing of *ICAM1* tended to take place inside or adjacent to speckles, whereas late splicing *TNFAIP3* transcripts were localized further away from speckles. The differential association of *TNFAIP3* and *ICAM1* transcripts with speckles could be linked to gene specific splicing kinetics and previous studies reported that speckle-enriched transcripts contained inefficiently spliced introns^37^.

Single-cell level analysis revealed that the heterogeneity of gene expression response across cells mainly arose from the transition from a non-responder to a responder state. High splicing levels were correlated with high total transcript levels indicating that splicing remains active as mature transcripts accumulate. This is consistent with a digital activation process in which first-responder cells contribute to the early activation dynamics of the population^44,51^.

The high labeling density, multiplexing capacity, and autonomous isothermal design of ProPER enable one-to-many proximity detection schemes that allow for the simultaneous resolution of diverse interaction partners and molecular/functional states on individual targets. Although our initial application focused on RNA life-cycle progression, particularly the splicing and translation stages, the same framework could be adapted to probe additional molecular states, including polyadenylation, RNA transport (e.g. based on proximity to RNA transport or sequestration machinery^52^), or RNA modifications (through metabolic labeling or modification-specific antibodies). More broadly, ProPER is readily applicable to protein-protein interactions as well as multimodal molecular-state measurements spanning proteins, RNAs and other biomolecules, enabling integrated analysis of diverse molecular interactions within their native spatial context. As exemplified by our licensing strategy (**Fig. 4f-h**) and previous in-solution implementations of PER^25^, ProPER offers programmability for complex proximity-driven logic, such as multivalent or multi-way detection, and is well suited for future many-to-many interaction detection implementations. For higher throughput, validated ProPER barcode sets could be combined with combinatorial imaging strategies, enabling simultaneous detection of multiple proximity-defined molecular states alongside highly multiplexed imaging of transcriptome using methods such as MERFISH^53^, SeqFISH^54^, HybRISS^55^. In conclusion, ProPER addresses longstanding technological limitations in spatial proximity detection and, by enabling multiplexed molecular-state analysis, expands the scope of spatial biology beyond molecular abundance and localization toward quantitative molecular-state measurements. By converting molecular proximity into programmable DNA readouts, ProPER provides a foundation for future higher-order encoding schemes and integration with imaging- and sequencing-based spatial omics workflows.

### Limitations of the study

It is important to note that like existing proximity detection technologies, ProPER reports the physical proximity of biomolecules, which is commonly interpreted as a molecular interaction, but does not definitively prove direct binding between the two molecules or activity of their complex.

Furthermore, while ProPER enables highly sensitive visualization of target proximities, absolute quantification is not always directly feasible for all modalities. For example, unlike for detection of single RNA spots, in protein-protein proximity detection the overlapping distribution of proteins, variable stoichiometry of interactions and bulky nature of antibodies can impose steric hindrances and amplification heterogeneities that hamper absolute quantification. In our RNA life-cycle imaging, transcription site-associated splicing spots naturally do not represent true single-molecule detections, as transcriptional bursting leads to the localized accumulation of multiple nascent transcripts at various stages of processing. To account for this in our quantitative kinetic modeling, we introduced multiplicative scaling parameters that successfully converted the molecule counts of the underlying state variables into the observed spot counts.

Finally, our control experiments identified the no-primer-docking condition, in which all reaction components including the primer are included but docking site for the primer is omitted, as the most stringent control for background. Generally, this background was negligible compared to the true proximity signal, but it could increase when the hairpin is presented on highly abundant targets, like ribosomes, or is locally concentrated in the same spot. In these cases, similar to PLA^56^, ProPER had false-positive detections which was most pronounced for the very high abundance target, the ribosome component 18S rRNA. Designing detection schemes that avoid high local concentration of the hairpin and optional intensity thresholding largely alleviates this problem. Additionally, we employed three different approaches which all helped reduce false-positives: (1) optimizing the primary probe performance on the proximity target, (2) introducing a licensing step with the licensing hairpin on the more abundant docking site and the extender hairpin on the (less abundant) transcript, (3) multiplexing with an orthogonal ProPER reaction for background suppression and filtering detected spots based on their overlap with the total transcript channel.

## Methods

### Design of primary probes

Primary probes for the RNA of target genes as well as life-cycle marker RNAs (18S, U1, U2) were selected from the oligoMiner hg38 newBalance database^57,58^ by a custom built pipeline implemented in R^59^: Coordinates of exonic regions of the target gene were retrieved from Ensembl genome annotation GRCh38.105. Exons were included starting from the 5’-end covering at least 2500 bases and probes in the selected regions were collected. Probe quality thresholds (on-target score, off-target score, probability for secondary structure) were iteratively lowered until the desired probe number was reached. Maximum values of 100 and 10 were allowed for the off-target score and the number of off-target kmers, respectively. If the desired probe number could not be reached with the selected sequence window, exons covering 11,000 bases from the 5’-end were included.

Selected probes were appended with docking sites on the 5’- and 3’-end or only on the 3’-end for probes targeting life-cycle stage markers. For target RNAs the docking sites were adapted from 20-nt MERFISH sequences^60^. For life-cycle marker RNAs the docking sites were 30-nt SABER sequence repeats. Adopting the previously published nomenclature^27^, the PER sequences are designated by two-digit numbers between 25 and 75 and the number of repeats is specified, e.g. 3x25 for 3 repeats of PER-primer sequence 25. Note that both types of docking sites have similar melting temperatures due to a higher G/C content of the shorter MERFISH sequences. Target gene probes were assigned to two sub-pools that alternated along the genomic coordinate. These subpools were appended with docking sequences M03 and M06, or M05 and M10. Probes for life-cycle markers were ordered as individual DNA oligos and probes for target genes as opools from Integrated DNA Technologies Inc. (IDT). Life-cycle marker probes for multiplex experiments were ordered with HPLC purification. Probe sequences are listed in **Supplementary Table S1**.

### Branches, hairpin, primer and imager probes

Primary probes on the target gene provide four different types of docking sites (adapted from MERFISH sequences), which bind 200-nt concatemeric branches for binding of primers or hairpins (with 30-nt branch-binding domains of 3-letter code typically on 5’-end, adapted from SABER sequences) and thereby amplify the number of probe pairs available for in situ PER. Imagers were 20-nt SABER sequences in 3-letter code with the fluorophore on the 5’-end. Hairpins and imagers had an inverted dT at the 3’-end for end protection. Imagers, hairpins and primers were ordered from IDT with HPLC purification. Sequences are listed in **Supplementary Table S1**.

### Antibody–DNA Conjugation

Donkey anti-mouse IgG (H+L) (Jackson ImmunoResearch, 715-005-150) and donkey anti-rabbit IgG (H+L) (Jackson ImmunoResearch, 711-005-152) antibodies were conjugated to 5′-thiol–modified DNA oligonucleotides (Integrated DNA Technologies) as described before^28^. Oligonucleotides were reduced by incubating 25 µL of 1 mM thiol-DNA with 20 µL of 500 mM dithiothreitol (DTT; Thermo Fisher, 20291) in PBS– EDTA (pH 8.0) for 2 h at room temperature, protected from light. Reduced oligonucleotides were desalted on NAP columns (GE Healthcare, 17-0853-02) equilibrated with nuclease-free water, and fractions containing the highest DNA concentrations (∼100 µM) were pooled. Antibodies (100 µg) were buffer-exchanged into PBS using Amicon Ultra 50 kDa centrifugal filters (Millipore, UFC510096) and reacted with a 4.5-fold molar excess of SM(PEG)₂ crosslinker (Thermo Fisher, A35397) freshly prepared at 2 nmol /µL in DMF for 1.5 h at 4 °C. Excess crosslinker was removed using Zeba Spin Desalting Columns (Thermo Fisher) equilibrated with PBS. Subsequently, reduced thiol-DNA (7.5 nmol; molar ratio antibody:DNA = 1:11) was added to the crosslinker-activated antibodies and incubated overnight at 4 °C, protected from light, to generate DNA–antibody conjugates. The conjugates were washed and concentrated by repeated PBS filtration (4×) using 50 kDa Amicon filters and stored at 4 °C until use. Successful conjugation was confirmed by non-reducing SDS– PAGE. For primary antibodies, alternative methods of adapter-based conjugation can also be utilized^61^.

### Pre-extension of primary probes by in vitro PER

Primary probes with concatemeric docking sites were produced by in vitro primer exchange reaction as previously described^27,28^. Final concentrations in 200 µL reaction volume were 1× PBS, 480U/ml Bst LF polymerase (NEB, M0275M), 10 mM MgSO₄, 0.6 mM dNTPs (dATP, dCTP, dTTP), 0.1 µM Clean.G hairpin (IDT), 0.6-1.0 µM hairpin and 0.75 µM primer. Exact hairpin concentration was chosen to empirically reach 300 – 500-nt long concatemers. The reaction mix without primers was incubated at 37 °C for 15 min. Diluted primers were then added, followed by 37°C for 2 h and 80 °C inactivation (20 min). Products were purified with MinElute columns (Qiagen; eluted in 25 µL water/IDTE) and stored at -20 °C. Products were analyzed on a 1% E-Gel EX gel (Thermo Fisher, G402001).

### Cell culture

Normal human lung fibroblasts (NHLF, Lonza, CC-2512) were cultured in Fibrolife medium (Lifeline Cell Technology, LL-0011), supplemented with 5% fetal bovine serum, the provided growth factors and antimicrobials and without hydrocortisone, L-glutamine or ascorbic acid supplementation. Lonza obtained informed consent for research use of the isolated primary cells from donated human tissue. HeLa cells were cultured in DMEM (Gibco, 41965039) with 10% fetal bovine serum and 100 U penicillin-streptomycin (Gibco, 15140-122).

Cells were seeded in collagen coated (rat tail collagen I, Roche,11179179001) 18 -well glass bottom slides (ibidi µ-slide, 81817). The next day cells were exposed to 10 ng/ml TNF-α (RD Systems, 10291-TA-020) in Fibrolife. For time course experiments TNF-α exposure started at the respective time points before fixation. For the metabolic labeling experiments 5-ethynyl uridine (Jena Bioscience, CLK-N002-10) was added 30 min before fixation to a final concentration of 1 mM. For the transcription inhibition experiments Actinomycin D was added one hour before fixation to a final concentration of 0.5 µg/ml. At the fixation time point an equal volume of 4% formaldehyde (Pierce, 28908) was added to the media for pre-fixation over 5 min (2% final formaldehyde concentration). After washing with PBS cells were fixed for 10 min in 4% formaldehyde, washed with PBS and stored in 70% ethanol (diluted in nuclease free water) at -20 °C.

### EU click reaction

The hairpin docking site was attached to the metabolically labeled nascent RNA in EU-treated cells by click reaction with an azide-modified oligo using click chemistry reagents from the Clicktech EdU imaging kit (Baseclick, BCK-EdU647IM100). Cells were permeabilized with 0.5% Triton-X-100 in PBS for 10 min. The click oligo (5’azide-3x25, 100µM in IDTE buffer [IDT, 11-05-01-09]) was pre-annealed 5 min at 50°C with a short blocking oligo (25*-15mer, 100 µM in IDTE) in a 1:2 molar ratio and then cooled down to room temperature. The click chemistry solution was prepared according to the manufacturer’s instructions and 27 µL of pre-annealed oligo were included in 1 ml solution. The reaction mix was added to the cells for 60 min at room temperature followed by washes with 1×PBS, three times formamide buffer (50% v/v formamide, PBS, 0.1% Tween-20), three times high-salt buffer (PBS, 1M NaCl, 0.1% Tween-20) and twice 1×PBS. For direct visualization of the EU-staining (“EU-FISH”) imager oligos binding to the 3x25 docking site were hybridized. In EU-ProPER the docking sites were used for hairpin hybridization.

### ProPER: Primary and secondary probe hybridization

Buffers were prepared with nuclease free water (Invitrogen, 10977-035). Fixed cells were washed once in phosphate-buffered saline (PBS, Invitrogen, AM9625) for 1 min at room temperature (RT) to remove storage ethanol, followed by permeabilization in PBS containing 0.5% Triton X-100 (Sigma Aldrich, T8787) for 10 min at RT. Samples were subsequently washed for 1 min in PBS with 0.1% Tween-20 (PBSTw, Sigma Aldrich, P9416).

After washing twice for 1 min in 2×SSC (Invitrogen, AM9763) containing 0.1% Tween-20 (2×SSCT) at RT, cells were incubated in hybridization buffer (2x SSC, 50% formamide [Ambion, AM9342], 10% dextran sulfate [Sigma Aldrich, D8906], 8mM Ribonucleoside Vanadyl Complex [VRC, NEB, S1402S], 0.4 mg/ml sheared salmon sperm DNA [Invitrogen, AM9680], 1 mg/ml yeast tRNA [Sigma, R8759], 0.1% Tween-20, 0.1% Triton) containing 80nM each of life-cycle target probes (U1, U2, 18S) and of the transcript targeting probe pool (*TNFAIP3*, *ICAM1* or *CBX5* probe pools). After denaturation at 60 °C for 3 min, hybridization was carried out at 37 °C overnight. Following hybridization, slides were washed on a flat-top thermocycler (Eppendorf): 150 µL of pre-warmed (60 °C) 2×SSCT was added, and samples were washed 4 × 5 min in pre-warmed 2× SSCT at 60 °C, followed by 2 × 2 min in 2× SSCT at RT.

Secondary probe hybridization of branches, hairpins and primers was performed for 1 h at 37 °C in secondary hybridization buffer (2×SSC, 25% formamide, 10% dextran sulfate, 0.2 mg/ml sheared salmon sperm DNA, 1 mg/ml yeast tRNA, 0.1% Tween-20). When appropriate, for cost efficiency, converter oligos were added to convert a docking sequence to a compatible sequence for the respective primer or hairpin. In experiments in which pre-extended primary probes (PER concatemers) were used (for *CBX5* and for EU experiments, **Fig. 1c-f**, **Fig. S5**) no branches were needed. The final concentration of each probe was 10 nM. After hybridization, 100 µL pre-warmed (45 °C) PBSTw was applied directly onto wells, which were then aspirated. Washes were done 3 × 5 min at 45 °C with pre-warmed PBSTw buffer. Samples were allowed to cool down to RT for 5 min to facilitate initial annealing of hairpin and primer before continuing with in situ PER. The combinations of probes for each individual experiment can be found in **Supplementary Table S2**.

### ProPER: In-situ PER

PER reaction solution (1×PBS, 5 mM MgSO_4_ [NEB, B1003S], 2.4 mM dATP, 2.4 mM dCTP, 2.4 mM dTTP [Carl Roth, K039.1], 1 μM cleanG hairpin, 1440 U/ml Bst I polymerase [NEB, M0275M]) was prepared and incubated 15 min at 45 °C to remove any dGTP contamination via the activity of the cleanG hairpin before adding the solution to the samples. The in-situ PER proceeded 2 h at 45 °C. After the reaction samples were washed 3 × 5 min in PBSTw at 37 °C. In the early optimization experiments lower concentrations of MgSO_4_ and dNTPs were used (2.5 mM and 1.2 mM, respectively).

### Imager hybridization

For multiplex detection, PER sequence-specific imagers were hybridized. Cells were rinsed once with PBSTw, incubated for 1 h at 37 °C in imager hybridization solution (200 nM imagers in PBS, 0.5% dextran, 0.1% Triton X-100, 1 mg/ml yeast tRNA), and washed 3 × 5 min in pre-warmed (45 °C) PBSTw. Samples were rinsed once at RT and stained with 5 µg/ml DAPI (ThermoScientific, 62248) for 10 min, then washed and stored at 4 °C in storage buffer (PBS, 1 M NaCl, 0.02% sodium azide) until acquisition.

### Immunofluorescence stainings after ProPER

Following ProPER labeling, cells were washed once in PBS and blocked in PBS with 2.5% (w/v) bovine serum albumin (BSA) containing 0.5% Triton X-100 for 15 min. Samples were incubated with anti-SON antibody (Proteintech, 83787-5-RR, rabbit recombinant) diluted 1:100 in PBS with 2.5% BSA for 1 h at RT. Following three washes in PBSTw (2 min, 5 min, 2 min), the secondary antibody (Alexa Fluor 750 anti-rabbit IgG, Invitrogen A-21039) diluted 1:250 in 2.5% BSA was applied for 30 min at RT. Cells were washed once with 1×PBS. Nuclei were stained with DAPI (5 µg/mL, 10 min), washed in 1×PBS, and stored in storage buffer until image acquisition.

### Protein-protein ProPER

NHLF cells were processed for protein–protein interaction analysis using antibodies targeting acetylated α-tubulin (mouse, #T7451, Sigma Aldrich, 1:125) and β-tubulin (rabbit, 10068-1-AP, Proteintech, 1:100). Cells were permeabilized and blocked in 2% BSA and 0.3% Triton X-100 in PBS for 60 min. Primary antibodies were applied in IF buffer (2% BSA, 0.1% Triton X-100, PBS) for 1 h at room temperature, followed by three 1×PBS washes. Secondary ssDNA-conjugated antibodies (anti-mouse bc09 and anti-rabbit bc30, prepared as described in the Antibody-DNA Conjugation section above) were blocked by pre-annealing with short 9mer oligos in secondary staining buffer (PBS, 2.5% BSA, 0.1% Triton X-100, sheared salmon sperm DNA, 2% dextran sulfate, 1:62.5 of each conjugated antibody and 1 μM blocking oligo complementary to the barcode sequences) at 50 °C for 5 min and equilibration at room temperature. The solution was applied to the samples for 1 h. Samples were washed in blocking buffer and three times in 1×PBS and post-fixed with 5 mM Bis(sulfosuccinimidyl)suberate (BS3, ThermoScientific, 21580) in PBS for 30 min, quenched with 100 mM ammonium chloride (Sigma-Aldrich, 09718) solution for 5 min, and blocking oligos were stripped by washing three times in 60% formamide in PBS followed by three 1×PBS washes. Probe hybridization was performed at 37 °C for 1 h in secondary probe hybridization buffer (composition as described above for RNA ProPER, but with 20 nM each for docking probe primer and hairpins). After hybridization, 100 µL pre-warmed (45 °C) PBSTw was applied directly onto wells and then aspirated. Washes were done 3 × 5 min at 45 °C with pre-warmed PBSTw buffer. In situ primer exchange reaction was carried out for 2h at 45 °C as described above in the In-situ PER section. Finally, fluorophore-labeled imagers (39*-565, 30*-488, and 28*-647) were hybridized at 37 °C for 1 h under the conditions described in the Imager hybridization section, followed by brief washes in prewarmed PBS-Tween, DAPI staining (5 µg/ml, 10 min), and imaging in PBS.

### Protein-protein isPLA

The proximity ligation assay was performed using the Duolink In Situ Red Starter Kit Mouse/Rabbit (Sigma Aldrich, DUO92101-1KT). Primary antibody staining of NHLF cells was done in parallel with the protein-protein ProPER experiment using the same antibodies: Cells were permeabilized and blocked in 2% BSA and 0.3% Triton X-100 in PBS for 10 min and then blocked for 60 min with Duolink blocking solution. Primary antibodies were applied in Duolink antibody diluent for 1 h at room temperature, followed by three 1×PBS washes. From this point on isPLA was performed following the manufacturer’s instructions. Briefly, samples were washed three times with Wash Buffer A and incubated with Plus and Minus PLA probes in a 1:5 dilution (in Antibody Diluent) for 1 h at 37 °C. After three washes with Wash Buffer A, ligation was carried out in Ligation Buffer for 30 min at 37 °C. The samples were washed two times with Wash Buffer A, then Amplification Buffer was added for 100 min at 37 °C. Finally, two washes with Wash Buffer B and one short wash with 0.1x Wash Buffer were done and the samples were mounted with the manufacturer’s mounting medium containing DAPI.

### Microscopy

Samples were imaged on an inverted Nikon Ti2 microscope in 1×PBS or storage buffer (PBS, 1 M NaCl, 0.02% sodium azide). The microscope was equipped with a CFI Plan Apo Lambda D 60× Oil objective (numerical aperture 1.42), Crest X-Light V3 confocal spinning disc unit (50 µm pinhole size), a Celesta laser light source with Despeckler (Lumencor) and a Kinetix Back Illuminated sCMOS camera (Photometrics). Laser lines 405 nm, 477 nm, 546 nm, 638 nm and 749 nm were used together with the DA/FI/TR/Cy5/Cy7 emission filter set (Celesta) to image DAPI and imagers labeled with ATTO 488, ATTO 565, ATTO 647 and Alexa 750. Typically, five fields of view were selected at distributed positions in the well. Z-stacks spanned 10 µm with 0.5 µm step size. For RNA life-cycle imaging and some other experiments 2x2 pixel binning was employed to boost signal strength (required especially for Alexa 750).

*CBX5* samples (**Fig. 1c-f**) were imaged on a Nikon Ti2 inverted microscope equipped with an ORCA-Fusion CMOS camera and a Lumencor SPECTRA III illumination system. Imaging was performed using a CFI P-Apo 20× Lambda air objective with a numerical aperture of 0.75. Excitation was provided using the following filter sets: 393/25 nm for DAPI, 555/28 nm for ProPER/SABER. A penta beamsplitter with a multiband emission filter was used for all imaging channels, in combination with pentaband emission filters mounted in the filter wheel (432/36, 515/30, 596/30, 682/42, and 809/89 nm). For each well, four fields of view were arbitrarily selected. The focal plane corresponding to the maximum intensity in the DAPI channel was identified as the central plane, and a z-stack spanning 7 µm centered on this plane was acquired with a step size of 1 µm.

For experiments that needed higher resolution (protein-protein ProPER/PLA, **Fig. 2a-f**, and U1/2-ProPER with speckle co-staining, **Fig. S8**) samples were imaged on a STELLARIS confocal microscope (Leica Microsystems) equipped with a HC PL APO 63×/1.40 OIL CS2 objective (Leica Microsystems), a diode laser at 405 nm and a white light laser with tunable excitation (440–790 nm). 8.75 µm z-stacks with a 0.25 µm step size and pixel size 90 nm were acquired. PMT detectors were operated in photon counting mode. For the U1/U2+SON co-staining experiment laser lines used were 405, 579, 638 and 750 nm with detection windows 425-480 nm, 580-650 nm, 660-690 nm and 750-850 nm. For protein-protein ProPER (**Fig. 2a-f**) the same settings applied but using additionally an ATTO 488 channel with laser excitation at 488nm and a detection window 500-550 nm. For PLA the 594 nm laser line was used with a detection window 615-655 nm.

### Tubulin fragment analysis

Tubulin fragment detection was performed on single plane images of isPLA and ProPER signal using EBImage^62^ in R^59^. Images were median-filtered and processed with a custom single-scale 2D Frangi vesselness filter^63^ (σ=3, β=0.25, c=15) based on Hessian eigenvalues from second-derivative Gaussian convolutions to detect ridge-like structures, yielding binary masks via thresholding and connected-component labeling. Small objects (<50 pixels) were excluded as noise. Object eccentricities were extracted with the EBImage function computeFeatures.moment for remaining fragments. Values of this quantity close to 0 indicate circular objects, whereas values close to 1 indicate fiber-like structures. Labeled masks were exported as pseudocolored images.

### Image processing for spot quantification

Image preprocessing combined 2D nuclei and cytoplasm segmentation, intensity normalization and artifact removal. The whole analysis workflow from raw images to spot counts is visualized in **Fig. S7**.

For each field of view a maximum intensity projection across z was generated. Cell nuclei were segmented on the DAPI channel using the cyto model of Cellpose (v2.1.1)^64^ on a high performance computing cluster with a fixed diameter parameter of 65 pixels and default model settings, producing 2D nuclear label masks for each field. For each nucleus mean DAPI intensity was computed with EBImage’s^62^ computeFeatures.basic function in R, and nuclei with weak DAPI signal (mean intensity below a global threshold) were excluded to remove low quality nuclei across all conditions.

To obtain whole-cell regions, the background signal of the four non-DAPI channels was used. For this purpose, nuclear masks were combined with multi-channel background fluorescence using EBImage. For each image, the four non-DAPI channels were mean-normalized, smoothed (Gaussian blur), locally background-subtracted by a broad Gaussian filter, and thresholded to define a permissive segmentation space that was dilated with a disc kernel. Nuclear masks were slightly dilated to create seed regions, and used as seeds for EBImage’s propagate function, yielding cytoplasmic label masks aligned to the nuclei within the allowed segmentation space. A unique background label at the image borders was included in the seed propagation step to avoid cytoplasm objects touching the image borders. In cases in which nuclei were touching image borders they were removed together with their corresponding cytoplasm mask.

For each fluorescence channel (1–4), cytoplasmic regions were used to estimate intensity statistics and define normalization factors. Specifically, for every image and channel, mean and quantile features were computed over cytoplasmic masks using computeFeatures.basic (EBImage), aggregated per condition and replicate, and the median of the cell intensities was used to compute a replicate- and condition-wise normalization factor that equalized mean cytoplasmic intensity across experimental replicates. Normalized images were then created by dividing each image by the normalization factor. Local background removal was achieved by subtracting the image after Gaussian blurring. To suppress sporadic bright imaging artifacts such as reflections, normalized images were thresholded, labeled, and objects larger than 70 pixels were dilated and converted to a binary mask, which was then used to replace affected pixels with the global mean of the filtered image.

### Spot detection

Spots were detected using the RS-FISH plugin for Fiji^65^, which implements radial symmetry-based spot localization. For each condition and channel, artifact-corrected, normalized images were processed twice with RS-FISH with Gaussian sigma values of 1.5 and 2.5 to capture small and slightly larger spots, respectively. For each experiment channel-specific intensity thresholds were chosen based on representative images from high-expression and negative control conditions. These thresholds were set leniently allowing for further intensity- or overlap-based filtering in later analysis steps. In some conditions the thresholds led to the exclusion of a low-intensity population of spots present in a punctate background pattern. Spot coordinates were saved and provided the basis for subsequent assignment of spots to nuclear and cytoplasmic masks and spot intensity measurements.

### Spot processing

Spot post-processing was performed in R using EBImage for image handling and feature extraction. Most analysis steps were identical for the ProPER method optimization/validation experiments and the full fourplex RNA life-cycle experiments. However, some of the following steps applied only to RNA life-cycle ProPER (**Fig. 5,6**, e.g. spot size classification, multichannel or compartment assignment). Multichannel spot assignment was also used for the two-color smFISH and smFISH+ProPER experiments (**Fig. 2g-i**).

### Spot measurement and compartment assignment

Border-proximal spots were removed, and each remaining spot was assigned a cell ID (from the cell mask) and a compartment label (“nucleus” or “cytoplasm”) based on the nuclear mask. Around every spot, a square patch of fixed radius on the corresponding normalized image was extracted and convolved with circular masks of different diameters (small, big, very big) to measure local mean intensities as well as intensities in concentric radial shells for radial intensity profiles.

### Reference-channel anchoring and removal of overlapping spots

The fluorescence channel of the total transcript signal was designated as a reference. For non-reference channels, only spots whose positions lay within a small Euclidean distance (3 pixels) of a reference-channel spot were kept, thereby filtering for high-confidence spots. Since spots were called twice with different parameters (to allow optimal calling of spots with variable sizes), it was necessary to remove overlapping spots in the same channel. Within each cell and channel, pairwise distances between remaining spots were computed and spot pairs closer than a threshold (3 pixels) were considered overlapping. In each overlapping pair, the weaker spot was removed, resulting in a non-overlapping set of spots per cell and channel. Reference-channel spots below a lower intensity threshold were discarded and all channels were filtered to exclude spots with large-region intensity above an upper cutoff to remove large artifacts.

### Spot size measurement and spot-type classification

Spots were classified as small (“dispersed”) or large (“cluster”) spots based on their radial intensity profiles. To make this classification robust to variable intensities (between experiments or due to local background levels) profiles were normalized between their center intensity and the baseline intensity which was defined per gene, experiment and channel. This baseline was calculated as the knee-point of the empirical cumulative distribution functions (ECDFs) of large-region intensities. The knee point was computed by finding the maximum distance of the ECDF from the diagonal in normalized intensity–ECDF space.

To obtain a continuous estimate of spot width, the discrete radial profile was interpolated using a 1D linear spline. For each spot, the profile was first baseline-corrected by subtracting baseline value. The half-width at half-maximum (HWHM) for a given spot was then defined as the smallest radius at which the interpolated radial profile fell below half of its maximum value; if no such radius existed within the sampled range, a default HWHM of 1.5 pixels was assigned. Spots with a HWHM greater or equal to 1.75 pixels (380 nm, visually chosen to separate the two spot populations in the bimodal HWHM histograms, **Fig. 5h** and **Fig. S6f-g**) were classified as large cluster spots.

### Multicolor spot assignment and aggregation

To distinguish multicolour spots, present at the same position in multiple channels, filtered spots from all channels were merged per image and cell and a distance matrix in coordinate space was computed. For each spot, neighboring spots within a small Euclidean distance (3 pixels) were identified, the set of channels present in this local neighborhood was recorded as a multichannel label, and close-contact spots were further pruned by retaining only the largest (by half-width-half-maximum, HWHM) spot within groups of very closely spaced detections.

### Spot counting

For each image, cell-resolved counts were derived by summing, per cell and channel, the total number of filtered spots, numbers of big and small spots, and their compartment-specific subsets (nuclear vs. cytoplasmic).

In the case of multicolor spot counting, per-cell counts were computed for each distinct multichannel combination and were stratified by spot type (cluster/disperse) and compartment. These combination-specific counts were then summed into biologically interpretable categories such as total, cytoplasmic translation-associated spots, and cluster or dispersed nuclear U1/U2 snRNP-associated spots, which formed the basis for per-cell and per-replicate time courses and responder/non-responder statistics.

Additional quantities were derived per cell or per cell population: By dividing total transcript numbers in the nucleus by those in the whole cell the nuclear transcript fraction was obtained. Dividing translating transcript numbers by total yielded the translating fraction of transcripts. Cells were classified as responders if at least one spot type was above a basal expression threshold (total ICAM1: 3, total TNFAIP3: 5, translating ICAM1: 1, translating TNFAIP3: 3, splicing disperse: 1, splicing cluster: 0). For each gene, condition and time point, the responder fraction was then calculated as the proportion of cells labeled as responders among all analyzed cells.

### Colocalization analysis of speckles with splicing signal

Images were analyzed in R using the EBImage package for image processing and segmentation. Z-stacks of DAPI, nuclear speckle marker SON, and U1- and U2-ProPER were processed for quantitative analysis. Nuclei were segmented in the DAPI channel by Gaussian blurring followed by global intensity thresholding, 3D labeling, and removal of masks in slices that did not retain at least 50% of their maximal projected area across z, yielding a per-slice nuclear mask. Nuclear speckles were segmented in the SON channel after Gaussian filtering and thresholding within the nuclear mask. A distance transform was computed to obtain, for every voxel, the distance to the nearest speckle.

U1 and U2 ProPER spots were independently detected in their respective channels by Gaussian blurring, intensity thresholding, and masking to the nuclear volume, followed by maximum projection across z and connected-component labeling to define individual spots. For further analysis, U1- and U2-spots were combined and treated as general splicing spots. For each labeled spot, the pipeline extracted 3D centroid coordinates (z determined from the intensity maximum along the stack at the spot position), which were used to determine the distance to the nearest nuclear speckle (sampled from the speckle distance transform at the spot xy-position). The distance measurement was performed in each z-plane in a range of +/- eight planes around the spot z-center. To obtain the 3D distance, the geometric mean of xy- and z-distance was calculated (accounting for pixel anisotropy). The final 3D distance measurement was chosen as the minimum across the values for the tested z-planes. Splicing spots were classified into cluster or dispersed spots based on area thresholds, and their speckle distance distributions were plotted.

### Comparison of ProPER to high-confidence dual-color smFISH

2-color smFISH and combined smFISH + ProPER images were processed using the same pipeline as described above for RNA life-cycle images. However, no filtering based on a reference channel was performed. For each spot its presence in channel 1 and 2 was recorded as described above for multichannel spot classification. Spot counts per cell were quantified for each channel individually and for FISH dual-color spots which represent a high-confidence reference. To exclude the non-responders, only responding cells with at least one spot in each channel and one double channel spot were considered. The channel agreement of smFISH (SABER-FISH) and ProPER was then calculated as the number of ProPER or FISH-spots relative to the spot number in the comparison channel (the channel with the higher count; FISH channel 2 as the comparison channel for ProPER, FISH channel 1 as the comparison channel for two-color FISH). We also reported the ratio of ProPER detections to double-labeled FISH spots because both represent the high confidence co-detections of two probe sets that target the same transcript. We, further, quantified the agreement between channels for dual-color spots by reporting the distance distribution of spot centers in channel 1 and 2. For spot intensity distributions of ProPER and FISH intensities of dual-labeled spots in channel 1 were quantified. Only spots in the cytoplasm were used to avoid the sporadic very bright spots in the nucleus. The coefficient of variation (CV) of these intensity distributions was used to compare variability of ProPER and FISH signal.

### Image processing for display

Image z-stacks were processed in Fiji^65^. Either individual z-slices were selected or maximum projections of 3-5 slices or of whole stacks were created as indicated in figure legends. Contrast was set for optimal signal visibility (linear contrast adjustment) and for each channel the same contrast settings were applied for images from the same experiment, except if otherwise indicated. Scale bars were added according to the pixel size from the image metadata.

### Statistics and data representation

Single cell representations of RNA life-cycle data were created with the Rtsne package in R using the following cell quantities: total transcripts, translating transcripts, U1-cluster, U1-disperse, U2-cluster, U2-disperse, nuclear fraction of transcripts. After an initial PCA, cells were represented by TSNE (perplexity 250, exaggeration factor 2.0). Cells were color-labeled by time point (**Fig. 6d**) or the individual quantities (**Fig. S10e-g**, **Fig. S11**).

Experiments for technical method optimization and for the exploratory analysis of EU-ProPER (**Fig. S5**) and splicing-speckle association (**Fig. S8**) were performed in one biological replicate whereas RNA life-cycle experiments had three replicates. Replicate, cell and (where applicable) spot numbers are recorded in **Supplementary Table S3**. Measurements are from distinct samples and not from repeat measurements of the same sample. Different aspects of the data set of RNA life cycle dynamics are presented distributed over **Fig. 5-7** and the corresponding supplementary figures.

Single-cell, -fragment or -spot data was represented by dotplots (or bar graph, **Fig. S1d**) or the empirical cumulative density function (**Fig. S8c**) using ggplot2^66^ in R and the mean and standard error of the mean (s.e.m.) were indicated (except **Fig. S14e,** where 95% confidence intervals for spot counts relative to the untreated control are indicated). For comparing these distributions, two-sided Mann-Whitney-Wilcoxon tests were performed. For comparing the average values of replicate experiments, two-sided Welch t-tests were performed. Before the Welch t-test, normality was confirmed with the Shapiro-Wilk test except for negative control conditions where the test was not suitable due to the high number of zero or close-to-zero counts. Statistical tests against technical negative controls (no TNF-α stimulation, no primer dock) were performed one-sided. Average values and test statistics can be found in **Supplementary Table S3**.

The resolution limit depicted in **Fig. 2f** was calculated from the Rayleigh diffraction limit (0.61*λ/NA) using a 670nm emission for the Alexa 647 fluorophore and the 1.4 NA of the objective.

### S/B calculation

A peak-based signal to background ratio (S/B) was calculated in numpy (v. 1.26.4) (**Fig. S1a,** **Fig. 1e**). For each spot, a 3D volume of 10 pixels surrounding spot centroid was cropped. S/B was defined as “signal” (spot intensity at maximum, *I*) divided by “background” (mean intensity from the outermost layer of the volume, *μ*): *S/B = I/μ*.

#### Mechanistic modelling of RNA life-cycle dynamics

A compartmental model was developed to describe the dynamics of the RNA life-cycle stages of *ICAM1* and *TNFAIP3*, accounting for gene specific distributed response delay across individual cells in the population.

#### Response-time distribution quantification

The time delay between TNF-α stimulation and TNFα-induced gene expression was quantified at the population level, using a mixture of exponential distributions, fitted to the full single-cell response-time dataset. Models with one, two and three mixture components were evaluated, and the two-component mixture was selected as optimal based on the Bayesian Information Criterion (BIC). Mixture models were fitted to single-cell response-time data using a maximum likelihood approach. For each cell, it was known whether TNF-α induced gene expression was present by the time of measurement.

Cells classified as non-responders contribute right-censored response-time observations, indicating that the response-time is greater than the measurement time in these cells. Cells classified as responders contribute interval-censored response-time observations, indicating that the response-time lies between TNF-α stimulation (time = 0) and the time of measurement in these cells. The likelihood function was constructed accordingly from the cumulative distribution functions of the exponential mixture model to account for the censoring structure of the data. Fitting of the parameters of the response-time distribution was performed in R. Further implementation details are provided in **Supplementary Note**.

#### Kinetic model of the RNA life cycle

The dynamics of the two-gene system were modelled using an ODE framework, with the following RNA life-cycle stages represented as state variables in units of molecule counts: nascent RNA, early splicing RNA, late splicing RNA, free mRNA, translating mRNA, protein. Transitions between life-cycle stages, as well as all considered degradation processes, were described using first-order mass-action kinetics.

The model includes a negative feedback loop in which the gene product of *TNFAIP3* inhibits transcription of both genes. This inhibitory effect is implemented as a hyperbolic function, characterized by an inhibition constant. It scales the transcription rate between the maximum value in the absence of the inhibiting protein, a half-maximal value determined by the estimated inhibition constant, and the asymptotically approached zero at high protein abundance.

#### Mapping of state variables to model observables

To map the state variables of the model to observables that can be compared to the measured average molecule count in the responder cell population data, a time-weighted averaging approach was employed. This approach accounts for the fact that individual cells respond asynchronously, with response times distributed according to the response-time distribution estimated as described above.

At each measurement time point, the respective observable corresponds to the average molecule count of a population of cells following the specified kinetic model but initiating expression after heterogeneous response delays. This is realized by introducing two normalized leaky integrator ODEs per measured state variable, corresponding to the decay rates of the two components of the fitted delay time distribution. These auxiliary state variables compute exponentially weighted temporal averages of the corresponding model state variable, each characterized by a distinct decay constant given by the estimated decay constants of the gene specific delay distribution. The model observables are then defined as weighted sums of these auxiliary state variables, with weights given by the mixture weights of the delay distributions. This construction yields an observation function that represents the convolution of the model dynamics with the empirically inferred distribution of response delay times, while preserving a purely ODE-based model formulation. More detailed information is provided in the modelling supplement (**Supplementary Note**).

For state variables that map to splicing cluster spot counts, which correspond to more than one molecule, multiplicative scaling parameters (unit: spots/molecule) were introduced in the observation function. Those parameters convert from molecule counts of the state variables to spot counts in the observables and were estimated from the data together with the kinetic and error model parameters. The last time point (1440 min) was excluded from the observable data set as we observed that the model had difficulties in producing a good fit when this time point was included. Potentially there is a slow change in the cell state over this prolonged time which is not part of our model. We, thus, decided to model the response only up to the 600 min time point.

#### Model calibration

Model calibration was performed by estimating the model parameters (kinetic + scaling + error model) from the available average spot count data of the responder population. Parameter estimation was carried out using the Python package pyPESTO^67^, with the Python package AMICI^68^ used for numerical simulation of the ODE model. The parameter estimation problem was specified in the PEtab^69^ standard format. Parameters were estimated in logarithmic space, with individually specified search boundaries for each parameter. A multi-start local optimization strategy was employed to reduce the risk of convergence to local optima. The error model assumed additive Gaussian noise in the model observables. Uncertainty in the maximum likelihood parameter estimates was quantified using a profile likelihood approach as implemented in pyPESTO, providing confidence intervals and identifiability assessment for individual parameters.

#### Selection of model structure

To identify the model structure, a set of alternative model variants was constructed manually. In this context, *model structure* refers to the number of state variables, the biochemical reactions included, and the connectivity between model states. Each model variant therefore represents a distinct mechanistic hypothesis about the organization of the RNA life cycle. Each candidate model structure was calibrated to the data using the parameter estimation procedure described above. All parameters were assumed to be gene-specific, yielding maximum models in which no parameter sharing between genes was enforced. Model variants were compared using the Bayesian Information Criterion (BIC), which balances goodness-of-fit with model complexity. The final model structure was selected as the maximum model with the lowest BIC among all evaluated variants. This model was subsequently used as the reference structure for the identification of gene-specific parameters.

#### Identification of gene-specific parameters by model selection

To determine which model parameters differ between the two genes, a systematic model-selection procedure was performed using PEtab Select^70^. The approach constructs a hierarchy of nested models in which parameters are either shared between genes or assumed to be gene-specific, in a forward selection approach starting with the smallest model in terms of the number of parameters. In each iteration all model variants with one additional parameter are evaluated and the best model is used as the starting model for the next iteration. Each model variant was fitted to the data using the parameter estimation procedure described above. Model variants were evaluated using the Bayesian Information Criterion (BIC). Parameters were classified as gene-specific if allowing them to differ between genes substantially improved model fit relative to the increase in model complexity. This procedure identifies the minimal set of parameters that must be gene-specific to explain the observed differences in dynamics.

#### Simulating the effect of transcription inhibition on induction dynamics

Time courses of expression induction with and without Actinomycin D addition between 60 and 120 min were simulated by Monte-Carlo simulations, since the addition of the drug and the resulting changes in single cell transitions from the non-responder to the responder state can be more easily implemented in this framework compared to the auxiliary ODE system described above. We assumed the transcription rate to gradually decrease to 10% of its original value over an interval of 15 min after inhibitor addition, as Actinomycin D inhibition has been reported to be incomplete and could have delays depending on the cell type and the concentration used^71,72^. After inhibition cells were also not able any more to transition from the non-responder to the responder state. Single cell trajectories were averaged to obtain the whole population behavior, and the data was presented normalized to the un-inhibited 120 min time point for each gene. A more detailed description of the simulation approach can be found in **Supplementary Note**.

## Resource availability

### Material availability

DNA probe, hairpin and primer sequences used in this study are provided in **Supplementary Table S1** and can be obtained through commercial vendors as designed.

### Data and code availability

The data and code used for analysis have been deposited on Zenodo and GitHub, respectively, and will be made publicly available upon formal publication. In the interim, access is available upon request to the corresponding author.

## Acknowledgements

This work was supported by core funding from European Molecular Biology Laboratory. JT acknowledges the financial support from European Union’s Horizon 2020 research and innovation program under Marie Skłodowska Curie Actions in the EIPOD4 program (Grant agreement number 847543). We thank Peng Yin for early discussions of the method, Miki Ebisuya and Mitsuhiro Matsuda for discussions of potential applications and Melike Lakadamyali for comments on the manuscript draft. We thank Felix Schneider (EMBL GenTechDev OpenLab) for advice on image data analysis and Lawrence Landrock for help with the ProPER animation. We thank EMBL Data Science Center and EMBL Advanced Light Microscopy Facility for their valuable technical support.

## Author Contributions

J.T., V.P., J.Y.K. and S.K.S. - conceived the project.

J.T. J.Z., V.P. - experimental design, sample preparation and imaging data generation.

J.T. - data analysis and manuscript drafting

E.M.G. - implemented kinetic modeling with input from J.T. and S.K.S.

R.C.M, R.R., S.A., J.H. - contributed to data generation

K.H. - contributed to supervision.

S.K.S. - experimental design, data interpretation, manuscript drafting and supervision. All authors contributed to writing of the manuscript.

## Declaration of Interests

S.K.S. and J.Y.K are inventors on patent applications related to the methods described here. S.K.S. is a scientific co-founder and shareholder, and J.Y.K. is a co-founder, shareholder and employee of Digital Biology, Inc. S.K.S receives research funding from GSK and Leica Microsystems for projects unrelated to this work.

## Notes

### Summary of Updates

Minor text and title revision for increased clarity

