## Supplementary Information for "ProPER: Ligation-free in situ proximity extension enables multiplexed molecular-state analysis"

**Supplementary Information** includes:

**Supplementary Figure S1.** Transcript detection by intramolecular ProPER and negative controls.
**Supplementary Figure S2.** Controls and additional characterizations of protein-protein and RNA transcript detection by ProPER.
**Supplementary Figure S3.** Negative controls and additional quantifications for hairpin-primer screen.
**Supplementary Figure S4.** Proximity detection of transcripts to life-cycle markers and approaches to suppress background.
**Supplementary Figure S5.** Nascent RNA detection by metabolic labeling dependent-ProPER and comparison to splicing.
**Supplementary Figure S6.** Four-plex ProPER with negative controls and gene-resolved spot statistics.
**Supplementary Figure S7.** Image analysis and spot calling for four-channel transcript life-cycle ProPER.
**Supplementary Figure S8.** Co-staining of splicing activity by ProPER with nuclear speckles by immunofluorescence of speckle marker SON.
**Supplementary Figure S9.** Full time series of representative images of transcript life-cycle ProPER after TNF-α induction.
**Supplementary Figure S10.** Replicate level and single cell readouts of transcript life-cycle dynamics of two TNF-α-induced genes.
**Supplementary Figure S11.** Splicing activity along gene induction trajectories.
**Supplementary Figure S12.** Comparison of total transcript levels in cells with and without splicing activity at TS.
**Supplementary Figure S13.** Kinetic modeling of gene-specific expression steps.
**Supplementary Figure S14.** Perturbation of TNF-α response by transcription inhibition.

#### List of Supplementary Files

**Supplementary Movie.** Animation depicting the mechanism of ProPER (using *Blender 5.1.0*)

**Supplementary Table S1.** List of oligonucleotide sequences.
Tab 1: Probes for protein-protein ProPER
Tab 2: CBX5 primary probes
Tab 3: ICAM1 primary probes
Tab 4: TNFAIP3 primary probes
Tab 5: ICAM1 primary probes for concatemer production by in vitro PER
Tab 6: TNFAIP3 primary probes for concatemer production by in vitro PER
Tab 7: Docking probes for RNA life-cycle targets (U1, U2, 18S, azide-oligo for EU-click)
Tab 8: Branch probes
Tab 9: Sequence converters
Tab 10: ProPER hairpins
Tab 11: ProPER primers
Tab 12: Imager probes
Tab 13: Available hairpin-primer sequences for PER with validation status for ProPER

**Supplementary Table S2.** Probe combinations used in experiments.
Tab 1: CBX5 detection with low probe numbers
Tab 2: Complete set of negative controls for ProPER detection of ICAM1 transcript
Tab 3: Protein-protein ProPER for microtubule detection
Tab 4: ProPER comparison to smFISH (two-color detection)
Tab 5: Hairpin-primer pair screen
Tab 6: Optimization or ProPER RNA life-cycle staining (U1, U2, 18S)
Tab 7: ProPER licensing for ICAM1-18S detection
Tab 8: Multiplex RNA life-cycle ProPER (total, U1, U2, 18S)
Tab 9: Multiplex ProPER for nascent RNA (EU) plus U1 and SABER-FISH

**Supplementary Table S3.** Statistics of spot and image quantifications.
Tab 1: CBX5 detection with low probe numbers
Tab 2: Complete set of negative controls for ProPER detection of ICAM1 transcript
Tab 3: Comparison of detected microtubule fragment geometries after ridge detection
Tab 4: ProPER comparison to smFISH (2-color detection)
Tab 5: Hairpin-primer sequence pair screen
Tab 6: ProPER licensing for ICAM1-18S detection
Tab 7: 18S hairpin docking probe comparison
Tab 8: Overlap of EU and U1 ProPER signal at TS (cluster spots)
Tab 9: Fourplex RNA life cycle with negative controls
Tab 10: Distance of splicing transcripts to next nuclear speckle
Tab 11: Average quantities of RNA life-cycle readouts over TNF time course
Tab 12: Total transcript counts of single cells stratified by presence of cluster splice sites
Tab 13: Kinetic modeling: Bayesian information criterion for model discrimination
Tab 14: RNA life-cycle stages after transcription inhibition with Actinomycin D

**Supplementary Note.** Detailed description of kinetic model and references

### Supplementary Figures

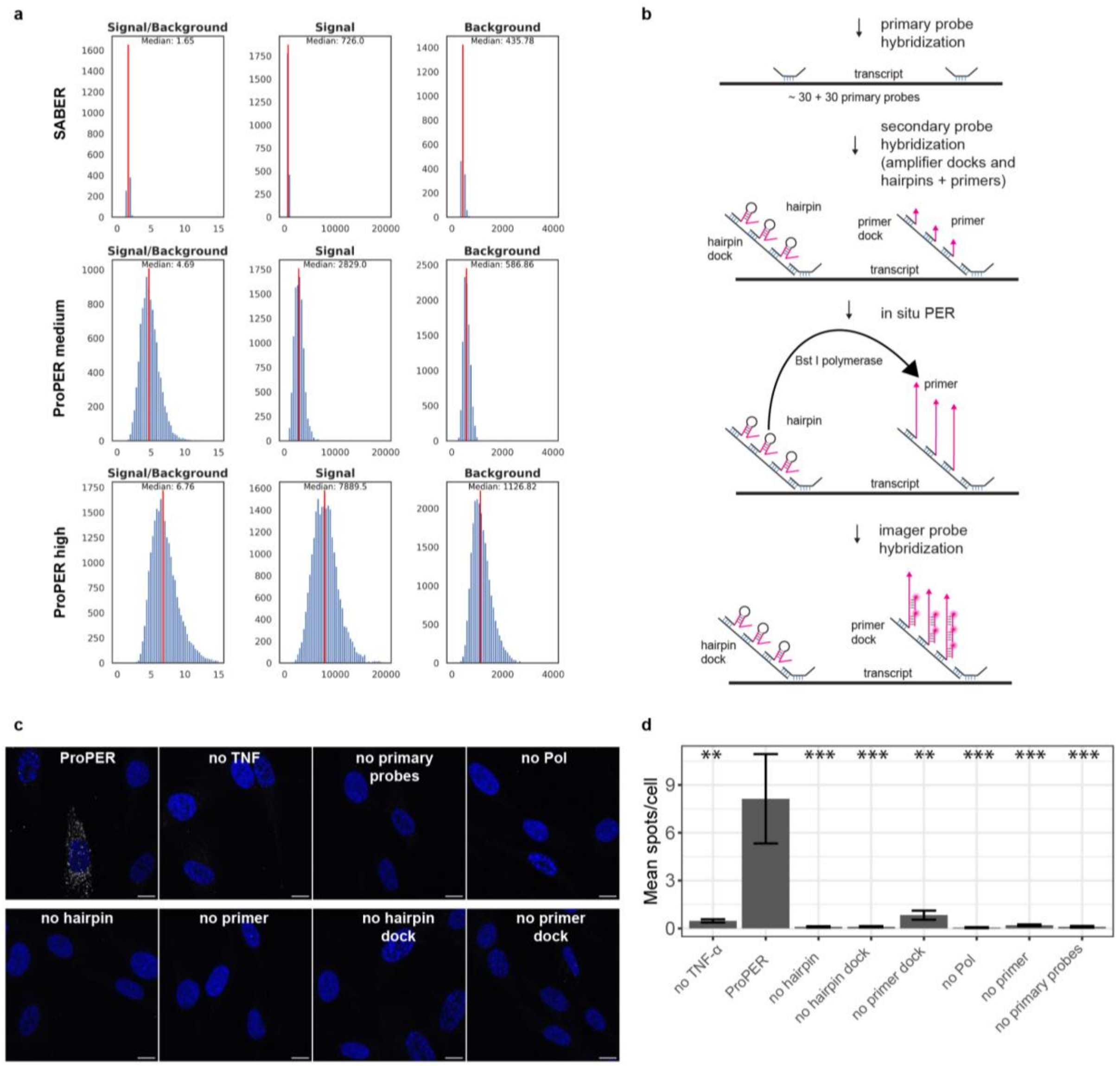


**Supplementary Figure S1. Transcript detection by intramolecular ProPER and negative controls.** **(a)** Histograms represent signal/background ratio, signal and background for *CBX5* transcript spots in HeLa cells for SABER-FISH, ProPER medium and high (**Fig. 1c-f**). For calculation of signal/background ratio are provided in **Methods**. Red lines represent median values for each distribution. **b,** Steps of an (intramolecular) ProPER experiment. After primary probe hybridization, branches, hairpins and primers are added together during secondary hybridization. In situ PER creates concatemers from primers that are in sufficient proximity (coincidence detection on the same transcript), which is then visualized by fluorescent probes added during imager hybridization. **(c-d)** ProPER detection of ICAM1 transcripts 120 min after TNF-α treatment in NHLF cells and negative controls using primary probe numbers that are not limiting. **(c)** ProPER stained transcripts in responding cells, as can be seen in representative images (max projection), while only infrequent spots were observed, when any of the required probes, treatment or the polymerase (no Pol) were left out. Scale bars, 10 µm. **(d)** Quantification of average spot numbers per cell confirmed this. Error bars +/- s.e.m., Mann-Whitney-Wilcoxon test of negative controls against full ProPER, two-sided, ** $p<0.01$, *** $p<0.001$ (exact p-values in **Table S3**). n=59-85 cells per condition.

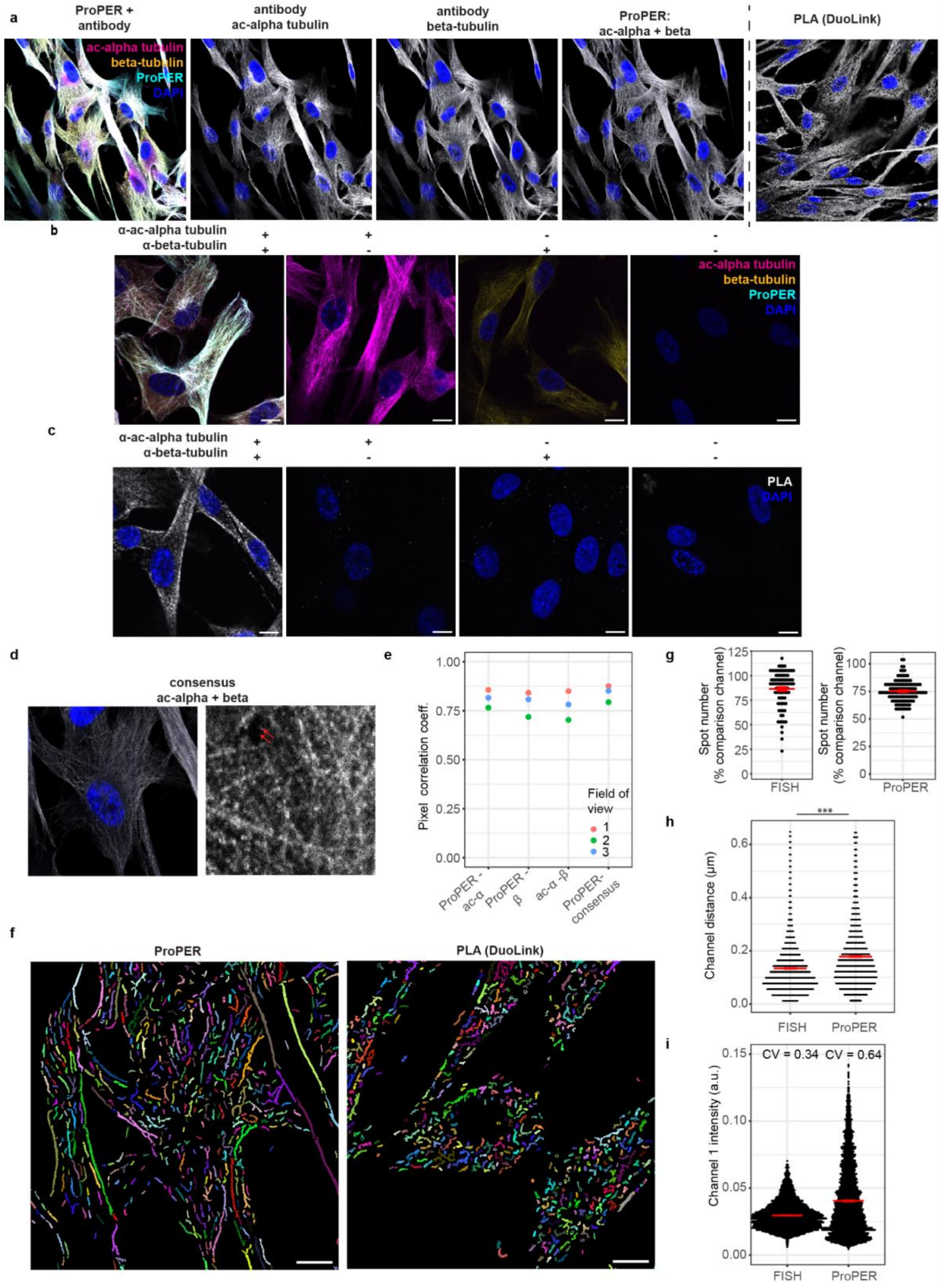


**Supplementary Figure S2. Controls and additional characterizations of protein-protein and RNA transcript detection by ProPER. (a)** Full field of view for ac-alpha- and beta-tubulin individual stainings, ProPER and PLA (commercial DuoLink assay). **(b)** Negative controls for tubulin ProPER leaving out either one or both primary antibodies. Note that due to the direct binding site for imagers on the antibody bound dock, the individual antibody stainings can still be visualized, when the other antibody is absent. **(c)** Negative controls for tubulin PLA leaving out either one or both primary antibodies. **(d)** Consensus image of the two antibody signals in the ProPER experiment created by minimum projection along the channel axis (minimum of individual antibody stainings of ac-alpha- and beta-tubulin). Only pixels with strong signal in both antibody channels will have high intensity. This represents a good reference for the ProPER signal. **(e)** Pixel-wise intensity correlations were high across all readouts and highest between ProPER and the consensus image. **(f)** Large fields of view of Frangi-segmented tubules from ProPER (left) or PLA (right) images. Individual fragments were false-colored. Tubules from PLA are visibly shorter and less aligned. Scale bars, 10 µm. **(g-i)** Two-color spot properties of data in **Fig. 2g-j**. **(g)** Distribution of distances of spot center coordinates between channel 1 and 2 for spots present in both channels. Mann-Whitney-Wilcoxon test, two-sided, *** $p<0.001$ (exact p-values in **Supplementary Table S3**). **(h)** Distribution of channel 1 spot intensities in the cytoplasm (ProPER or 2-color-FISH) in arbitrary fluorescence units for spots present in both channels. In the nucleus there is a small population of large and very bright spots, so that nuclear spots were excluded from this analysis. CV reports the coefficient of variation for the distributions and indicates a higher variability of spot intensities for ProPER compared to smFISH. **(i)** The relative number of spots for FISH or ProPER was calculated in each cell compared to the channel with higher number of spots (for ProPER, that comparison channel is ch2, FISH; for dual-color-FISH, it is ch1) to determine channel agreement. Only cells with a minimum number of spots in each channel were considered. n=92-128 cells per condition. Red bars indicate the mean, red error bars +/- s.e.m.

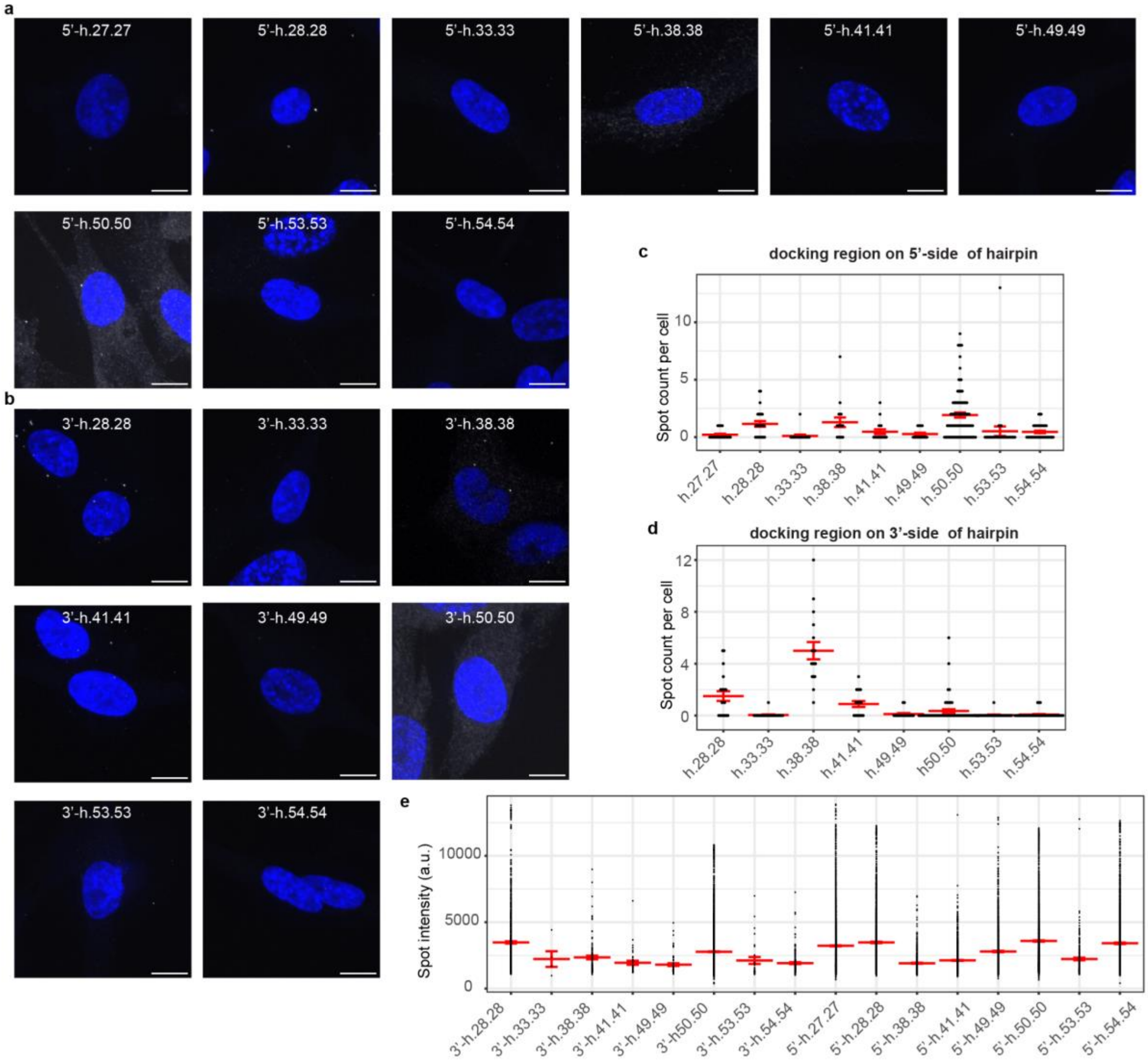


**Supplementary Figure S3. Negative controls and additional quantifications for hairpin-primer screen. (a-b)** Representative images of *ICAM1*-ProPER signal for untreated cells, in which no gene induction is expected for hairpins with docking site on the **(a)** 5'- and **(b)** 3' end. Scale bars 10 µm. **(c-d)** Average spot counts per cell in negative controls for each sequence pair were quantified for hairpins docking on their **(c)** 5'- side, or **(d)** on 3'-side. n=17-96 cells. **(e)** Distribution of spot intensities in arbitrary fluorescence units (a.u.) for all hairpin-primer pairs for TNF-α induced cells (**Fig. 3**). Pairs for which no spots were detected are left out. Red bars indicate the mean. Error bars +/- s.e.m.

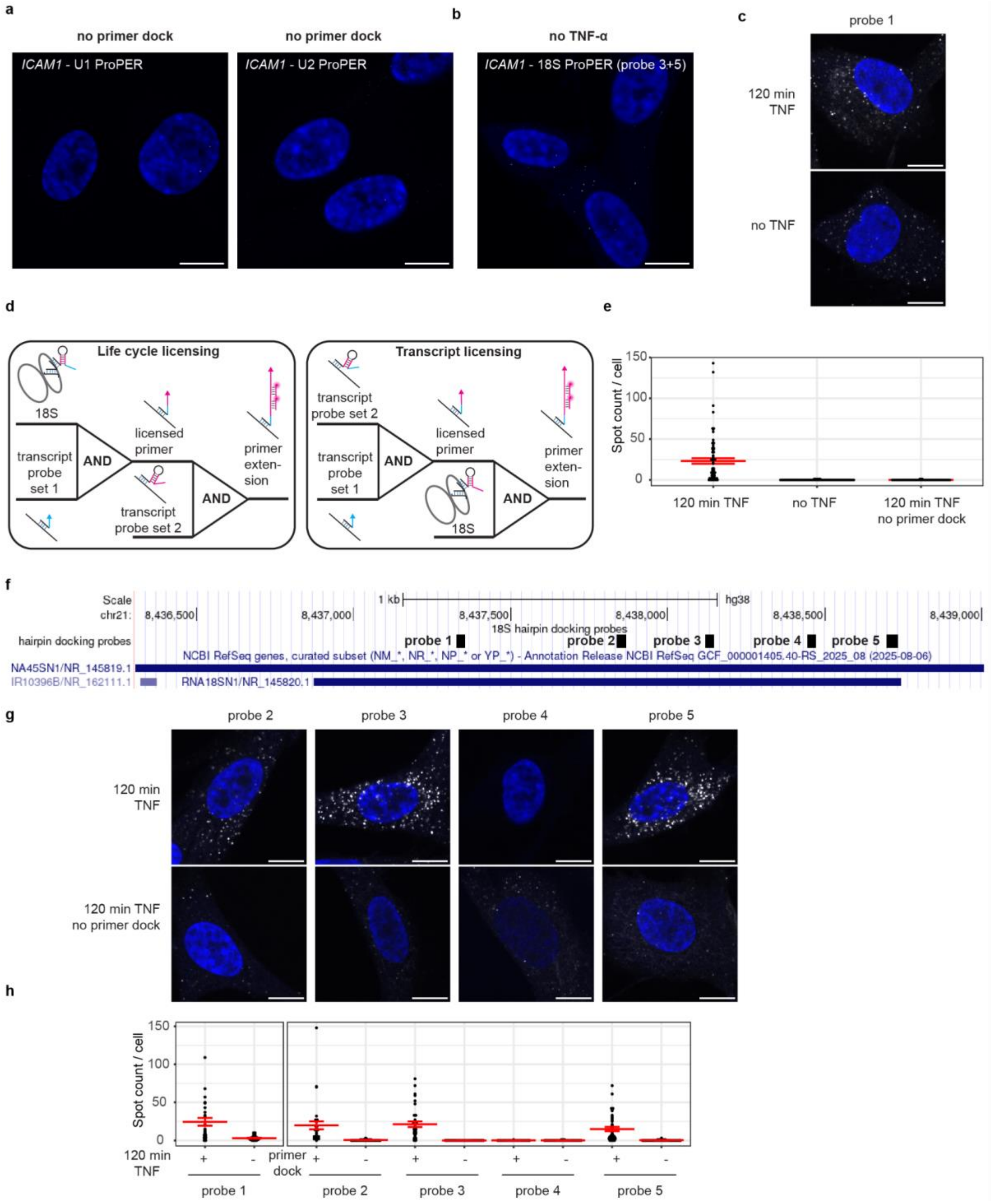


**Supplementary Figure S4. Proximity detection of transcripts to life-cycle markers and approaches to suppress background. (a)** Representative images for negative controls of U1- and U2-ProPER for *ICAM1* 120 min after TNF-α treatment without primer dock. **(b)** Representative image for negative controls of 18S-ProPER without TNF-α treatment (using 18S probes 3+5). **(c)** Representative image for 18S-ProPER using suboptimal 18S probes 1. **(d)** Licensing strategy to create a logic gate requiring first the proximity of the life-cycle marker (18S) and target transcript and then proximity of the reaction product with an additional probe set on the transcript. For transcript licensing the order of detecting the proximities is swapped. **(e)** Spot quantification for life-cycle licensing with relaxed intensity thresholds recovers higher spot numbers (compared to the more stringent analysis in **Fig. 4h**) without increasing false-detections in the negative controls (uninduced cells or leaving out the primer dock). **(f)** Selection of life-cycle target probe(s) for 18S ProPER. The Genome Browser track shows positions of candidate probes for hairpin docking on the 18S gene (18SN1) indicated by black boxes. **(g)** Representative images for 18S-ICAM1 ProPER using single candidate life-cycle target probes on NHLF cells 120 min after TNF-α treatment, left untreated or leaving out the primer dock during ProPER. **(h)** Spot number distribution per cell showed probes 3 and 5 had low background counts, while preserving similar average counts in the induced full-ProPER condition compared to probe 1. Probe 1 corresponds to the first probe that was tested and for which higher background levels were observed (panel **c**). Scale bars 10 µm. Red bars indicate the mean. Error bars +/- s.e.m.

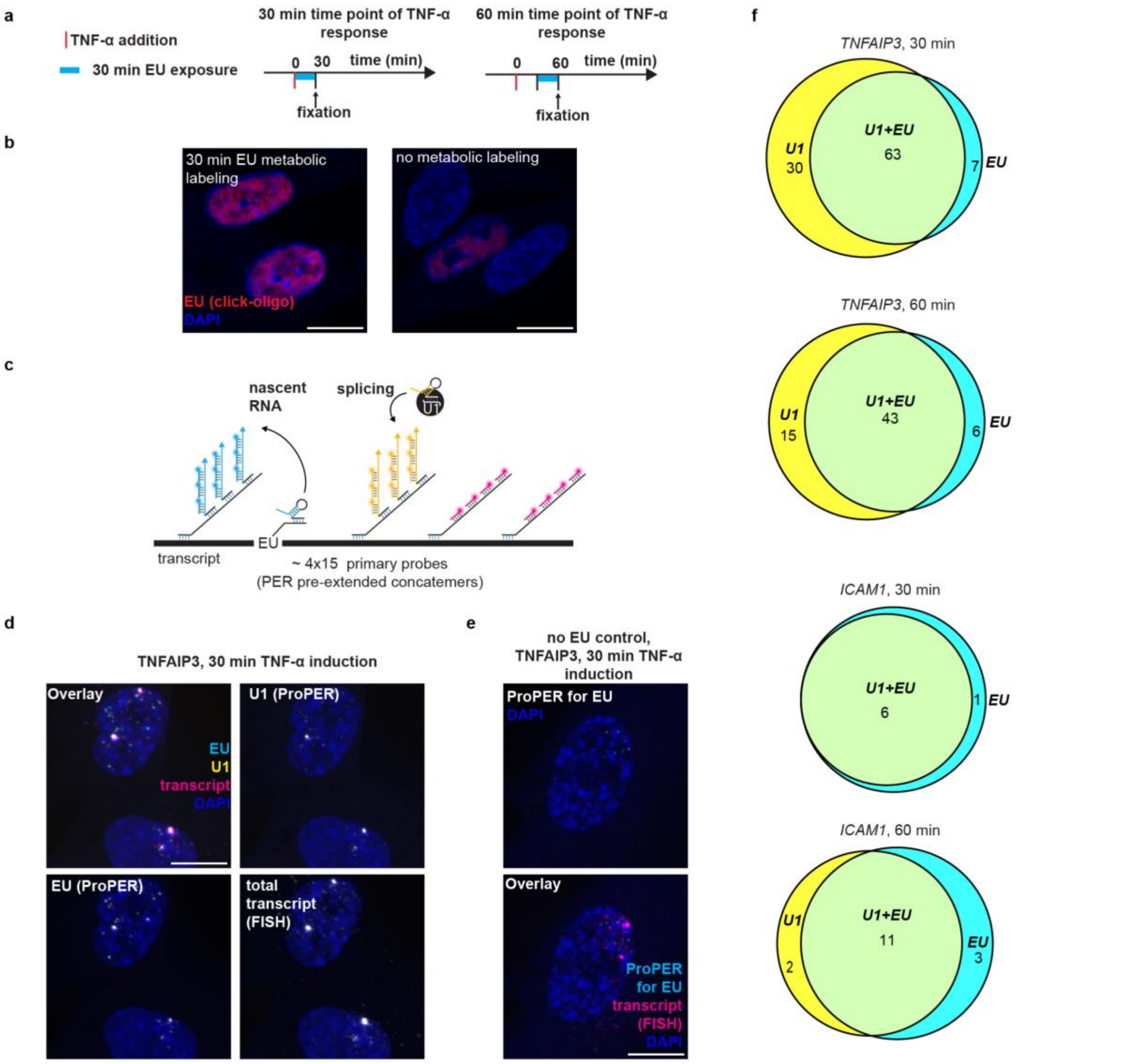


**Supplementary Figure S5. Nascent RNA detection by metabolic labeling dependent-ProPER and comparison to splicing. (a)** Metabolic labeling of nascent RNA by 5-ethynyl uridine (EU) pulse. Experimental time line: Cells were exposed to TNF-α at time 0, for 30 or 60 min, respectively for the two time points. In each case, to label nascent RNA, 1 mM EU was added to the media 30 min before fixation and sample collection. **(b)** Representative images showing anchoring of oligo-DNA docking sites to nascent RNA by click reaction of azide-oligo with EU. Clicked oligo is visualized by hybridization of fluorescent imager oligos to the docking site. Some background binding in cells without metabolic labeling can be seen (bottom), though this background from the click reaction is much weaker compared to the case with EU supplementation (top). **(c)** Multiplex ProPER scheme for detecting EU metabolic labeling and splicing (by U1 proximity) together with total transcript detection by SABER-FISH. For this application, pre-amplified concatemers were used as primary probes. For EU-proximity detection, the hairpin was bound to the docking oligo attached to EU via click reaction. **(d)** Representative image of two-plex ProPER + SABER-FISH detection of TNFAIP3 30 min after TNF-α treatment. The channel overlay is shown as a multicolor image (EU-ProPER: ATTO 565, cyan; U1-ProPER: ATTO 488, yellow; FISH for total TNFAIP3 transcript: Alexa Fluor 647, magenta) and individual channels in gray scale with DAPI in dark bluel. EU-ProPER displays a similar pattern as U1-ProPER including dispersed and cluster spots. **(e)** The negative control of 2-plex-ProPER plus SABER-FISH without EU pulse shows no detectable ProPER signal (EU-ProPER channel in gray scale, top). In the multicolor overlay of the same image (bottom) total transcripts measured by FISH can be seen, but not their EU-ProPER signal (EU-ProPER: ATTO 565, cyan; FISH for total TNFAIP3 transcript: Alexa Fluor 647, magenta; DAPI: dark blue). **(f)** Venn diagrams showing cluster spots present in U1- (yellow) and EU-channel (blue) or in both (green) after 30 or 60 min of TNF-α treatment. Most cluster spots are detected both in the EU- and in the U1-ProPER channel. Scale bars 10 µm.

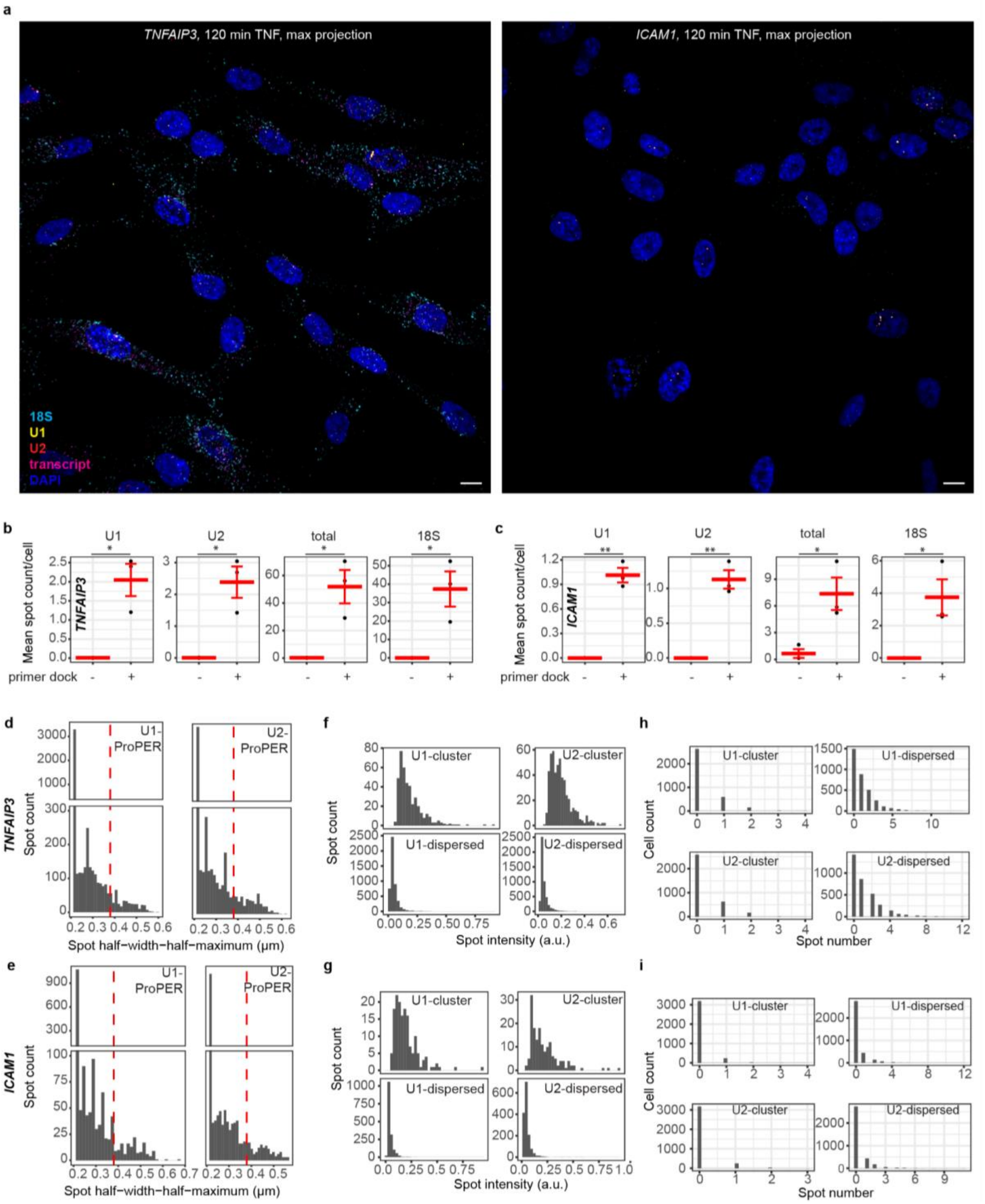

**Supplementary Figure S6. 4-plex ProPER with negative controls and gene-resolved spot statistics. (a)** Full FOV for the images shown in **Fig. 5c** (max. projection of 3 z-slices). Scale bars 10 µm. (**b-c)** Average spot number per cell for each ProPER readout for **(b)** *TNFAIP3* and **(c)** *ICAM1*. Red bars indicate the mean of n=3 replicates (single cell data shown in **Fig. 5e-f**). Error bars +/- s.e.m. Welch t-test, one-sided, * p<0.05, ** p<0.01, *** p<0.001 (exact p-values in **Supplementary Table S3**). **(d-i)** Distributions of U1/U2 spot properties corresponding to **Fig. 5h-j** separated by gene. **(d-e)** Spot radius measured by half-width-half-maxium for **(d)** *TNFAIP3,* and **(e)** *ICAM1*. The red dashed line indicates the classification cut-off for dispersed/cluster spots (see **Methods**). **(f-g)** Spot intensities (in arbitrary fluorescence units, a.u.) for **(f)** *TNFAIP3,* and **(g)** *ICAM1*. **(h-i)** Detected number of spots of the respective spot classes (U1/U2, cluster/dispersed) per cell for **(h)** *TNFAIP3,* and **(i)** *ICAM1*.

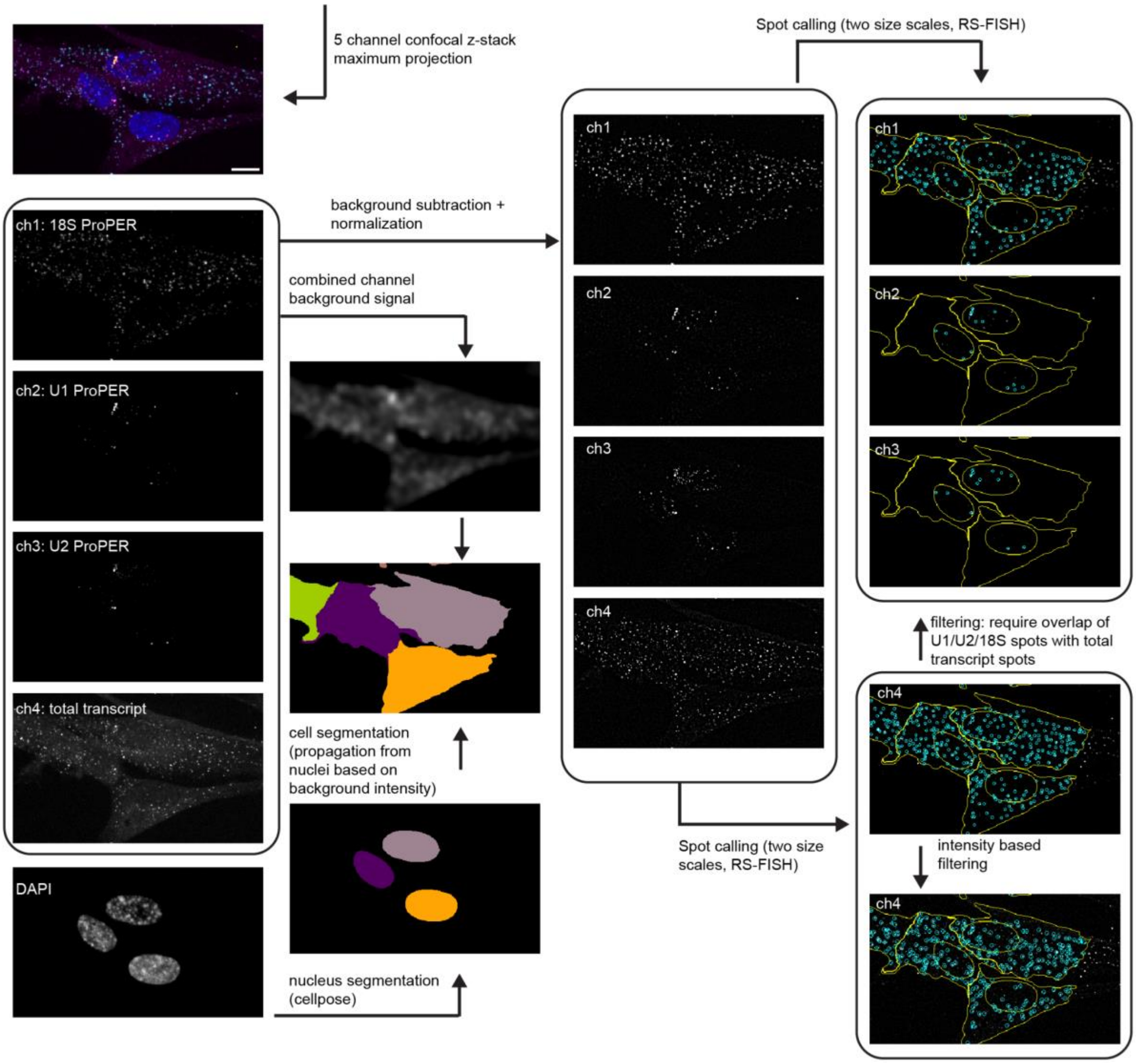


**Supplementary Figure S7. Image analysis and spot calling for four-channel transcript life-cycle ProPER.** The input data consists of confocal z-stacks with four ProPER channels plus DAPI. Hyperstacks are maximum projected and split into the individual channels. The DAPI channel is used for nucleus segmentation by *cellpose*. Background signal in blurred combined ProPER channels is used for cell segmentation using nucleus masks as seed points for mask propagation. ProPER channels are prepared for spot calling by background subtraction, artifact removal and intensity normalization. Spots are then called in each individual channel based on radial symmetry using the RS-FISH plugin in ImageJ. The total transcript channel is used as a reference to filter spots in the other three ProPER channels. Overlap with a total-transcript spot (channel 4) within a 3-pixel radius is required for channels 1, 2 and 3. Afterwards, total transcript spots in channel 4 are filtered by applying a threshold derived from their intensity distribution. The coordinates of each spot allow to assign the spots to a cell and compartment (nucleus or cytoplasm). For additional information on image and spot processing see **Methods**.

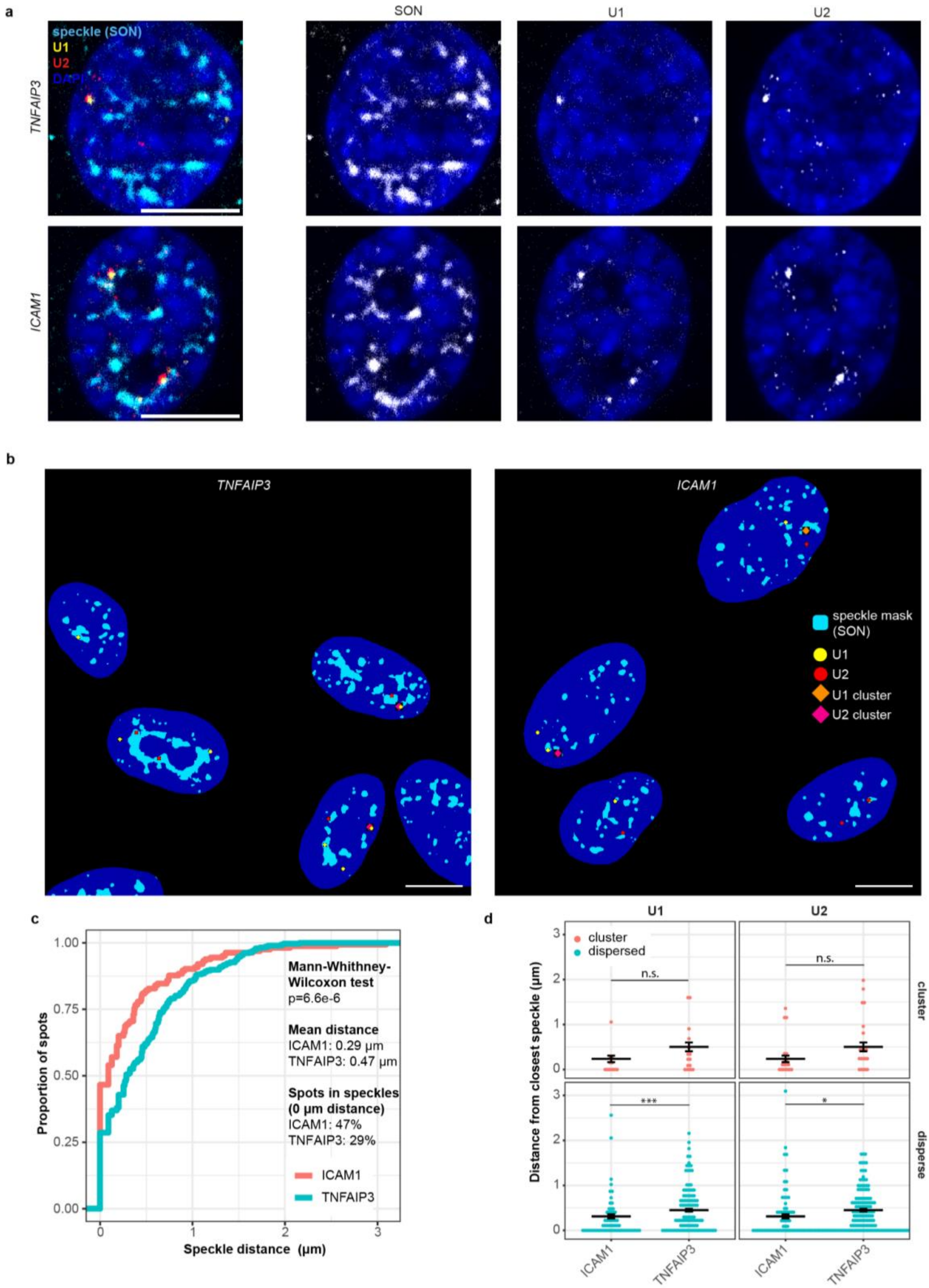


**Supplementary Figure S8. Co-staining of splicing activity by ProPER with nuclear speckles by immunofluorescence of speckle marker SON. (a)** Antibody staining for the speckle marker SON after ProPER for U1 and U2. Representative max. projections of three z-slices. Splicing transcripts only partially overlap with speckles and can be touching or even distant to speckles. **(b)** Representative single z-planes of segmentation masks of nuclei, speckles, U1 and U2 spots (classified as cluster spots if they are above a size threshold). **(c)** Distribution (cumulative distribution function) of distances of splicing spots from the next speckle. **(d)** Distribution of distances of U1- and U2-spots from the next nuclear speckle. If the spot is inside a speckle this distance is zero. Mann-Whitney-Wilcoxon test, two-sided, n.s. not significant, * $p<0.05$,** $p<0.01$ (exact p-values in **Supplementary Table S3**). n=21-23 cells per condition. Error bars s.e.m.

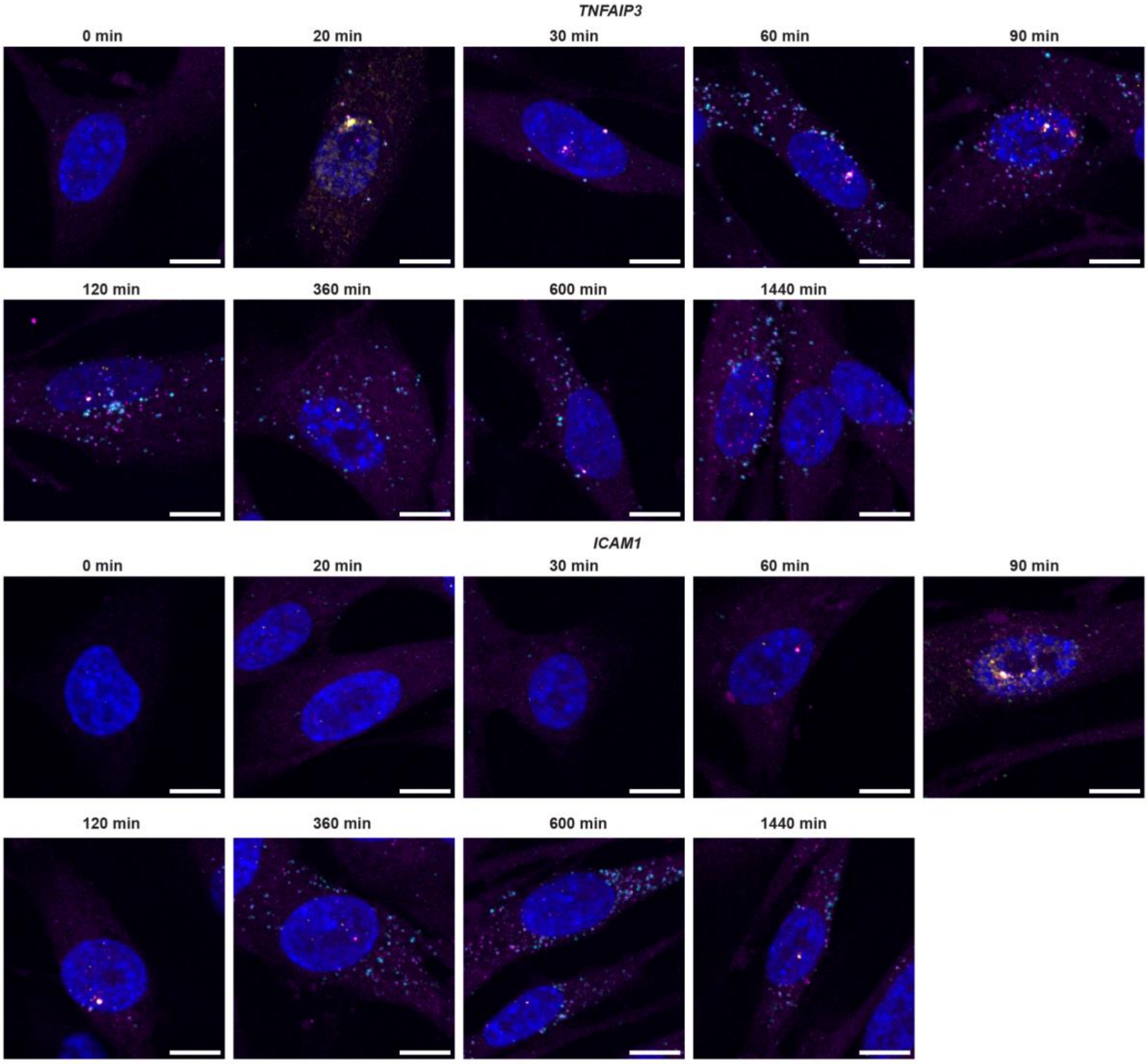


**Supplementary Figure S9. Full time series of representative images of transcript life-cycle ProPER after TNF-α induction.** Full image series of **Fig. 6a**. Multicolor images of NHLF cells after fourplex ProPER for U1-, U2-, 18S-proximity and transcript detection for *TNFAIP3* (top) or *ICAM1* (bottom). Color assignment as in **Fig. 6a** (magenta: total transcript, yellow: U1, red: U2, cyan: 18S). Maximum projection of three z-planes. Scale bars, 10 µm.

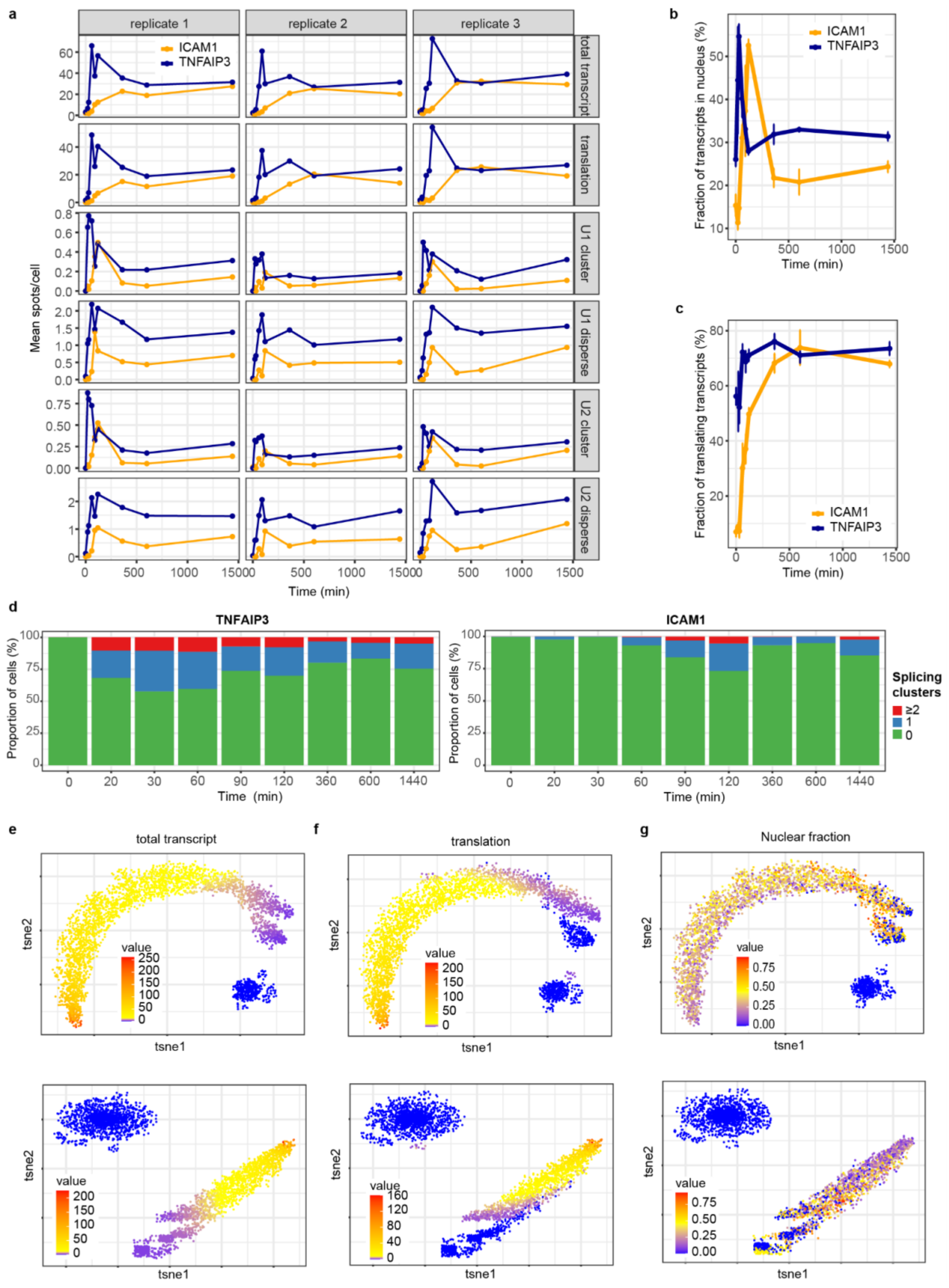


**Supplementary Figure S10. Replicate level and single cell readouts of transcript life-cycle dynamics of two TNF-α-induced genes. (a)** Average spot count time courses for individual replicates underlying the averaged time course shown in **Fig. 6b** for *ICAM1* (orange) and *TNFAIP3* (dark blue). 91-162 cells per time point and replicate. (**b**) Dynamics of the fraction of transcripts in the nucleus. This was calculated for each cell as the number of spots in the total-transcript channel inside the nucleus divided by the sum of those in nucleus and cytoplasm. (**c)** Dynamics of the fraction of translating transcripts. Calculated for each cell as the ratio of translating and total transcript. Error bars show +/- s.e.m. from n=3 replicates. (**d)** Proportion of cells having either 0, 1 or 2 and more splicing cluster sites for each time point. (**e-g)** Single cell representation by t-SNE from **Fig. 6d** showing **(e)** total transcript counts, **(f)** translating transcript counts and **(g)** nuclear fraction of transcripts for *TNFAIP3* (top) and *ICAM1* (bottom).

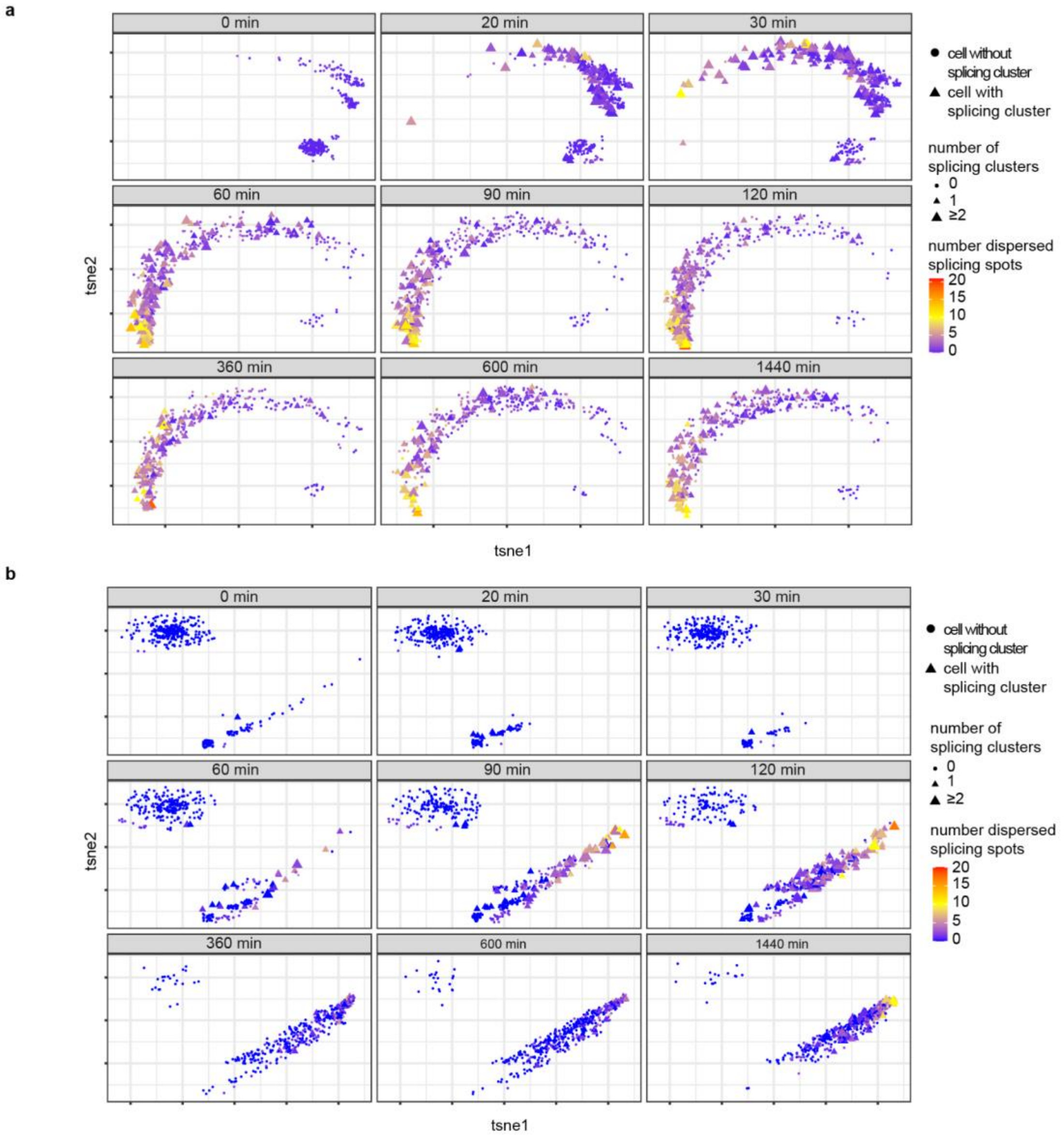


**Supplementary Figure S11. Splicing activity along gene induction trajectories. (a)** *TNFAIP3* and **(b)** *ICAM1* t-SNE plots corresponding to **Fig. 6d** split by time point and indicating the number of dispersed splicing spots (by color) and number of cluster splicing spots (by symbol). U1 and U2 spots were pooled into a single class of "splicing" spots and were classified as cluster if the spot was a cluster spot in either one or both channels. High splicing activity is observed in the leading front of the trajectory (parts of the trajectory that are more densely populated at later time points).

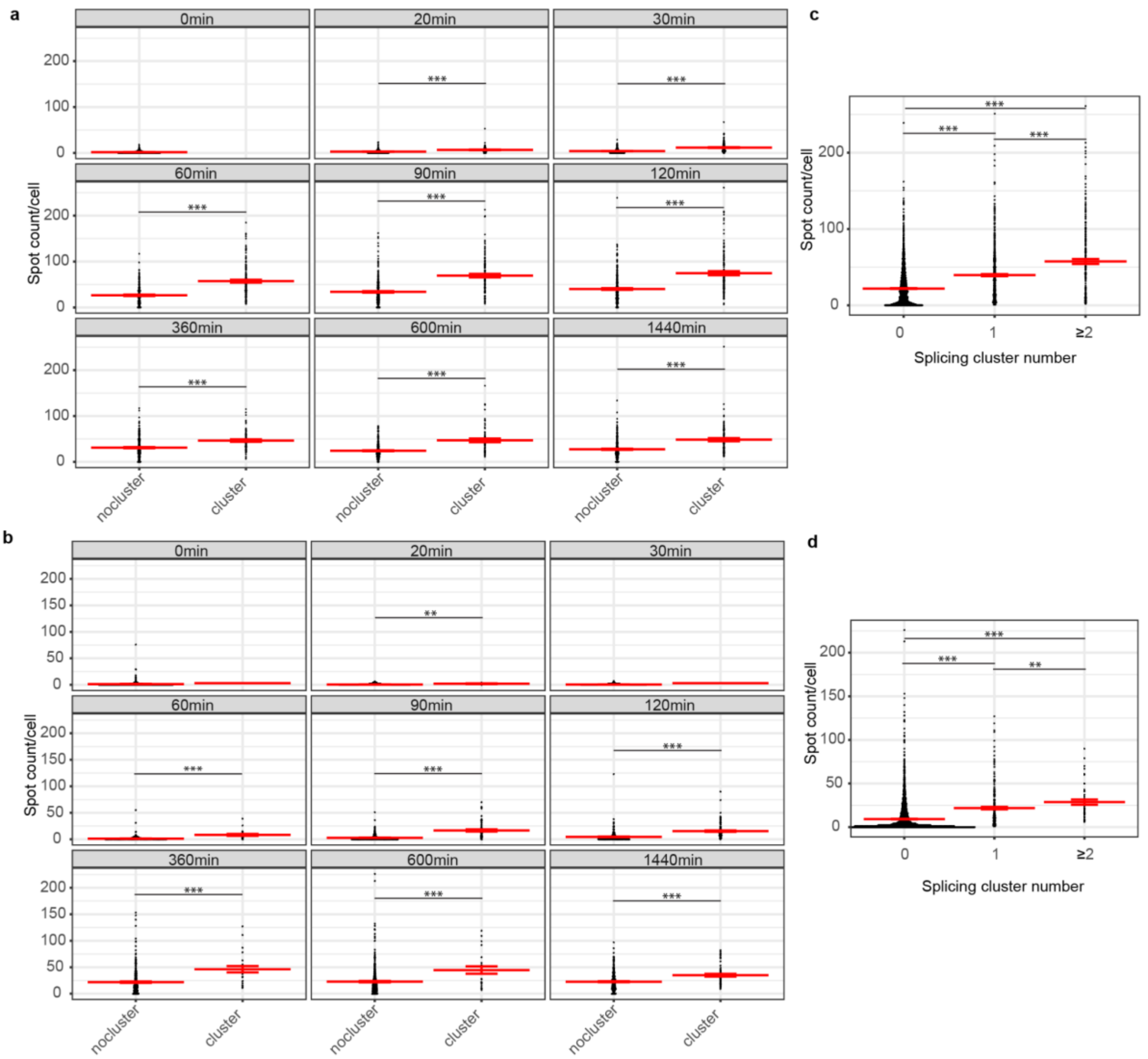


**Supplementary Figure S12. Comparison of total transcript levels in cells with and without splicing activity at TS.** For each time point, cells were stratified into two groups based on the presence or absence of at least one cluster splicing spot (see **Supplementary Fig. S11)**. The distribution of total transcript counts per cell are shown for (**a)** *TNFAIP3* and (**b)** *ICAM1*. (**c-d)** The same data is shown pooled from all of the time points and resolved by the number of splicing cluster sites for **(c)** *TNFAIP3,* and **(d)** *ICAM1*. Mean indicated by red line. Error bars, +/- s.e.m. Mann-Whitney-Wilcoxon test, two-sided, ** $p<0.01$, *** $p<0.001$ (exact p-values in **Supplementary Table S3**).

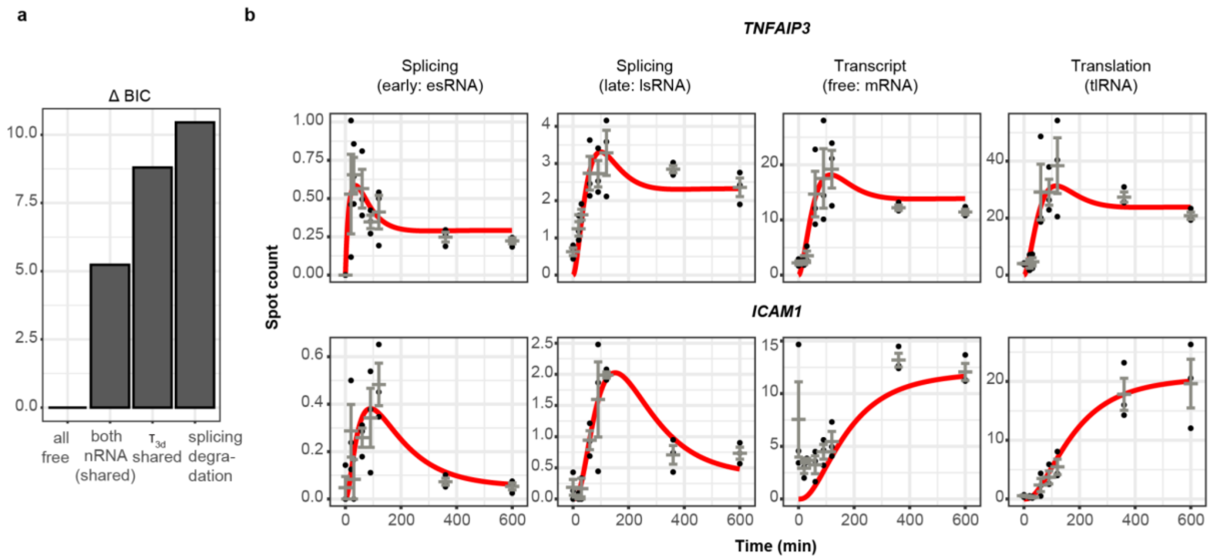


**Supplementary Figure S13. Kinetic modeling of gene-specific expression steps. (a)** Model pre-selection for early modeling decisions. Three scenarios were tested against the base model (**Fig. 7a**) in which all parameters of interest are allowed to take on gene-specific values ("all free"): in the "both nRNA (shared)" case, not only ICAM1, but also TNFAIP3 has the nRNA state. In "$\tau_{3d}$ shared" RNA degradation of both genes has the same rate constant. In "splicing degradation" splicing transcripts in the tlRNA state can be degraded. (For more details see the **Methods** and **Supplementary Note**). The goodness of fit (taking the number of free parameters into account) is reported by the difference of the Bayesian information criterion to the best performing model (ΔBIC). **(b)** Model fits of best model (**Fig. 7b-c**) of the responder-only cell population. The whole-population plots in **Fig. 7c** are derived from this by multiplying with the respective responder rates for each time point and gene. Black points, data points; red line, model fit. Error bars: s.e.m.

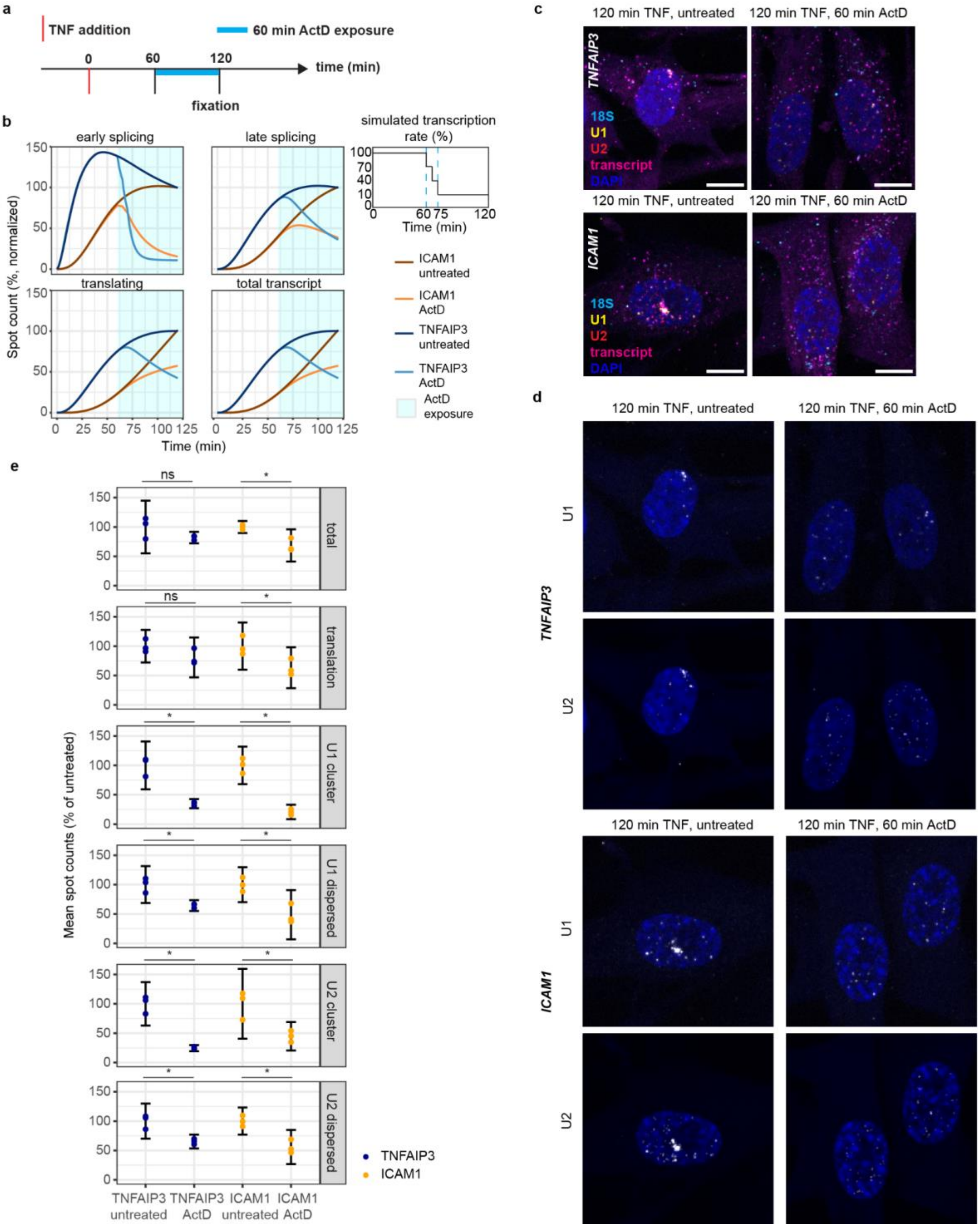


**Supplementary Figure S14. Perturbation of TNF-α response by transcription inhibition. (a)** Timeline of transcription inhibition experiment. 60 min after TNF-α exposure Actinomycin D (0.5 µg/ml) was added to inhibit transcription and cells were fixed at time point 120 min. **(b)** Simulation of transcript life-cycle stage trajectories with and without inhibition of transcription by Actionomycin D (ActD, cyan shaded area) between 60 and 120 min. Trajectories were calculated using the selected model and its parameters from **Fig. 7d**. In the simulation the transcription rate was reduced from 100% to 10% in two steps between 60 and 75 min (schematic, right). For details see **Methods** and **Supplementary Note**. Trajectories are shown normalized to the untreated 120 min time point. **(c)** Exemplary multicolor images of fourplex RNA life-cycle ProPER for U1-, U2-, 18S-proximity and transcript detection for *TNFAIP3* or *ICAM1* with and without transcription inhibition. Scale bars, 10 µm. **(d)** Individual channels of cells shown in **c**. **(e)** Average spot counts of life-cycle resolved transcripts per cell relative to the average value of the untreated condition (at 120 min). There was a reduction of nearly all RNA life-cycle stages, but the most prominent reductions were for U1 and U2 cluster spots. Error bars, 95% confidence intervals from n=3 replicates. Two-sided Welch t-test, ns not significant, * $p < 0.05$, (exact p-values in **Supplementary Table S3**). 515-559 cells per condition (from n=3 replicates combined).
