## Supplementary Note for "ProPER: Ligation-free in situ proximity extension enables multiplexed molecular-state analysis"

### 1 Parametric quantification of the response delay time

The time delay between TNF$\alpha$ stimulation and the onset of TNF$\alpha$-induced gene expression was quantified at the population level using single-cell response delay time information. Individual cells respond with a delay time on which partial information is available for each cell. To quantify this delay time on the population level, cells were assumed to respond after a stochastic delay $\tau$ following TNF$\alpha$ stimulation, with response delay times drawn independently from a parametric population-level distribution.

To flexibly capture heterogeneity in response timing, while taking advantage of the advantageous properties of the exponential distribution (see below), the delay time distribution was modeled as a finite mixture of exponential distributions. Mixture models with one, two, three, and four exponential components were evaluated. The optimal number of mixture components was selected based on the Bayesian Information Criterion (BIC), which balances goodness-of-fit against model complexity. Based on this comparison, a two-component exponential mixture was selected as the most parsimonious model explaining the observed response delay time data.

#### 1.1 Parametric form of the response delay time distribution

For the selected two-component exponential mixture model, the probability density function (pdf) of the response delay time $\tau$ is given by

$$p(\tau) = w_1 \lambda_1 e^{-\lambda_1 \tau} + w_2 \lambda_2 e^{-\lambda_2 \tau}, \tag{1}$$

where $\lambda_1$ and $\lambda_2$ denote the rate parameters of the individual exponential components, and $w_1$ and $w_2$ are the corresponding mixture weights, subject to the constraints

$$w_1 \geq 0, \quad w_2 \geq 0, \quad w_1 + w_2 = 1. \tag{2}$$

The corresponding cumulative distribution function (cdf) of the mixture distribution, representing the probability for a cell to have responded by time $t$ and also the fraction of cells that have responded by time $t$, is given by

$$F(t) = \Pr(\tau \leq t) = w_1 \left(1 - e^{-\lambda_1 t}\right) + w_2 \left(1 - e^{-\lambda_2 t}\right). \tag{3}$$

With this parameterization, the fraction of cells in the population that have responded by any time $t$ is given by the cumulative distribution function $F(t)$. In addition, the underlying delay-time distribution $p(\tau)$ fully specifies the distribution of response delays among responding cells. Together, $F(t)$ and $p(\tau)$ provide a quantitative description of the temporal heterogeneity in gene induction across the cell population. This information is essential for interpreting population-averaged RNA measurements within the mechanistic modeling framework employed in this study, as it enables explicit accounting for asynchronous response timing when analysing the dynamics of the RNA life cycle, as described in Section 2.

#### 1.2 Censoring structure of the single-cell response delay time data

For each individual cell $i$ in the dataset, the presence of TNF$\alpha$-induced gene expression was assessed at a single experimental time point $t_m$ following TNF$\alpha$ stimulation. The information obtained at this time point provides only partial insight into the cell-specific response delay, as it reveals whether gene expression had occurred by time $t_m$, but not the exact response delay time $\tau_i$. Consequently, the precise value of $\tau_i$ is not directly observed and the response delay time data are subject to censoring, giving rise to two types of observations:

- **Interval-censored observations:** For cells in which TNF$\alpha$-induced gene expression was detected at time $t_m$ (responder), the true response delay $\tau_i$ is known to lie within the interval $[0, t_m]$. These observations therefore correspond to interval-censored data.
- **Right-censored observations:** For cells in which no TNF$\alpha$-induced gene expression was detected at time $t_m$ (non responder), the true response delay $\tau_i$ is assumed to exceed $t_m$. These observations therefore correspond to right-censored data.

Response-time information from all individual cells (responders + non responders) across all measurement time points was incorporated jointly into the likelihood function used to estimate the unknown parameters in equation 3 as described in the next subsection.

##### 1.3 Likelihood formulation for mixture distribution estimation

The parameters of the delay distribution were estimated using a maximum likelihood approach that explicitly accounts for the censoring structure of the data.

For an interval-censored observation at time $t_m$, corresponding to a cell that has responded by $t_m$, the likelihood contribution is given by the probability that the response delay $\tau_i$ lies in the interval $[0, t_m]$, which is equal to the cumulative distribution function evaluated at $t_m$:

$$L_i = \Pr(\tau_i \leq t_m) = F(t_m). \tag{4}$$

For a right-censored observation at time $t_m$, corresponding to a cell that has not responded by $t_m$, the likelihood contribution is given by the probability that the response delay exceeds $t_m$:

$$L_i = \Pr(\tau_i > t_m) = 1 - F(t_m). \tag{5}$$

Assuming independence of response delays across cells, the full likelihood is given by

$$\mathcal{L}(\theta) = \prod_{i=1}^{N} L_i, \tag{6}$$

where $\theta = \{w_1, w_2, \lambda_1, \lambda_2\}$ denotes the set of model parameters and $N$ is he total number of cells. Parameter estimation was performed by maximum likelihood, which corresponds to finding the parameter vector

$$\theta = \{\lambda_1, \lambda_2, w_1, w_2\}$$

that maximizes the likelihood function:

$$\hat{\theta} = \arg\max_{\theta} \mathcal{L}(\theta). \tag{7}$$

Equivalently, this can be formulated as minimizing the negative log-likelihood:

$$\hat{\theta} = \arg\min_{\theta} \big(-\log \mathcal{L}(\theta)\big) = \arg\min_{\theta} \ell(\theta), \tag{8}$$

where $\ell(\theta) = -\log \mathcal{L}(\theta)$.

##### 1.4 Mixture model fitting and selection

All candidate mixture models were fitted by maximum likelihood estimation of the mixture weights and exponential rate parameters. Model comparison across different numbers of mixture components was performed using the Bayesian Information Criterion (BIC)

$$\mathrm{BIC} = 2\,\ell_{\min} + k \log N, \tag{9}$$

where $\ell_{\min}$ denotes the minimum negative log-likelihood achieved by the model, $k$ is the number of free parameters, and $N$ is the number of data points. The BIC balances model fit and complexity by penalizing the number of estimated parameters, thereby favoring parsimonious models that adequately explain the data. Competing model hypotheses were ranked according to their BIC values, with lower BIC indicating stronger support by the data.

All response delay time distribution fitting and model selection was performed in `R`.

### 2 Mechanistic model of the RNA life cycle with distributed delay times

##### 2.1 Mechanistic modelling of RNA life-cycle dynamics

We developed a mechanistic, ordinary differential equation (ODE)–based model to describe the dynamics of RNA life cycle stages for two genes, *ICAM1* and *TNFAIP3*, in a typical cell. The model explicitly represents transitions between RNA life cycle stages and degradation of RNA and protein molecules in selected stages. All transitions between life cycle stages, as well as degradation processes, were formulated using first-order mass-action kinetics.

In the model, expression starts with production of transcripts in an early splicing state. These can either go on to a late splicing state or directly be processed into mature mRNA. We mapped the early splicing state to the average number of cluster splicing spots (U1 and U2 combined) and late splicing to disperse spots. This was based on the parallel dynamics of U1 and U2 and our interpretation that disperse splicing transcripts will have left the transcription site and will, thus, be in a later stage of splicing. mRNA can be translated into protein or be degraded.

The model incorporates a negative feedback loop in which the *TNFAIP3* protein inhibits transcription of both genes. We used TNFAIP3 protein, A20 [3], as a proxy for several transcription dependent negative feedback regulators that inhibit both genes. For example NFKBIA is another negative feedback regulator and belongs to the same kinetic group of early response genes as TNFAIP [2]. The inhibitory effect is implemented as a hyperbolic function, characterised by an inhibition constant Ki. The hyperbolic function scales the transcription rate constant between the maximum value in the absence of the inhibiting protein, a half-maximal value determined by the estimated inhibition constant, and the asymptotically approached zero at high protein abundances. A graphical representation of the model is shown in Fig. 7 in the main manuscript.

An initial model pre-selection, comparing differnt model variants based on their BIC after model calibration, revealed that ICAM1 requires an additional step before early splicing, which is likely very fast for TNFAIP3 and thus not necessary for obtaining a good fit (Fig. S13a). Thus, we included the additional state ("nRNA") only for ICAM1. The model preselection also showed that the degradation rate of mRNA has to be gene-specific[2] and that an additional degradation step for transcripts in the late splicing state is not necessary.

The model was implemented and calibrated using a combination of computational tools following a workflow similar to that described by Villaverde *et al.* [8]. Model simulations were performed with AMICI [1], parameter estimation and model calibration were performed using pyPESTO [7], and the model was encoded in PEtab format [6]. Observables were defined using auxiliary ODEs to account for population-averaged measurements, incorporating single-cell response delays. Systematic model selection to determine gene-specific parameters was performed using PEtab Select [5]. Tellurium [4] was used for model construction and export to PEtab.

#### 2.2 Model structure

The *maximum model* for a typical cell comprises 10 state variables: 5 representing different RNA life cycle stages of *ICAM1*, 4 representing *TNFAIP3* RNA life cycle stages, and 1 for the *TNFAIP3* protein. There are 16 additional (auxiliary) state variables, 8 per gene, that are needed to implement the conditional weighting due to the distributed onset of transcription (see Section 2.5.3). The kinetic part of the model includes 16 rate constants and 2 inhibitory constants. The error model comprises 8 parameters, and the observable mapping requires 10 parameters: 8 for the exponential mixtures describing response delays and 2 for the scaling of molecule counts to spot counts for measurements of splicing cluster spots. Eight observables (four per gene) are compared to the population-average responder data during model calibration.

This model is referred to as the *maximum model*, since it contains gene-specific kinetic parameters for each life cycle stage. Parameters for the *TNFAIP3* part are defined as the corresponding *ICAM1* parameter multiplied by a parameter-specific scaling factor. Smaller model variants, in which parameters are shared between genes, can be encoded by fixing the respective scaling factor to 1.

Tables 1 and 2 provide an overview of all state variables and parameters. The model, encoded in Tellurium format, is provided in the Appendix.

#### 2.3 Observable Mapping

Model observables are quantities, calculated from the state variables, that correspond to the experimentally measured quantities. For example, an observable can be the sum of several state variables if the measurement detects all corresponding biological species (e.g., via an antibody), or a state variable multiplied by a scaling factor if the measurement is known to be proportional to a modeled quantity.

In our model, state variables represent RNA molecule counts, while the experimental data are reported in units of spot counts. For small spots, assumed to correspond to single RNA molecules, no

| state variable | description | gene | Fig. 7a |
|---|---|---|---|
| ICX0 | nascent RNA | *ICAM1* | nRNA |
| ICspliEarl | early splicing | *ICAM1* | esRNA |
| ICspliEarlY1 | early splicing (aux 1) | *ICAM1* | |
| ICspliEarlY2 | early splicing (aux 2) | *ICAM1* | |
| ICspliLate | late splicing | *ICAM1* | lsRNA |
| ICspliLateY1 | late splicing (aux 1) | *ICAM1* | |
| ICspliLateY2 | late splicing (aux 2) | *ICAM1* | |
| ICmrna | free mRNA | *ICAM1* | mRNA |
| ICmrnaY1 | free mRNA (aux 1) | *ICAM1* | |
| ICmrnaY2 | free mRNA (aux 2) | *ICAM1* | |
| ICtransl | translating mRNA | *ICAM1* | tlRNA |
| ICtranslY1 | translating mRNA (aux 1) | *ICAM1* | |
| ICtranslY2 | translating mRNA (aux 2) | *ICAM1* | |
| TNspliEarl | early splicing | *TNFAIP3* | esRNA |
| TNspliEarlY1 | early splicing (aux 1) | *TNFAIP3* | |
| TNspliEarlY2 | early splicingv (aux 2) | *TNFAIP3* | |
| TNspliLate | late splicing | *TNFAIP3* | lsRNA |
| TNspliLateY1 | late splicing (aux 1) | *TNFAIP3* | |
| TNspliLateY2 | late splicing (aux 2) | *TNFAIP3* | |
| TNmrna | free mRNA | *TNFAIP3* | mRNA |
| TNmrnaY1 | free mRNA (aux 1) | *TNFAIP3* | |
| TNmrnaY2 | free mRNA (aux 2) | *TNFAIP3* | |
| TNtransl | translating mRNA | *TNFAIP3* | tlRNA |
| TNtranslY1 | translating mRNA (aux 1) | *TNFAIP3* | |
| TNtranslY2 | translating mRNA (aux 2) | *TNFAIP3* | |
| TNprot | *TNFAIP3*protein | *TNFAIP3* | protein |

Table 1: Model state variables. Auxiliary (aux) state variables have no mechanistic meaning and are solely needed for the observable mapping accounting for distributed response delay times in the cell population.

| parameter | description | gene | data for estimation |
|---|---|---|---|
| k01 | kinetic | ICAM1 | responder population average count |
| k12_aux | factor scaling k12 | *TNFAIP3* | responder population average count |
| k12 | kinetic | ICAM1 | responder population average count |
| k13_aux | factor scaling k13 | TNFAIP3 | responder population average count |
| k13 | kinetic | ICAM1 | responder population average count |
| k23_aux | factor scaling k23 | TNFAIP3 | responder population average count |
| k23 | kinetic | ICAM1 | responder population average count |
| k3d_aux | factor scaling k3d | *TNFAIP3* | responder population average count |
| k3d | degradation | ICAM1 | responder population average count |
| k34_aux | factor scaling k34 | *TNFAIP3* | responder population average count |
| k34 | kinetic | *ICAM1* | responder population average count |
| k43_aux | factor scaling k43 | TNFAIP3 | responder population average count |
| k43 | kinetic | *ICAM1* | responder population average count |
| k5d | degradation | TNFAIP3 | responder population average count |
| Ki | inhibitory constant | *ICAM1* | responder population average count |
| Ki_aux | factor scaling Ki | *TNFAIP3* | responder population average count |
| Vmax | maximum transcription rate | ICAM | responder population average count |
| Vmax_aux | factor scaling Vmax | *TNFAIP3* | responder population average count |
| scale_ICAM1_SE | spots per molecules (cluster spots) | *ICAM1* | responder population average count |
| scale_TNFAIP3_SE | spots per molecules (cluster spots) | *TNFAIP3* | responder population average count |
| sd_ICAM1_M | error model | *ICAM1* | responder population average count |
| sd_ICAM1_SE | error model | *ICAM1* | responder population average count |
| sd_ICAM1_SL | error model | *ICAM1* | responder population average count |
| sd_ICAM1_TL | error model | *ICAM1* | responder population average count |
| sd_TNFAIP3_M | error model | *TNFAIP3* | responder population average count |
| sd_TNFAIP3_SE | error model | TNFAIP3 | responder population average count |
| sd_TNFAIP3_SL | error model | TNFAIP3 | responder population average count |
| sd_TNFAIP3_TL | error model | TNFAIP3 | responder population average count |
| IClambY1 | exponential rate 1 | ICAM1 | single cell response times (censored) |
| IClambY2 | exponential rate 2 | ICAM1 | single cell response times (censored) |
| ICw1 | mixture weight 1 | *ICAM1* | single cell response times (censored) |
| ICw2 | mixture weight 2 | *ICAM1* | single cell response times (censored) |
| TNlambY1 | exponential rate 1 | TNFAIP3 | single cell response times (censored) |
| TNlambY2 | exponential rate 2 | TNFAIP3 | single cell response times (censored) |
| TNw1 | mixture weight 1 | TNFAIP3 | single cell response times (censored) |
| TNw2 | mixture weight 2 | TNFAIP3 | single cell response times (censored) |

Table 2: Overview parameters as part of the maximum model.

scaling is necessary. For large spots, which may contain an unknown number of molecules, a scaling parameter converts the corresponding state variables from molecule counts to spot counts.

Importantly, the kinetic model describes the dynamics of a typical cell responding to TNF$\alpha$ stimulation without delay. In contrast, the data are averages over all responder cells at the time of measurement. Individual cells do not respond instantaneously, but rather with heterogeneous delays. Therefore, each data point reflects the population-average over many cells, each having expressed the gene for a different, only partially known duration. SN Figure 2.3 provides a conceptual illustration of this data-generating process.

While it is not possible to determine the exact expression time of a specific cell, the population-level distribution of response times is available from the delay distribution quantified in Section 1. To account for these distributed response delays in model calibration, we do not simulate individual cell trajectories. Instead, we compute population-level observables as the conditional expectation of the state variables given the response delay distribution, as described in the following subsections.

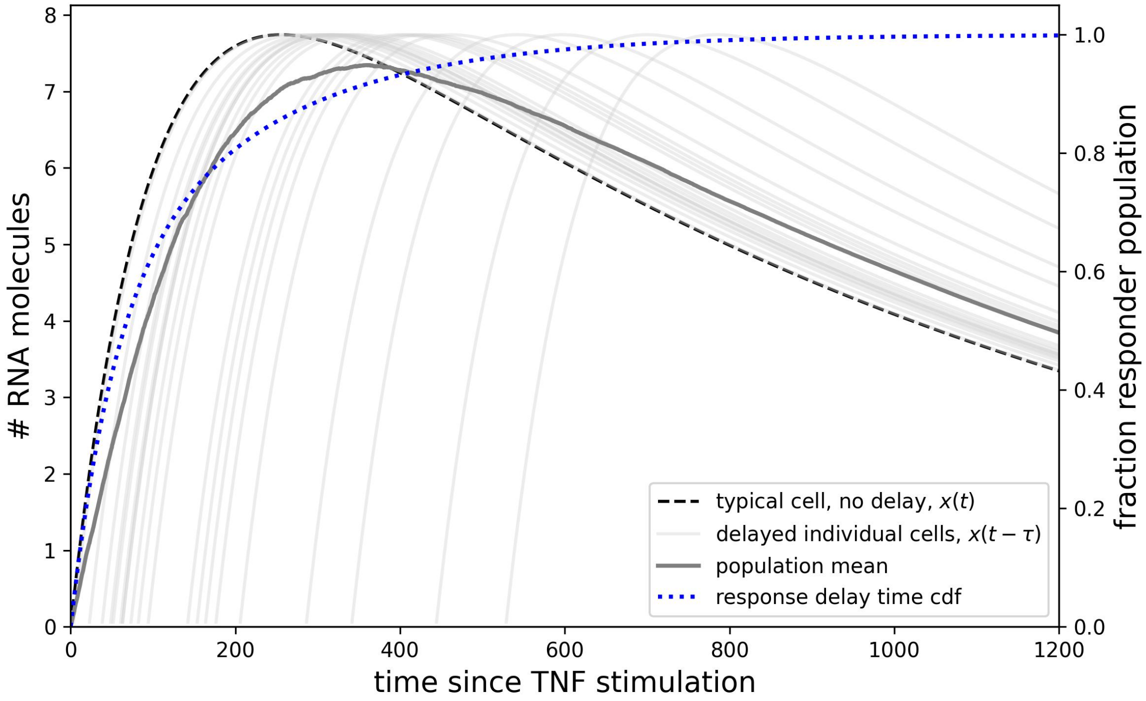

Conceptual illustration of the data-generating process. Individual cells respond to TNF$\alpha$ stimulation after a stochastic delay, but the underlying kinetic trajectory of each cell is assumed to be identical. At the time of measurement, cells are therefore at different points along this trajectory. The population-average observable corresponds to an average over cells with heterogeneous response "ages," as would be obtained by simulating multiple trajectories each starting after a cell-specific delay and averaging their values at the measurement time. The cdf of the response delay time distribution used in this illustration is provided for reference. Note: This SN Figure is purely illustrative; the model implementation does not simulate individual cell trajectories.

#### 2.4 Conditional expectation

To map model state variables $x_i(t)$ to population-level observables $y_i(t)$ , we account for asynchronous cellular responses on the population level, which we quantified by a two-component exponential mixture distribution:

$$p(\tau) = w_1 \lambda_1 e^{-\lambda_1 \tau} + w_2 \lambda_2 e^{-\lambda_2 \tau}, \tag{10}$$

where $\tau$ is the response delay time, $w_1, w_2$ are the mixture weights ($w_1 + w_2 = 1$), and $\lambda_1, \lambda_2$ are the decay rates of the two exponential mixture components.

The experimentally measured quantity corresponds to the average spot count among cells that have already responded to TNF$\alpha$ stimulation at experimental time $t$ (responders). Following induction, all cells are assumed to follow identical deterministic single-cell dynamics $x(t)$, where time is measured since induction. The observable is therefore the conditional expectation of $x_i(t-\tau)$ given the response delay time distribution and conditional on $\tau \leq t$:

$$y_i(t) = \mathbb{E}[x_i(t-\tau) \mid \tau \leq t] = \frac{\int_0^t x_i(t-\tau)\, p(\tau)\, d\tau}{F(t)}, \tag{11}$$

with

$$F(t) = \Pr(\tau \leq t) = \int_0^t p(\tau)\, d\tau. \tag{12}$$

The numerator represents a time-weighted population average, while the denominator corresponds to the probability that a cell has responded at time $t$ (or the fraction of cells that have responded by time $t$), implementing the conditioning.

In the following we explain how this conditional expectation is computed within a standard workflow of calibrating ODE models instead of using averaged of Monte Carlo simulations of single cells.

#### 2.5 From conditional expectation integrals to auxiliary ODEs

The derivation starts from the conditional time-weighted population average over responder cells as given in equation (11). The numerator of this expression is convolution in delay time space, which—after a change of variables to time-since-induction and assuming exponential delay distributions—admits an exact representation in form of additional ODEs, sometimes called leaky integrator ODEs.

##### 2.5.1 Convolution Formulation

The numerator of the conditional expectation is a convolution of the single-cell trajectory $x_i(t)$ with the delay distribution $p(\tau)$:

$$\int_0^t x_i(t-\tau)\, p(\tau)\, d\tau. \tag{13}$$

This convolution accounts for heterogeneous response times across the cell population.

##### 2.5.2 Change of Variables

To obtain a formulation amenable to an ODE-based representation, a change of variables is introduced by defining $s = t - \tau$, corresponding to time-since-induction (or expression age). Under this transformation, the numerator becomes

$$\int_0^t x_i(s)\, p(t-s)\, ds. \tag{14}$$

The delay distribution was modeled as a mixture of exponential distributions,

$$p(\tau) = \sum_{k=1}^{2} w_k \lambda_k e^{-\lambda_k \tau}, \tag{15}$$

which implies

$$p(t-s) = \sum_{k=1}^{2} w_k \lambda_k e^{-\lambda_k (t-s)}. \tag{16}$$

Substituting this expression into the convolution yields

$$\int_0^t x_i(s)\, p(t-s)\, ds = \sum_{k=1}^{2} w_k \int_0^t x_i(s)\, \lambda_k e^{-\lambda_k (t-s)} ds. \tag{17}$$

###### 2.5.3 Auxiliary ODEs

Each of the 2 integrals on the right hand side of equation (17) are of the form

$$z_k(t) = \int_0^t x_i(s)\,\lambda_k e^{-\lambda_k(t-s)} ds. \tag{18}$$

Differentiation with respect to time yields a differential equation for $z_k(t)$:

$$\frac{dz_k(t)}{dt} = \lambda_k\left(x_i(t) - z_k(t)\right). \tag{19}$$

which is known as a leaky integrator ODE.

This means that by introducing an additional (auxiliary) state variable $z_k(t)$ with dynamics defined by the ODE in equation (19), the integral in equation (18) is solved by numerical integration whenever the model is simulated. Since equation (17) has 2 of these integrals, we have to introduce 2 additional state variables for each of the state variables $x_i$, that corresponds to a measured quantity. The ODEs of these state variables then read

$$\frac{dz_{i,1}(t)}{dt} = \lambda_1\left(x_i(t) - z_{i,1}(t)\right), \tag{20}$$

$$\frac{dz_{i,2}(t)}{dt} = \lambda_2\left(x_i(t) - z_{i,2}(t)\right) \tag{21}$$

where $\lambda_1, \lambda_2$ are the decay rates of the two components of the exponential mixture as defined in equation (10). The ODEs provide an exact representation of the unnormalized convolution terms.

###### 2.5.4 Conditioning on fraction of cells that have already responded ($\tau \leq t$)

To obtain to The fraction of cells that have responded by time $t$ is given by the cumulative distribution function

$$F(t) = \int_0^t p(\tau)\,d\tau = \sum_{k=1}^{2} w_k\left(1 - e^{-\lambda_k t}\right). \tag{22}$$

The conditional population-average observable is therefore obtained as

$$y_i(t) = \frac{\sum_{k=1}^{2} w_k z_{i,k}(t)}{\sum_{k=1}^{2} w_k\left(1 - e^{-\lambda_k t}\right)}. \tag{23}$$

where, $w_1, w_2$ are the mixture weights and $\lambda_1, \lambda_2$ the exponential rates as defined in equation (10). This expression defines the observable mapping as it is implemented in the PeTab file that specifies the observables.

###### 2.5.5 Summary

Conceptually the mapping of the state variables to the model observables proceeds like this:

1. Simulate single-cell trajectory $x_i(t)$ assuming instantaneous response.
2. Simultaneously simulate the auxiliary ODEs $z_{i,1}(t), z_{i,2}(t)$ for each state variable that has a corresponding measurement.
3. Compute a weighted sum of the auxiliary state variables $z_{i,1}(t), z_{i,2}(t)$ using mixture weights $w_1, w_2$.
4. Normalize the weighted sum by the cumulative fraction of responder cells $F(t)$ to obtain the population-level observable $y_i(t)$.
5. For splicing cluster spot counts apply multiplicative scaling parameter to map molecule counts to experimental spot counts $y_i(t)$.

This approach provides a fully ODE-based mapping from typical single-cell dynamics to population-average observables while explicitly accounting for heterogeneous response timing.

#### 2.6 Model calibration

Model calibration was performed by simultaneously estimating kinetic, scaling, and error model parameters from the available average spot count data of the responder population. The final measurement time point (1600 minutes) was excluded from parameter estimation, since the late-time dynamics reflect processes not captured by the model. All data points were treated as independent observations at the population level.

Parameter estimation was carried out using the Python package pyPESTO [7], with the Python package AMICI [1] used for numerical simulation of the ODE model. The parameter estimation problem was specified in the PEtab standard format [6]. Parameters were estimated in logarithmic space, with individually specified search boundaries for each parameter. The bounds for the splicing cluster scaling parameters were informed by empirical measurements comparing fluorescence intensities of cluster spots to those of single-molecule spots, which resulted in intensity ratios between 1.2 and 12.8. As the scaling parameters map molecule counts to spot counts (spots per molecule), the reciprocal of this empirically observed range was used to define the admissible parameter bounds.

A multi-start local optimization strategy was employed to reduce the risk of convergence to local optima. The error model assumed additive Gaussian noise in the model observables.

#### 2.7 Quantification of parameter uncertainty

For the model selected by the automated model selection procedure, uncertainty in the maximum likelihood parameter estimates was assessed using a profile likelihood approach as implemented in pyPESTO [7], providing confidence intervals and identifiability assessment for individual parameters.

During multi-start optimization, strong parameter correlations between the rate constants $k_{34}$ and $k_{43}$ were observed, indicating that the available data primarily constrain their ratio rather than the individual parameter values. To improve parameter identifiability and optimizer convergence, the model was therefore reparameterized by replacing the parameter $k_{43}$ with the product $k_{34} \cdot K_D$, where

$$K_D = \frac{k_{43}}{k_{34}}. \tag{24}$$

Parameter confidence intervals where assessed based on this model. Parameter boundaries for profiling where set to the Maximum likelihood estimate (MLE) plus/minus 3 on the log10 scale, with the exception of the 2 scale_TNFAIP3_SE and scale_ICAM1_SE, for which the upper and lower bound where kept the same as for parameter estimation, since the boundaries were derived from measurements of spot intensity.

SN Figure 2.7 shows the 90% and 95% confidence intervals resulting from the profiling of each parameter. The model is not finitely identified to the confidence level 95%, given the experimental data. The parameters Ki, k34, k3d, k3d_aux, scale_TNFAIP3_SE and scale_ICAM1_SE are not finitely identified to the confidence level 95%. Ki_aux (corresponding to Ki of *ICAM1* over Ki of *TNFAIP3*), is finitely identified to the confidence level 95%, supporting inference of relative differences in gene-specific inhibition.

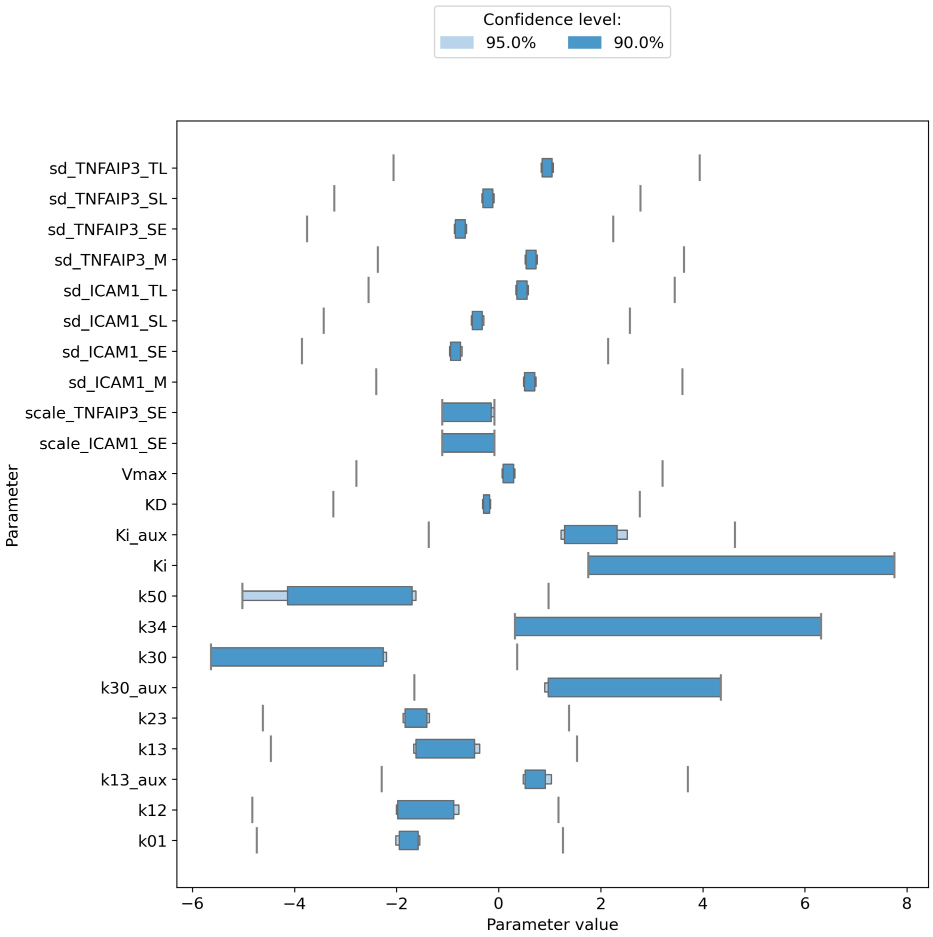

Vertical grey line indicate the boundaries the parameter was allowed to vary in, which were set to the maximum likelihood estimate plus/minus 3 on the log10 scale for all parameters except the scale$_parameters.Pleasenotethatk30correspondstok3$[illegible]

#### 2.8 Model selection

Alternative model hypotheses differing in model structure, network topology, and parameter sharing between the two genes were assessed using a systematic model-comparison approach. Model variants differed in the inclusion or exclusion of specific state variables or life-cycle stage transitions, as well as in whether individual kinetic parameters were shared between genes or allowed to be gene-specific. For all model variants, parameters were estimated by maximum likelihood using the calibration procedure described above.

Model comparison was performed using the Bayesian Information Criterion (BIC), defined in terms of the maximized log-likelihood as

$$\mathrm{BIC} = 2\,\ell_{\min} + k \log N, \tag{25}$$

where $\ell_{\min}$ denotes the minimum negative log-likelihood achieved by the model, $k$ is the number of free parameters, and $N$ is the number of data points. The BIC balances model fit and model complexity by penalizing the number of estimated parameters, thereby favoring parsimonious models that adequately explain the data. Competing model hypotheses were ranked according to their BIC values, with lower BIC indicating stronger support by the data.

Two complementary model-selection strategies were employed. First, a set of biologically motivated model variants was constructed manually to test alternative hypotheses regarding model topology and included mechanisms. These manually defined variants were calibrated independently and compared based on their BIC values.

Second, systematic model selection with respect to gene-specific parameters was performed using PEtab Select [5]. PEtab Select constructs a hierarchy of nested model variants in which selected parameters are either shared between genes or allowed to be gene-specific. Starting from the smallest model in terms of the number of free parameters, a forward-selection strategy was applied, iteratively introducing additional gene-specific parameters when supported by an improvement in BIC. This procedure enables the identification of the minimal set of parameters that must differ between genes to explain the observed differences in RNA dynamics.

Using the maximum model as a base model, we performed systematic model discrimination to identify the optimal combination of free and shared parameters. Using the Bayesian Information Criterion (BIC), we found positive evidence ($\Delta$ BIC 2.3 to the next best model 36) that one model produces a better fit of the data while limiting the number of free parameters (Fig. 7b). In this model direct mRNA production from early splicing with a characteristic time $\tau_{13}$ and the negative feedback with the inhibitory constant Ki had gene specific values while the other parameters were shared between the genes. Taking into account the mRNA degradation ($\tau_{3d}$) and the initial delay step ($\tau_{01}$), the best model had four gene-specific parameters and was able to produce a fit that captured the dynamics of all observables (Fig. 7c, Fig. S13b). The confidence interval (CI) for the ratio of the TNFAIP3- and ICAM1-specific parameter values did not include 1 (Fig. 7d) indicating that these expression steps, indeed, need to happen with different rates. Early splicing of ICAM1 required an average time of 29 min (3-51 min, 95% CI), but was five times faster (3.3-11.1-fold, 95% CI) for TNFAIP3. ICAM1 transcripts were quite long-lived with an average lifetime of 434min ($\geq$142 min, 95% CI), whereas TNFAIP3 lifetime was 25-fold shorter ($\geq$ 10-fold, 95% CI). Finally, ICAM1 was much more sensitive to the negative feedback, with a 54-fold (19-459-fold, 95% CI) lower inhibitory constant.

### 3 Monte Carlo simulation of Actinomycin D treatment 60 minutes after TNF$\alpha$ stimulation

To simulate a scenario in which cells are treated with Actinomycin D at a time 60 minutes after stimulation with TNF$\alpha$, we performed a Monte Carlo simulation of a population of cells and calculated the average of the whole cell population by averaging the single cell trajectories.

Treatment with Actinomycin D inhibits gene transcription, which in the model corresponds to a decrease in the parameters $k_{01}$ and $V_{\max}$. In addition, we assume that after Actinomycin D treatment no additional cells respond to TNF$\alpha$ stimulation. Consequently, the fraction of responding cells does not increase further after Actinomycin D treatment. This constraint is not easily implemented within the framework of auxiliary ODEs and conditional expectations (Section 2), which assumes that cells respond according to the response delay time distribution. Therefore, we adopted a Monte Carlo approach. The simulations were implemented in R with the RxODE package.

Individual cells were simulated using the model structure selected in the model selection procedure, minus the auxiliary state variables, with parameters calibrated to the average mRNA counts of the responder population as described in Section 2. To allow transcription of each gene to start at different times after TNF$\alpha$ stimulation, separate parameters were introduced for $V_{\max}$ in the ODEs describing *ICAM1* and *TNFAIP3* transcription.

To account for the distributed response times observed across individual cells, a response delay time was sampled for each cell and gene (*ICAM1* and *TNFAIP3*) from the corresponding response delay time distributions. Simulations of the individual cells were carried out in several parts, corresponding to different parameter values, with each part using the last value of the the previous part as the initial condition for the simulation. The different simulation parts were joined to retrieve the full time course trajectory of a single cell. Simulations for each cell were initialized at clock time 0 without transcription in either gene ($V_{\max,\mathrm{IC}} = V_{\max,\mathrm{TN}} = 0$). At the sampled response delay time, the corresponding $V_{\max}$ parameter was set to its maximum likelihood estimate and the simulation was continued to 60 minutes, the time of Actinomycin D treatment.

To simulate Actinomycin D treatment at 60 minutes, the parameters $V_{\max,\mathrm{TN}}$, $V_{\max,\mathrm{IC}}$, and $k_{01}$ were gradually reduced from 100% of their original value to 10% within 15 minutes after treatment (from 65 minutes: 70%, from 70 minutes: 40% , from 75 minutes: 10%). Simulations were then continued until 120 minutes of clock time after TNF$\alpha$ stimulation. Cells that were nonresponders in both genes (both response delay times $\leq$ 60 minutes) were added to the population with zero trajectories for all state variables. The trajectories of all simulated cells were averaged to obtain the whole population average, which also includes nonresponders in one or both genes.

Comparing the whole population average predicted based on the conditional expectation population model approach with the average based on Monte Carlo single cell simulations (SN Figure 3), shows that the conditional expectation model implementation slightly overestimates the average of *ICAM1* state variables. This is caused by the negative regulation of *ICAM1* transcription by the *TNFAIP3* protein, that will lead to different trajectories for the *ICAM1* state variables depending on the time lag between the induction of *TNFAIP3* and *ICAM1* when simulating individual cells. The conditional expectation model implementation corresponds identical trajectories for each cell, just shifted according to the delay time distribution. This is fulfilled by the *TNFAIP3* state variable trajectories, but not by the *ICAM1* state variable trajectories due to the inhibitory interaction between *TNFAIP3* and *ICAM1*.

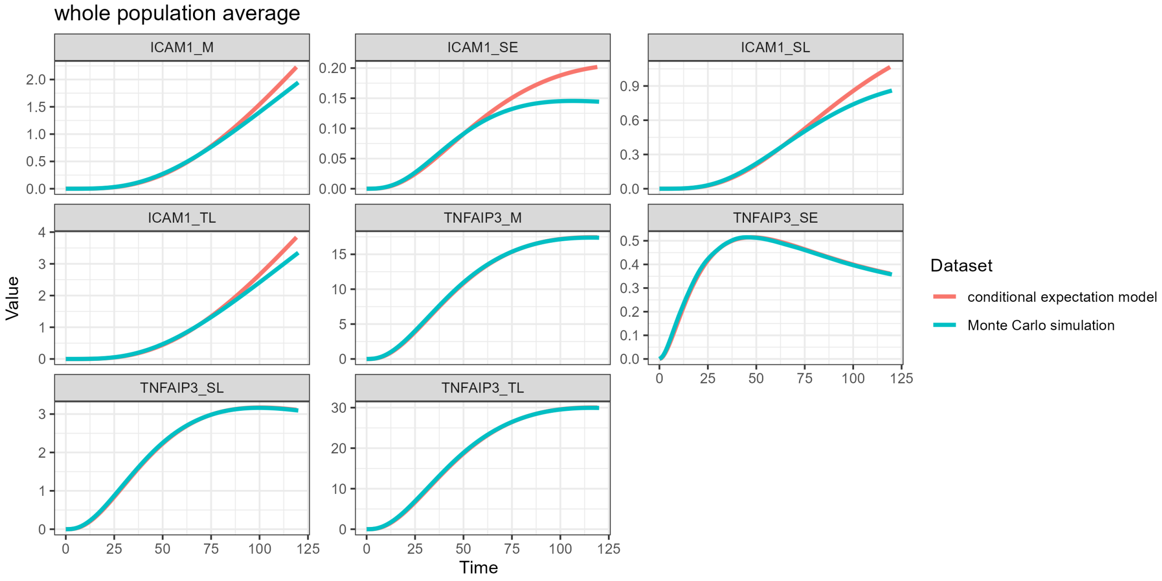

Comparison of whole population average trajectories between conditional expectation model and Monte Carlo simulation

### 4 Appendix

#### 4.1 Specification of maximum model in tellurium format

Please note that the model specified here does not include the actual observable mapping, which was implemented in the dedicated table in the PEtab standard we used to specify the parameter estimation problem.

```
# import library
import tellurium as te

# specify model
model_selection = te.loada('''
model ModelNorm_X0_mRNA_no20_select

  // Species:
  species ICX0,  ICspliEarl, ICspliEarlY1, ICspliEarlY2, ICspliLate, ICspliLateY1,
      ICspliLateY2, ICmrna, ICmrnaY1, ICmrnaY2, ICtransl, ICtranslY1, ICtranslY2,
      TNspliEarl, TNspliEarlY1, TNspliEarlY2, TNspliLate, TNspliLateY1, TNspliLateY2,
      TNmrna, TNmrnaY1, TNmrnaY2, TNtransl, TNtranslY1, TNtranslY2 , TNprot;

  // Reactions:
  ICX0R : => ICX0; (Ki/(TNprot + Ki)) * Vmax;
  ICtranscription:  ICX0 => ICspliEarl; ICX0 * k01  ;
  ICsplicing: ICspliEarl  => ICspliLate;  k12  *ICspliEarl;
  ICendsplice:  ICspliEarl => ICmrna; k13  * ICspliEarl;
  ICposttrans: ICspliLate => ICmrna; k23  * ICspliLate;
  ICdegmrna: ICmrna => ; k3d  * ICmrna;
  ICtranslat: ICmrna => ICtransl; k34  * ICmrna;
  ICrelease: ICtransl => ICmrna; k43  * ICtransl ;
```

```
TNtranscription:  => TNspliEarl; (Ki * Ki_aux /(TNprot + Ki * Ki_aux)) * Vmax *
    Vmax_aux  ;
TNsplicing: TNspliEarl  => TNspliLate; k12 *k12_aux *TNspliEarl;
TNendsplice:  TNspliEarl => TNmrna; k13 * k13_aux * TNspliEarl;
TNposttrans: TNspliLate => TNmrna; k23 * k23_aux * TNspliLate;
TNdegmrna: TNmrna => ; k3d *k3d_aux * TNmrna;
TNtranslat: TNmrna => TNtransl; k34 *k34_aux   * TNmrna;
TNrelease: TNtransl => TNmrna + TNprot; k43 *k43_aux  * TNtransl ;

prot_deg: TNprot => ; k5d *TNprot;

// auxilliary ODEs for time weighted population means
// ICAM
ICspliEarlY1' = IClambY1 *(ICspliEarl - ICspliEarlY1);
ICspliEarlY2' = IClambY2 *(ICspliEarl - ICspliEarlY2);
ICspliLateY1' = IClambY1 *(ICspliLate - ICspliLateY1);
ICspliLateY2'= IClambY2 *(ICspliLate -  ICspliLateY2);
ICmrnaY1' = IClambY1 *( ICmrna - ICmrnaY1);
ICmrnaY2'   = IClambY2 *( ICmrna - ICmrnaY2)   ;
ICtranslY1' = IClambY1 *( ICtransl - ICtranslY1);
ICtranslY2' = IClambY2 *( ICtransl - ICtranslY2);

//\textit{TNFAIP3}
TNspliEarlY1' =  TNlambY1 * ( TNspliEarl - TNspliEarlY1);
TNspliEarlY2' =  TNlambY2 * ( TNspliEarl - TNspliEarlY2);
TNspliLateY1' =  TNlambY1 * ( TNspliLate - TNspliLateY1);
TNspliLateY2'=  TNlambY2 * (TNspliLate -  TNspliLateY2);
TNmrnaY1' =  TNlambY1 * (TNmrna -  TNmrnaY1);
TNmrnaY2'   =  TNlambY2 * (  TNmrna -  TNmrnaY2)  ;
TNtranslY1' =  TNlambY1 * (  TNtransl -  TNtranslY1);
TNtranslY2' =  TNlambY2 * (  TNtransl -  TNtranslY2);

// Species initializations:

ICX0 = 0;
ICspliEarl = 0;
ICspliEarlY1 = 0;
ICspliEarlY2 = 0;
ICspliLate = 0;
ICspliLateY1= 0;
ICspliLateY2= 0;
ICmrna= 0;
ICmrnaY1= 0;
ICmrnaY2= 0;
ICtransl= 0;
ICtranslY1= 0;
ICtranslY2= 0;

TNspliEarl= 0;
TNspliEarlY1= 0;
TNspliEarlY2= 0;
TNspliLate= 0;
TNspliLateY1= 0;
TNspliLateY2= 0;
TNmrna= 0;
TNmrnaY1= 0;
TNmrnaY2= 0;
TNtransl= 0;
TNtranslY1= 0;
TNtranslY2= 0;
TNprot= 0;

// Variable initializations

IClambY1 = 0.5;
IClambY2  = 0.5;
TNlambY1  = 0.5;
TNlambY2  = 0.5;
```

```
  k01  = 1;
  Ki   = 1;
  Vmax   = 1 ;
  k12   = 1 ;
  k13   = 1 ;
  k23   = 1;
  k3d   =  1 ;
  k34   = 1;
  k43   = 1;

  Ki_aux    = 1;
  Vmax_aux    =  1 ;
  k12_aux    = 1;
  k13_aux    = 1 ;
  k23_aux    = 1;
  k3d_aux    =   1 ;
  k34_aux   = 1;
  k43_aux   = 1;

  k5d   = 1;

end
''')
```

#### 4.2 Selected final model reparametrized in terms of KD

Version of the model containing only the parameters selected by automated model selection, and reparametrized by replacing parameter k43 with k34 * KD, where KD = k43/k34. All parameters that were ruled out by the automated model selection have been removed from the model definition manually.

```
# sepcify model
M_0010001_KD = te.loada('''
model M_0010001_KD

  // Species:
  species ICX0,  ICspliEarl, ICspliEarlY1, ICspliEarlY2, ICspliLate, ICspliLateY1,
      ICspliLateY2, ICmrna, ICmrnaY1, ICmrnaY2, ICtransl, ICtranslY1, ICtranslY2,
      TNspliEarl, TNspliEarlY1, TNspliEarlY2, TNspliLate, TNspliLateY1, TNspliLateY2,
      TNmrna, TNmrnaY1, TNmrnaY2, TNtransl, TNtranslY1, TNtranslY2 , TNprot;

  // Reactions:
  ICX0R : => ICX0; (Ki/(TNprot + Ki)) * Vmax;

  ICtranscription:  ICX0 => ICspliEarl; ICX0 * k01  ;
  ICsplicing: ICspliEarl  => ICspliLate;  k12  *ICspliEarl;
  ICendsplice:  ICspliEarl => ICmrna; k13  * ICspliEarl;
  ICposttrans: ICspliLate => ICmrna; k23  * ICspliLate;
  ICdegmrna: ICmrna => ; k3d  * ICmrna;
  ICtranslat: ICmrna => ICtransl; k34  * ICmrna;
  ICrelease: ICtransl => ICmrna; k34*KD  * ICtransl ;

  TNtranscription:  => TNspliEarl; (Ki * Ki_aux  /(TNprot + Ki * Ki_aux  )) * Vmax   ;
  TNsplicing: TNspliEarl  => TNspliLate; k12  *TNspliEarl;
  TNendsplice:  TNspliEarl => TNmrna; k13  *k13_aux  * TNspliEarl;
  TNposttrans: TNspliLate => TNmrna; k23   * TNspliLate;
  TNdegmrna: TNmrna => ; k3d  *k3d_aux  * TNmrna;
  TNtranslat: TNmrna => TNtransl; k34   * TNmrna;
  TNrelease: TNtransl => TNmrna + TNprot; k34*KD  * TNtransl ;

  prot_deg: TNprot => ; k5d *TNprot;

  // auxilliary ODEs for weighted population means
  // ICAM

  ICspliEarlY1' = IClambY1 *(ICspliEarl - ICspliEarlY1);
```

```
  ICspliEarlY2' = IClambY2 *(ICspliEarl - ICspliEarlY2);
  ICspliLateY1' = IClambY1 *(ICspliLate - ICspliLateY1);
  ICspliLateY2'= IClambY2 *(ICspliLate -  ICspliLateY2);
  ICmrnaY1' = IClambY1 *( ICmrna - ICmrnaY1);
  ICmrnaY2'   = IClambY2 *( ICmrna - ICmrnaY2)   ;
  ICtranslY1' = IClambY1 *( ICtransl - ICtranslY1);
  ICtranslY2' = IClambY2 *( ICtransl - ICtranslY2);

  // TNFAIP3

  TNspliEarlY1' =  TNlambY1 * ( TNspliEarl - TNspliEarlY1);
  TNspliEarlY2' =  TNlambY2 * ( TNspliEarl - TNspliEarlY2);
  TNspliLateY1' =  TNlambY1 * ( TNspliLate - TNspliLateY1);
  TNspliLateY2'=  TNlambY2 * (TNspliLate -  TNspliLateY2);
  TNmrnaY1' =  TNlambY1 * (TNmrna -  TNmrnaY1);
  TNmrnaY2'   =  TNlambY2 * (  TNmrna -  TNmrnaY2)  ;
  TNtranslY1' =  TNlambY1 * (  TNtransl -  TNtranslY1);
  TNtranslY2' =  TNlambY2 * (  TNtransl -  TNtranslY2);

  // Species initializations:

  ICX0 = 0;
  ICspliEarl = 0;
  ICspliLate = 0;
  ICmrna= 0;
  ICtransl= 0;

  TNspliEarl= 0;
  TNspliLate= 0;
  TNmrna= 0;
  TNtransl= 0;

  TNprot= 0;

  // Variable initializations:

  k01  = 1;
  Ki   = 1;
  Vmax   = 1 ;
  k12   = 1 ;
  k13   = 1 ;
  k23   = 1;
  k3d   =  1 ;
  k34   = 1;
  KD =1;
  Ki_aux  = 1;
  k13_aux  = 1 ;
  k3d_aux  =   1 ;
  k5d   =1;

  IClambY1 = 0.5;
  IClambY2  = 0.5;
  TNlambY1  = 0.5;
  TNlambY2  = 0.5;

end
''')
```

##### 4.3 Model for Monte Carlo simulation of Actinomycin D treatment

Model as implemented in RxODE for use in Monte Carlo simulations.

```
mod3 <- function() {
  ini({
    k01 = 0.018286057
    k12 = 0.014886163
    k13_aux =   5.13613196667099
    k13 = 0.034484845
```

```
    k23 =        0.023952131
    k30_aux =    22.48448588
    k30 =        0.00230178
    k34 = 2092.128474
    k50 = 0.009518319
    Ki  = 56692.21092
    Ki_aux=      42.52016172
    KD  = 0.580717241

    # introduced parameters to be able to switch them on individually
    Vmax_IC = 1.625136291
    Vmax_TN = 1.625136291
  })
  model({
    ICX0(0) = 0
    ICspliEarl(0) = 0
    ICspliLate(0)= 0
    ICmrna(0) = 0
    ICtransl(0) = 0
    TNspliEarl(0) = 0
    TNspliLate(0) = 0
    TNmrna(0) = 0
    TNtransl(0) = 0
    TNprot(0) = 0

    # ODES describing a typical cell
    d/dt(ICX0) = Ki/(TNprot+Ki)*Vmax_IC - ICX0*k01
    d/dt(ICspliEarl) = ICX0*k01 - k12*ICspliEarl - k13*ICspliEarl
    d/dt(ICspliLate) = k12*ICspliEarl - k23*ICspliLate
    d/dt(ICmrna) = k13*ICspliEarl + k23*ICspliLate - k30*ICmrna- k34*ICmrna + k34*KD*
        ICtransl
    d/dt(ICtransl) = k34*ICmrna - k34*KD*ICtransl
    d/dt(TNspliEarl) = Ki*Ki_aux/(TNprot+Ki*Ki_aux)*Vmax_TN - k12*TNspliEarl - k13*
        k13_aux*TNspliEarl
    d/dt(TNspliLate) = k12*TNspliEarl - k23*TNspliLate
    d/dt(TNmrna) = k13*k13_aux*TNspliEarl + k23*TNspliLate - k30*k30_aux*TNmrna - k34*
        TNmrna + k34*KD*TNtransl
    d/dt(TNtransl) = k34*TNmrna - k34*KD*TNtransl
    d/dt(TNprot) = k34*KD*TNtransl - k50*TNprot

  })
}
```
